# DNA origami vaccines program antigen-focused germinal centers

**DOI:** 10.1101/2025.02.21.639354

**Authors:** Anna Romanov, Grant A. Knappe, Larance Ronsard, Heikyung Suh, Marjan Omer, Asheley P. Chapman, Vanessa R. Lewis, Katie Spivakovsky, Josue Canales, Boris Reizis, Ryan D. Tingle, Christopher A. Cottrell, Torben Schiffner, Daniel Lingwood, Mark Bathe, Darrell J. Irvine

## Abstract

Recruitment and expansion of rare precursor B cells in germinal centers (GCs) is a central goal of vaccination to generate broadly neutralizing antibodies (bnAbs) against challenging pathogens such as HIV. Multivalent immunogen display is a well-established method to enhance vaccine-induced B cell responses, typically accomplished by using natural or engineered protein scaffolds. However, these scaffolds themselves are targets of antibody responses, with the potential to generate competitor scaffold-specific B cells that could theoretically limit expansion and maturation of “on-target” B cells in the GC response. Here, we rationally designed T-independent, DNA-origami based virus-like particles (VLPs) with optimal antigenic display of the germline targeting HIV Env immunogen, eOD-GT8, and appropriate T cell help to achieve a potent GC response. In preclinical mouse models, these DNA-VLPs expanded significantly higher frequencies of epitope-specific GC B cells compared with a state-of-the-art clinical protein nanoparticle. Optimized DNA-VLPs primed germinal centers focused on the target antigen and rapidly expanded subdominant broadly neutralizing antibody precursor B cells for HIV with a single immunization. Thus, avoiding scaffold-specific responses augments priming of bnAb precursor B cells, and DNA-VLPs are a promising platform for promoting B cell responses towards challenging subdominant epitopes.

## Main

Generation of broadly neutralizing antibodies (bnAbs) that recognize diverse viral variants is a central goal for successful vaccination against HIV^1^ and universal vaccines for other pathogens such as influenza^2, 3^, coronaviruses^4^, and dengue virus^5^. Precursor B cells capable of evolving to produce bnAbs are characteristically rare in the naive repertoire and often have low affinity for target immunogens^6^. Engaging rare B cell clones that belong to immunologically subdominant bnAb lineages to promote their somatic hypermutation and affinity maturation in germinal centers (GCs) remains an outstanding challenge and the primary aim of rational vaccine design^1, 7-10^.

One powerful approach to augment humoral responses to vaccine antigens is via multivalent display of many copies of an antigen on the surface of protein nanoparticles^11-19^. Nanoparticulate antigens exhibit improved lymph trafficking^18, 20, 21^ and enhance B cell receptor (BCR) crosslinking^22, 23^, inducing signal amplification downstream of the BCR^24^ and enabling potent valency-dependent B cell activation across a broad range of BCR affinities^25^. However, multimerization of antigens on protein scaffolds may be a double-edged sword. When protein scaffolds are used to display target antigens, they act as thymus-dependent (TD) repetitively arrayed antigens themselves, eliciting priming of scaffold-specific B cells towards irrelevant protein substrates that potentially compete in GCs against the desired, epitope-specific bnAb precursor B cells^17, 26-30^.

Scaffold-specific antibody responses against bacterially-derived scaffolds such as ferritin^17, 27, 28^ and lumazine synthase^29, 30^, and computationally-designed two-component nanoparticles^26^ have been quantified via serology and neutralization assays in animal models and humans. However, it remains poorly understood how these scaffold-specific competitor B cells shape the clonal competition dynamics within GCs, which rely on finite populations of B cells and helper T cells. While strategies to limit scaffold-specific responses have been used, such as glycosylation^30-32^ or physical masking of exposed scaffold epitopes^26, 33^, they have generally failed to fully eliminate off-target responses.

Importantly, the presence of competitive scaffold-specific B cells could lead to several outcomes that remain unclear due to the lack of suitable tools to test these hypotheses: 1) these B cells may outcompete antigen-specific GC B cells for limited T cell help and antigen availability^9^, particularly if the scaffold is more immunogenic than the antigen itself^34^; 2) they might contribute to immunological imprinting or ‘original antigenic sin’ upon repeated dosing or repurposing of the nanoparticle platform^35, 36^, and 3) they could lead to high titers of anti-scaffold antibodies that facilitate epitope masking and antibody feedback that alters the subsequent immune response^37^.

Each of these competitive effects are of particular concern when designing a vaccine against immunologically subdominant epitopes, such as the CD4 binding site (CD4bs) in HIV Env antigens, which is the target of VRC01-class bnAbs^26^. This task is further complicated by the extremely low frequency of bnAb precursor B cells in the naive B cell repertoire^6, 34, 38^.

One strategy to prime these rare subdominant precursors is immunization with engineered germline targeting immunogens, such as the engineered outer domain of gp120, eOD-GT8, which was designed to bind germline reverted VRC01-class BCRs with high affinity^39^. A protein nanoparticle form of eOD-GT8 (eOD-GT8 60mer) was developed by fusing eOD-GT8 with the self-assembling bacterial protein lumazine synthase (LumSyn); 60 copies of LumSyn self-assemble to form an icosahedral nanoparticle displaying eOD-GT8 at high density on the particle surface^38, 40^. This ∼30 nm protein nanoparticle has been shown to effectively activate VRC01-class B cell precursors in humanized mouse models and is currently undergoing clinical trials as a germline targeting immunogen for VRC01-class bnAbs against HIV^29, 41, 42^. While eOD-GT8 60mer has been shown to successfully activate VRC01-class precursor B cells in humans^29^, it also elicits a strong LumSyn-specific antibody response^29, 30^, and it remains unknown whether, and how, this scaffold-specific response may limit the priming of the “on-target” bnAb precursors of interest. Because germline targeting immunizations are just the first step toward shepherding B cells to become bnAb producers, it is critical that precursor activation and expansion be as efficient as possible^1, 43^.

As an alternative to protein-based scaffolds, we previously demonstrated that virus-like particles (VLPs) formed by the programmed assembly of thymus-independent (TI) DNA origami (DNA-VLPs) enable precise antigen display^44^ and enhance antigen-specific antibody titers while avoiding anti-scaffold B cell responses, as there is no thymus-dependent (protein) substrate except for the antigen^28^. Furthermore, DNA origami has been shown to be minimally immune-stimulatory^45, 46^, unless specific CpG motifs are included to activate innate Toll-like receptor pathways^47, 48^. Thus, DNA origami may serve as an attractive vaccine scaffold due to its highly programmable geometry and size^49, 50^, quantitative spatial precision for display of biomolecular cargoes^51,52^, and immunologically inert TI scaffold. Accordingly, we hypothesized that DNA origami scaffolds could enable a rigorous test of the effects of scaffold-specific competitor B cells on the recruitment and affinity maturation of rare “on-target” precursors in primary GCs.

To test our hypothesis, we rationally designed icosahedral DNA-VLPs conjugated with eOD-GT8 as a clinically-relevant test immunogen. We identified a set of DNA-VLP design rules, principally optimizing antigen density, particle size, and incorporating synthetic T cell helper epitopes to trigger follicular targeting of the VLPs, which led to 15-fold higher frequencies of antigen-specific GC B cells compared to eOD-GT8 monomer. In a humanized mouse model of VRC01-class B cell priming, we found that GCs formed by scaffold-silent DNA-VLPs contained 25-fold higher ratios of on-target epitope-specific B cells to competitor B cells — approximately 12 times higher than the protein-based eOD-GT8 60mer nanoparticles that elicited significant scaffold-specific competitor responses. Further, DNA-VLPs effectively expanded VRC01-class precursors within 14 days following low-dose immunization, conditions where the protein nanoparticle failed to expand bnAb-lineage B cells. Thus, eliminating competitive scaffold-specific humoral responses can have a substantial impact on the priming of rare B cell precursors, with DNA-VLPs showing promise as a platform technology to prime antigen-focused germinal center responses.

## Scaffolding eOD-GT8 on DNA origami virus-like particles

We previously demonstrated that SARS-CoV-2 receptor-binding domain multimerized on icosahedral DNA-VLPs (30 antigens/nanoparticle) elicited high serum IgG titers in mice after a prime-boost vaccination that potently neutralized the virus^28^. To begin exploring how DNA-VLP design impacts earlier stages of the immune response, and particularly the formation of primary germinal center responses that are crucial to effective HIV vaccines, we employed the same DNA-VLP and surface-conjugated 30 equally spaced copies of the eOD-GT8 antigen to the particles (Fig. 1a). This wireframe DNA origami nanoparticle was computationally designed using DAEDALUS software^49, 53^ and synthesized following established protocols, utilizing a custom ssDNA scaffold folded with 30 DBCO-modified staple oligonucleotides and reacted with eOD-GT8 containing a C-terminal azido linker^51, 52^ (Supplementary Fig. 1). Successful DNA-VLP self-assembly was confirmed through agarose gel electrophoresis (AGE) and dynamic light scattering (Extended Data Fig. 1a-c). Post-antigen coupling, the hydrodynamic radius of the nanoparticles was ∼40 nm, which we annotate as d40_30mer. The percentage of alkyne groups on the DNA-VLP successfully conjugated with antigen was quantified using a bicinchoninic acid protein assay; functionalization efficiency across independent sample preparations was consistently in the range of 90-100%. The measured antigen concentrations were used to standardize eOD-GT8 doses across experimental groups (Extended Data Fig. 1d). Cryo-electron microscopy and negative stain transmission electron microscopy validated the preservation of the icosahedral geometry of the origami structure post-antigen conjugation (Fig. 1b, Extended Data Fig. 1e). The bioavailability and proper orientation of the antigen were further confirmed through binding assays with murine VRC01, a broadly neutralizing antibody targeting the CD4 binding site of gp120 that binds with high affinity to eOD-GT8^39^. VRC01 antibody successfully bound eOD-GT8 displayed on DNA-VLPs, as evidenced by decreased mobility in gel electrophoresis (Supplementary Fig. 2a). As a functional quality control metric for antigen bioactivity, we incubated antigen-loaded particles with Ramos B cells expressing the germline VRC01 B cell receptor, and measured calcium flux using a calcium-dependent fluorescent dye. Consistent with previous findings^44^, culture of VRC01-expressing Ramos B cells with 5 nM eOD-GT8-functionalized DNA-VLPs led to rapid B cell activation as assessed by calcium signaling, while the same dose of soluble eOD-GT8 monomer failed to trigger calcium release (Fig. 1c).

**Fig 1.**
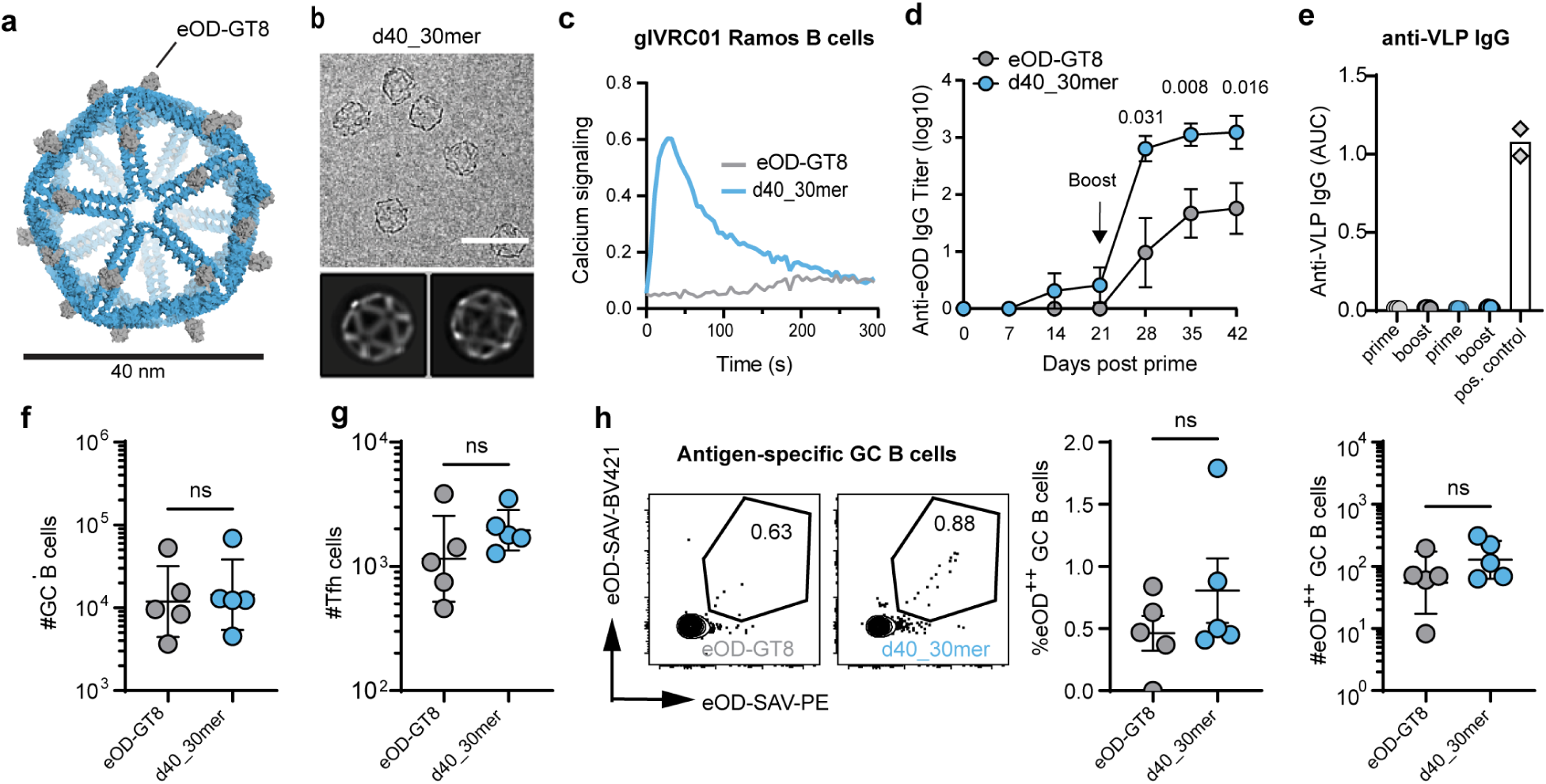
Nanoparticulate eOD-GT8 assembly on icosahedral DNA-VLP scaffolds. **a,** Atomic model of d40_30mer, which consists of an icosahedral DNA origami nanoparticle (52 base pairs/edge) with one eOD-GT8 antigen conjugated per edge of origami using SPAAC chemistry. **b,** Cryo-electron micrographs of d40_30mer and 2D class averages. The scale bar represents 50 nm. **c,** Representative calcium flux trace of glVRC01 B cells incubated with 5 nM eOD-GT8 monomer or d40_30mer from *n*=3 independent experiments. Traces were normalized to fluorescence background measured prior to antigen addition. **d,** C57BL/6 mice (*n*=5/group) were injected subcutaneously (s.c.) with equimolar doses of monomer or d40_30mer (equivalent to 5 μg of eOD-GT8 monomer) together with 5 μg SMNP adjuvant. Blood was collected weekly for serum eOD-GT8 antibody titer measurement by ELISA. Statistical significance was calculated using multiple Mann Whitney tests with Benjamini, Krieger, Yekutieli correction. **e,** Anti-origami IgG responses represented as area under the curve (log10) after prime (day 21) and boost (day 42) time points. An anti-dsDNA antibody was used as positive control. **f,** C57BL/6 mice (*n*=5/group) were primed with 5 μg of monomer or d40_30mer and sacrificed on day 14 for analysis of GC responses by flow cytometry. Total counts of GC B cells (B220^+^/CD38^lo^GL7^hi^). **g,** Total counts of Tfh cells (CD4^+^/CXCR5^hi^ PD1^hi^). **h,** Representative flow plots showing eOD-tetramer staining, frequency of eOD^++^ GC B cells out of all GC B cells and counts of eOD^++^ GC B cells. Statistical testing was performed with Mann Whitney U tests.

## d40_30mer DNA-VLPs enhance serum antibody titers in prime-boost immunization but are weak priming immunogens

Based on the results from *in vitro* B cell activation and previous *in vivo* studies of eOD-GT8 protein nanoparticles^25^, we hypothesized that d40_30mer would induce higher antibody responses in mice compared to eOD-GT8 monomer due to its multivalency. For immunization studies, we employed a saponin-MPLA nanoparticle adjuvant (SMNP), a potent ISCOMs-like adjuvant^54^. We verified that mixing DNA-VLPs with SMNP did not affect the stability of the DNA-VLPs (Supplementary Fig. 2b). C57BL/6 mice were primed and boosted with equimolar doses of eOD-GT8 monomer or d40_30mer co-administered with SMNP, and antigen-specific serum IgG titers were monitored over time. Both eOD-GT8 monomer control and DNA-VLPs resulted in weak responses post-prime, but following boosting the DNA-VLP elicited endpoint antibody titers ∼1 log higher than the eOD-GT8 monomer (Fig. 1d). We additionally assessed antibody responses against the bare DNA-VLP, and consistent with our prior studies^28^, we observed no elevation in anti-origami IgG or anti-dsDNA IgG after prime or boost (Fig. 1e, Extended Data Fig. 2a,b). We also measured IgM responses to dsDNA and detected no response above the nonspecific background triggered by protein eOD-GT8/SMNP immunization in the complete absence of DNA (Extended Data Fig. 2c).

To gain insight into the early stages of the response to monomer vs. DNA-VLP immunization, we evaluated germinal center responses in draining lymph nodes at two weeks after a single immunization by flow cytometry (Supplementary Fig. 3). Unexpectedly, immunization with d40_30mer did not increase the number of GC B cells (B220^+^/CD38^lo^ GL7^hi^) or follicular helper T cells (CD4^+^/CXCR5^hi^ PD1^hi^) relative to eOD-GT8 monomer vaccination (Fig. 1f-g). The frequency and counts of antigen-specific GC B cells, which were identified by fluorescent eOD-tetramer staining, were also not enriched and remained low— below 1% (Fig. 1h). Alternative adjuvants including alum, AddaVax, AS01b, and CpG ODN 1826 also did not enhance GC B cell responses elicited by the DNA-VLPs (Extended Data Fig. 3).

A primary concern for DNA origami-based nanoparticles *in vivo* is whether they remain stable over a sufficient time period in the presence of DNAses present in tissues. To test whether the priming efficiency of DNA-VLPs was limited by endonuclease-mediated degradation, we assessed vaccine responses in DNase I-deficient mice (Extended Data Fig. 4a)^55, 56^. We verified via serum stability assays that d40_30mer remained intact in DNase I KO serum for longer than seven days (Extended Data Fig. 4b). Further, we validated that mutant DNase I KO mice established robust GCs in response to protein nanoparticle vaccination identically to WT controls (Extended Data Fig. 4c). To determine if DNAse-mediated particle degradation was a limiting factor *in vivo*, we primed WT or DNase I KO mice with d40_30mer and analyzed germinal centers after two weeks. Surprisingly, total GC size, Tfh responses, and antigen-specific GC B cells formed by d40_30mer in DNase I KO animals were low and identical to responses in WT littermates (Extended Data Fig. 4d). These data suggested that the d40_30mer nanoparticle design may provide sufficient avidity to expand antibody-producing cells upon boosting, but the nanoparticle design, irrespective of enzymatic stability, is not optimal for supporting primary GC responses.

## d40_30mer DNA-VLPs have restricted access to follicular dendritic cells

To understand why the d40_30mer DNA-VLP poorly primed GC responses, we considered the physiological barriers limiting antigen delivery to B cells *in vivo*. Antigen arriving at draining lymph nodes during primary immunization is often quickly cleared by lymph flow, protease activity in the tissue, or by endocytosis and degradation by lymphocytes^20, 57-59^. Alternatively, antibody- or complement-opsonized antigen can be captured by follicular dendritic cells (FDCs) and presented to B cells in germinal centers for prolonged periods to support the GC response^60-63^. We thus analyzed antigen biodistribution in draining lymph nodes (dLNs) following immunization, comparing eOD-GT8 monomer and d40_30mer DNA-VLPs with the protein nanoparticle eOD-GT8 60mer (hereafter, p60mer), which we have previously shown becomes decorated with complement *in vivo* via the lectin pathway, leading to rapid robust accumulation on FDCs (which express high levels of complement receptor)^61^. To better understand the *in vivo* fate of DNA-VLPs versus protein immunogens, we immunized mice with fluorescently tagged antigens and collected inguinal dLNs two hours later for cryo-sectioning and immunofluorescence staining. Early after immunization, eOD-GT8 monomer was primarily detected in central non-follicular areas of dLNs, largely colocalizing with F4/80^+^ medullary sinus macrophages (MSMs) (Fig. 2a, b). By contrast, eOD-GT8 delivered as d40_30mer was present at substantially higher levels in the draining lymph nodes, and colocalized with CD169^+^ subcapsular sinus macrophages (SSMs) lining the edges of the lymph node and MSMs (Fig. 2a, b). The p60mer particle also colocalized with SSMs but was already at this early timepoint accumulating on the dendrites of CD35^+^ FDCs (Fig. 2a lower panel, white arrows). Quantitative pixel analysis revealed that d40_30mer and p60mer had similar levels of antigen colocalization with SSMs (Fig. 2c), but only p60mer showed enrichment on FDCs (Fig. 2c). At later time points such as day 7, p60mer had become highly concentrated on FDCs, whereas follicles in lymph nodes immunized with d40_30mer had no retention of fluorescent antigen (Fig. 2d). Thus, antigen trafficking and retention of DNA-VLPs was distinct from similarly-sized protein nanoparticles.

**Fig. 2.**
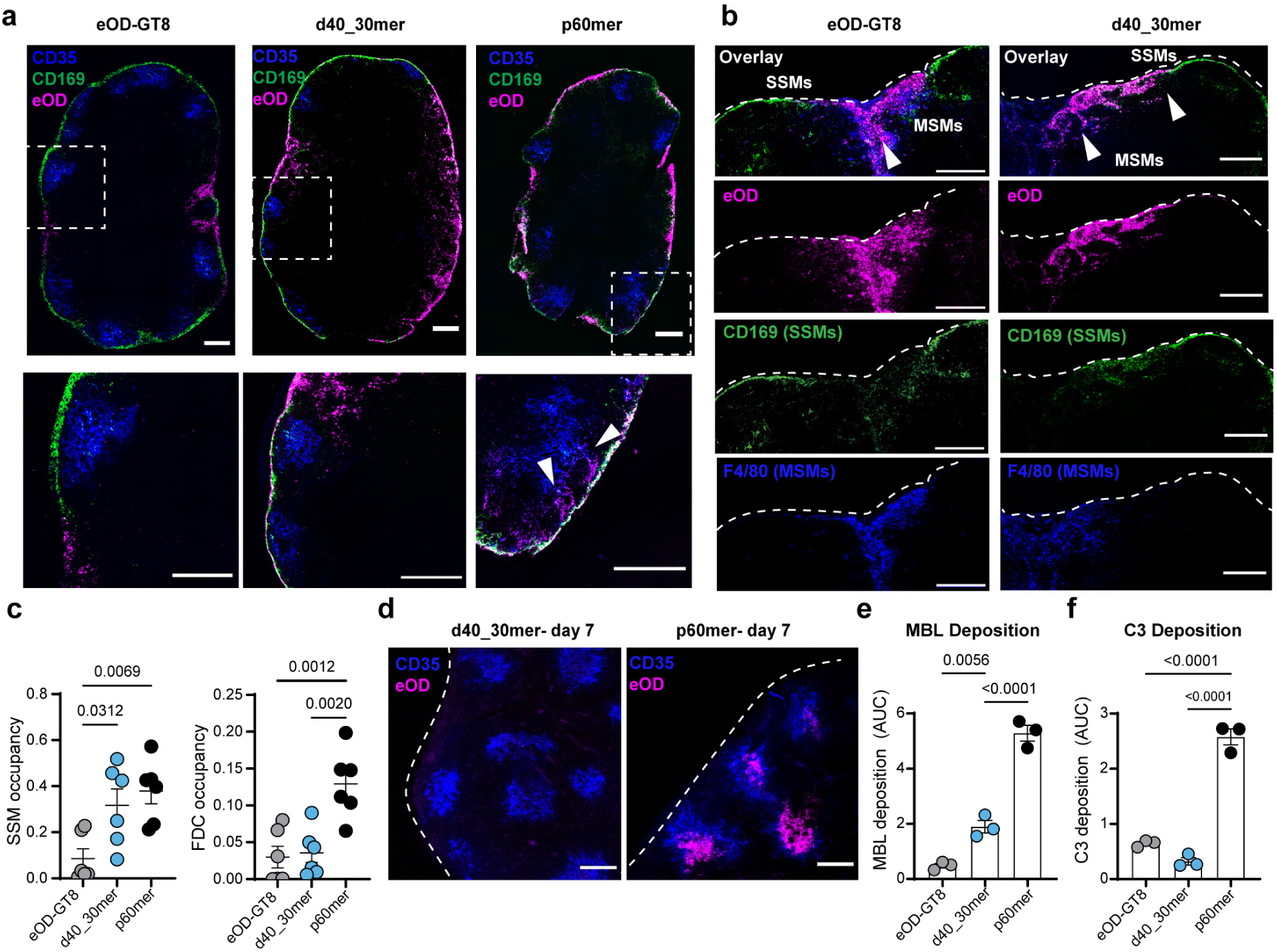
d40_30mer DNA-VLPs are poorly retained on follicular dendritic cells in draining lymph nodes. **a,** C57BL/6 mice (*n*=3 mice, 6 lymph nodes/group) were primed s.c. with AF647-labeled eOD-GT8, d40_30mer, or p60mer. At 2 hr post-injection, inguinal lymph nodes were flash frozen, cryosectioned, and stained with anti-CD35 antibody (blue) and anti-CD169 antibody (green). Shown are representative lymph node images collected on a laser scanning confocal microscope, with close-up images of subcapsular sinus regions and follicles (lower panel). White arrows point to antigen (magenta) colocalized with FDC dendrites. Scale bars represent 200 μm. **b,** Lymph node sections from mice immunized with AF647-labeled eOD-GT8 monomer or d40_30mer (magenta) were stained with anti-CD169 (green) and anti-F4/80 (blue) antibodies for visualization of subcapsular sinus or medullary sinus macrophages, respectively. Arrows point to antigen colocalization with MSM or SSM populations. **c,** Quantification of fraction of CD169^+^ SSMs and CD35^+^ FDCs containing antigen at 2 hrs. **d,** Optically cleared inguinal LNs 7 days after immunization with AF647-labeled d40_30mer or p60mer. Follicles were labeled *in situ* by pre-injection of anti-CD35 antibody prior to fixing and clearing. Scale bar = 200 μm. **e,** MBL deposition measured by ELISA. Antigen formulations were immobilized on assay plates and treated with dilutions of fresh mouse serum. Anti-MBL antibody was used to detect MBL deposited on the antigens. **f,** C3 deposition measured by ELISA. Antigen formulations were immobilized on assay plates and treated with dilutions of fresh mouse serum. Anti-C3 antibody was used to detect C3 bound to the antigens. Statistics were calculated with one-way ANOVA with post-hoc Tukey test.

We hypothesized that trapping of d40_30mer in the subcapsular sinus could be due to high expression of DNA-binding scavenger receptors on macrophages and/or lymphatic endothelial cells^64^ or inadequate triggering of complement pathways to mediate the capture of the particles by FDCs^62^. To test the first hypothesis, we passivated d40_30mer by coating it with polylysine-PEG polymer^65^, which has been used *in vivo* with other DNA origami formulations for stabilization, charge neutralization^66, 67^, and avoidance of macrophage uptake^68, 69^. PEGylated d40_30mer exhibited enhanced stability in mouse serum and reduced association with murine macrophage cells *in vitro* (Extended Data Fig. 5a-c). However, d40_30mer nanoparticles with or without PEGylation showed similar patterns of early accumulation in medullary and subcapsular sinus regions (Supplementary Fig. 5d), and PEG-coated DNA-VLPs elicited weaker germinal centers compared to the uncoated d40_30mer (Extended Data Fig. 5e-h), likely due to interference of the polylysine-PEG coating with antigen recognition on BCRs^70^.

Heavily glycosylated nanoparticle-antigens become decorated with complement *in vivo* via the lectin pathway, when mannose binding lectin (MBL, an innate immune protein present in blood and lymph) binds to the particles and triggers complement deposition^61, 71^. To test the hypothesis of insufficient complement activation by the DNA-VLPs, we measured binding of MBL and C3 from naïve mouse serum to eOD-GT8 monomer, d40_30mer, or p60mer particles immobilized on ELISA plates. MBL and C3 both showed significantly lower binding to eOD-GT8 monomer and d40_30mer compared to p60mer (Fig. 2e, f). Similarly, very little binding of recombinant MBL to eOD-GT8 monomer or d40_30mer was detected by ELISA compared to p60mer (Extended data Fig. 6). Thus, DNA-VLPs exhibited considerably less effective engagement of the lectin pathway for complement activation than the p60mer nanoparticles.

## Engineering DNA-VLPs with high antigen density induces follicle targeting and augments antigen-specific GC B cell responses

We hypothesized that poor complement activation explained the lack of FDC accumulation of the DNA-VLPs, and that this was a result of insufficient antigen density on the DNA origami scaffold^71^. MBL is a large protein complex consisting of trimeric stalks containing a collagen domain and carbohydrate-binding domain (CBD), which oligomerize forming hexamers. Each CBD in the stalk trimer is estimated to be ∼5 nm apart from its neighbor with similar or larger distances to CBDs in neighboring stalks of the oligomer^72^. Each CBD has weak, millimolar affinity for mannose, but strong binding can be achieved through the avidity of multiple CBD stalks binding to numerous glycans^61^; thus MBL binding is highly sensitive to the density of glycans on a viral or bacterial surface. To assess the role of antigen density on MBL binding and complement activation by DNA-VLPs, we synthesized a set of 4 different particles, with diameters of approximately ∼23 or ∼34 nm, functionalized with either 30 or 60 copies of eOD-GT8 (Fig. 3a), providing a range of predicted glycan densities (0.04 to 0.14 glycans/nm^2^) and a range of inter-antigen distances from 11 nm (the original d40_30mer) down to 4.5-6 nm between neighboring antigens (d30_60mer, Fig. 3b). Upon antigen conjugation, which was quantified by BCA (Extended Data Fig. 1d), the hydrodynamic diameter of these particles was measured by DLS to be approximately ∼30 nm and ∼40 nm, respectively, denoted as d30 and d40 (Extended Data Fig. 1c). Each of the 4 DNA-VLP designs had sufficient avidity to robustly activate cognate B cells and elicited indistinguishable calcium flux in germline VRC01-Ramos B cells *in vitro* (Fig. 3c). Amongst these designs, antigen attachment sites in d30_60mer are spaced similarly to the distances between CBD domains on MBL, and this design also closely resembles the geometric configuration of eOD-GT8 antigen on lumazine synthase p60mer.

**Fig. 3.**
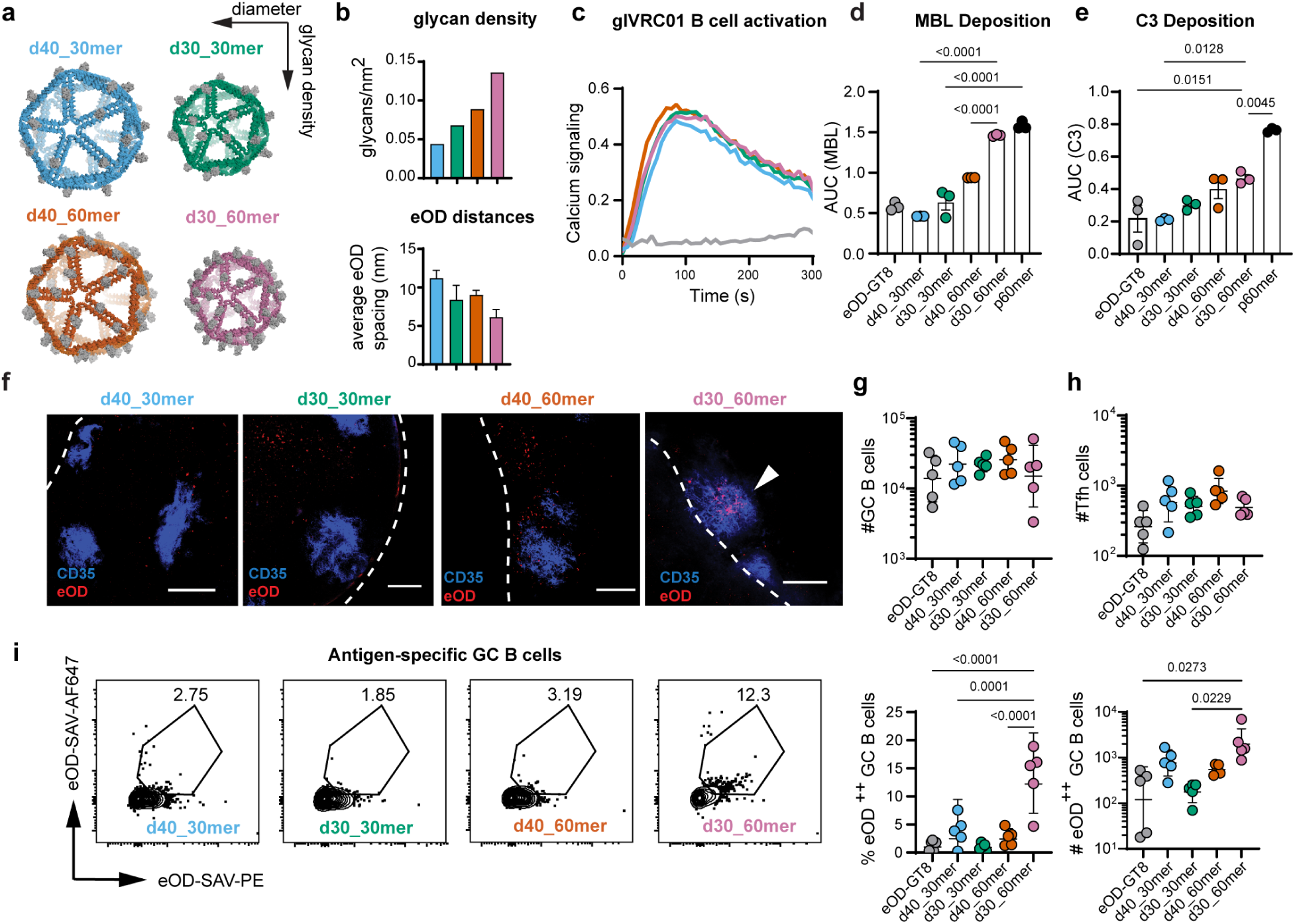
High density antigen display enhances lectin pathway activation, FDC targeting, and antigen-specific GC B cell responses. **a,** Atomic models of icosahedral DNA origami nanoparticles with diameters of 30 or 40 nm (d30_ and d40_) and eOD-GT8 valency of 30 or 60. **b,** Predicted glycan densities assuming 4 glycan sites per eOD and inter-antigen distances calculated as the average distance to neighboring 5 antigens based on atomic models. **c,** Representative calcium signaling of glVRC01 B cells incubated with eOD formulations from 3 independent experiments. **d,** MBL deposition on different VLPs following incubation in fresh mouse serum detected by ELISA. Shown is the area under the curve of three technical replicates. **e,** C3 deposition on different VLP formulations, assayed by incubating in dilutions of fresh mouse serum and detected by ELISA. Shown is the area under the curve from three technical replicates. **f,** DNA-VLPs were formulated with AF647-labeled eOD-GT8. C57BL/6 mice (*n*=3, 6 lymph nodes/group) were immunized s.c. with 10 μg of antigen with SMNP adjuvant. Anti-CD35 antibody was injected prior to necropsy for in situ labeling of FDCs. Inguinal lymph nodes were collected at 96 hrs and cleared for whole tissue imaging. Shown are representative follicles (projection over 100 μm). Scale bar represents 200 μm. **g, (**C57BL/6 mice (*n*=5/group) were primed with equimolar antigen doses (equivalent to 5 μg eOD-GT8) with SMNP adjuvant. Inguinal LNs were analyzed on day 14. Shown are total counts of GC B cells (B220+ CD38 lo GL7 hi) and **h**, total counts of Tfh cells (CD4+/CXCR5+ PD1+). Absolute counts plots are shown with log axes; geometric mean and geometric S.D. are shown **i,** Representative flow plots of eOD-tetramer staining (left) and frequencies and counts of eOD^++^ GC B cells. Absolute counts plots are shown with log axes; geometric mean and geometric S.D. are shown Statistical significance was determined by 1-way ANOVA with post-hoc Tukey test.

We conducted complement deposition assays with fresh mouse serum to evaluate how antigen spacing affected MBL and C3 deposition on the DNA-VLP surfaces, using the p60mer nanoparticle as a positive control for the assay. We observed a monotonic increase in both MBL and C3 deposition on DNA-VLPs as antigen density increased, with the highest amount of complement depositing on d30_60mer (Fig. 3d, e, Extended data Fig. 6b). MBL binding with d30_60mer was nearly identical to p60mer; however, the amount of C3 opsonized on DNA-VLPs was significantly lower. Next, we immunized mice with the panel of DNA-VLPs and harvested dLNs for cleared lymph node imaging at day 4 after injection. Strikingly, fluorescent eOD-GT8 signal was detected in lymph nodes from all groups, but colocalization with FDCs was only observed for d30_60mer particles (Fig. 3f). Thus, as previously observed with other synthetically mannosylated protein nanoparticles^71^, stronger deposition of MBL on DNA-VLPs, which can be increased by increasing antigen density, correlates with increased antigen capture on FDCs *in vivo*.

Next, we investigated how antigen density impacts germinal center formation. Mice immunized with equimolar antigen doses of eOD-GT8 monomer or DNA-VLPs showed similar total counts of GC B cells and Tfh cells two weeks post immunization (Fig. 3g, h). However, we observed a ∼15-fold increase in the frequency and 10-fold increase in total counts of eOD-specific GC B cells elicited by the d30_60mer compared to the eOD-GT8 monomer control (Fig. 3i). Thus, a DNA-VLP design that promoted particle capture by FDCs correlated with substantially stronger expansion of antigen-specific B cells in active GCs. Furthermore, this effect was driven by antigen density, rather than valency, since the d40_60mer failed to elicit antigen capture on FDCs or expansion of antigen-specific cells.

## Synthetic T cell epitopes provide focused T cell help to antigen-specific GC B cells

In addition to retention of antigen on FDCs, stimulation of GC B cells by helper CD4 T cells is a critical cue for driving strong germinal center reactions^9, 73^. DNA-VLPs are T-independent scaffolds; therefore, all T cell epitopes must be derived from the conjugated protein antigen. In contrast, protein scaffolds such as lumazine synthase that forms the basis of the p60mer provide abundant scaffold-derived T cell help^74^. Consequently, following p60mer immunization, antigen-specific B cells compete with scaffold-specific B cells for co-stimulation from the same pool of CD4 T cells recognizing scaffold-derived peptides. Given the small size of the eOD-GT8 antigen, which may provide insufficient T cell help on its own, we next tested whether the incorporation of the universal helper peptide pan-HLA DR-binding epitope (PADRE) on the terminus of the antigen could further amplify GC responses elicited by DNA-VLPs without introducing off-target B cell epitopes. PADRE binds with high affinity to a broad spectrum of human and mouse MHC haplotypes and has been shown to be safe in human clinical trials^75-77^. We tested this hypothesis by comparing GC responses induced by three different 60mer formulations: d30_60mer, d30_60mer carrying eOD-PADRE (d30_60mer-PADRE), and p60mer. Fusion of the T cell epitope to eOD-GT8 had no effect on formation of the DNA-VLPs (polydispersity, diameter, or antigen functionalization efficiency, Extended Fig. 1f). Furthermore, all three 60mers strongly activated germline VRC01 Ramos B cells *in vitro* (Fig. 4a) and bound MBL and C3 in mouse serum (Extended Data Fig. 6c).

**Fig. 4.**
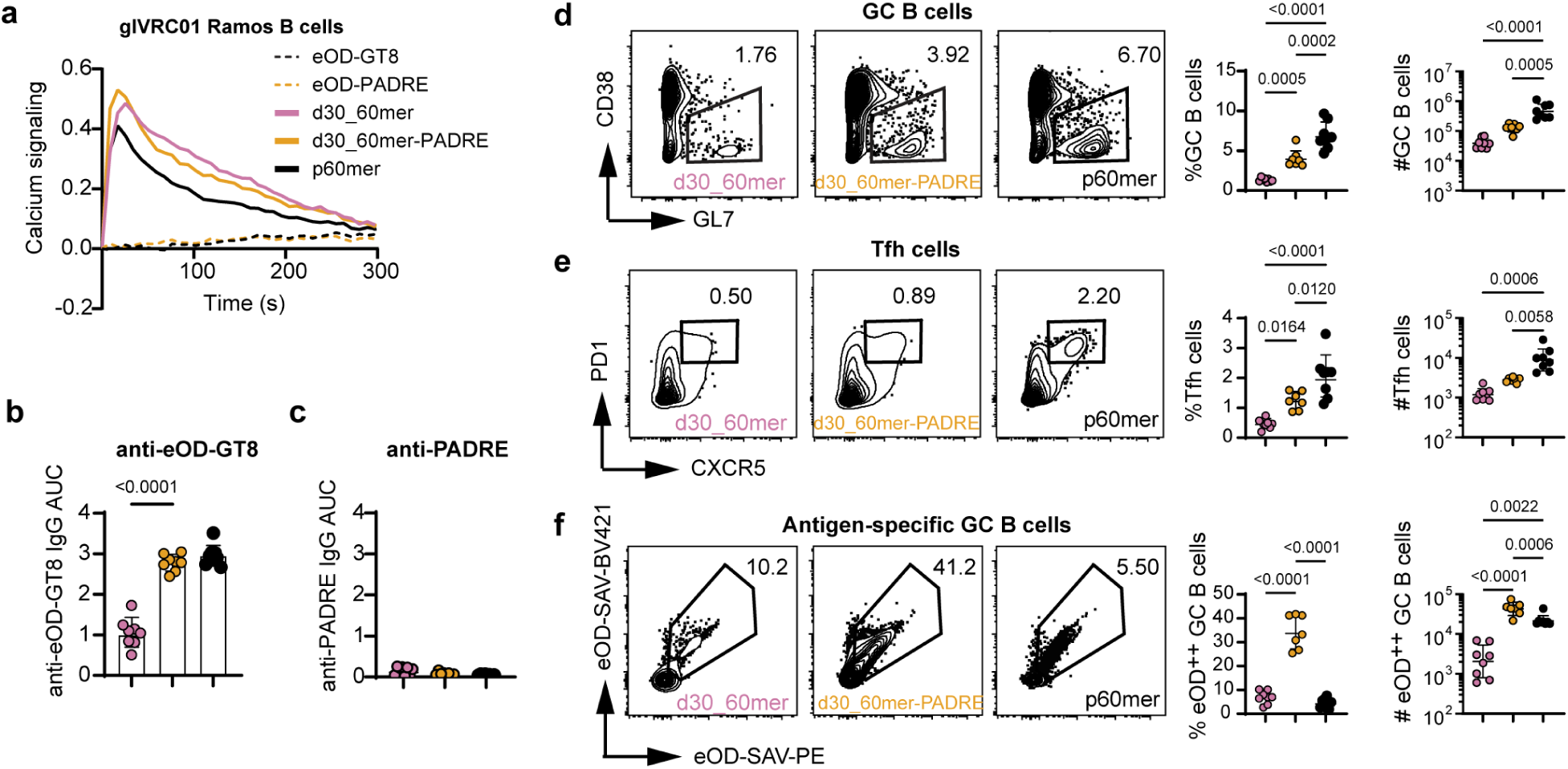
Incorporating synthetic helper T cell epitopes boosts GC size after DNA-VLP vaccination. **a,** Representative calcium signaling traces from VRC01 Ramos B cells incubated with 5 nM eOD-GT8 equivalent concentrations of indicated particles (*n*=3 independent experiments). **b,** C57BL/6 mice (*n*=8 mice/group) were primed s.c. with equimolar doses (equivalent to 5 ug of eOD) of the 60mer formulations with SMNP and sacrificed on day 14 for serum titer analysis and germinal center analysis by flow cytometry. Anti-eOD-GT8 IgG titers were determined by ELISA (against eOD-GT8 without PADRE). Shown are AUC values (log10). **c,** Anti-PADRE IgG titers determined by ELISA against biotinylated PADRE peptide. Shown are AUC values (log10). **d,** Representative flow plots, frequencies (%GC out of total B cells), and counts of germinal center B cells (B220+/CD38lo GL7^hi^). **e,** Representative flow plots, frequencies (%Tfh out of CD4 T cells), and counts of Tfh cells (CD4^+^ /CXCR5^+^ PD1^+^). **f,** Representative flow plots, frequencies (%eOD^++^ out of GC B cells), and counts of antigen-specific eOD^++^ GC B cells. Absolute counts plots are shown with log axes; geometric mean and geometric S.D. are shown. Data was analyzed with one-way ANOVA with post-hoc Tukey test.

We primed mice with the three 60mer formulations and assessed serum IgG titers by ELISA and GC responses by flow cytometry. The addition of PADRE significantly enhanced early anti-eOD-GT8 IgG responses compared to d30_60mer (Fig. 4b), while minimal IgG responses were detected against the PADRE epitope itself (Fig. 4c). Mice primed with d30_60mer-PADRE also had significantly increased frequencies and total counts of GC B cells and Tfh cells compared to d30_60mer, though the total GC B cell count remained 4-fold lower than that elicited by LumSyn-scaffolded p60mer (Fig. 4d, e). However, GCs initiated by d30_60mer-PADRE expanded a much higher frequency of eOD-specific GC B cells (30-40%) compared to p60mer (∼5%, Fig. 4f). This was also reflected in the total counts of eOD-specific GC B cells, which were highest in d30-60mer-PADRE immunizations (2.1-fold greater than p60mer), despite the induction of an overall smaller GC response than that produced by p60mer. Thus, engineering of T cell help in DNA-VLPs enabled priming of a robust antigen-focused GC response.

## DNA-VLPs prime epitope-focused germinal centers in humanized mice

We hypothesized that the increased frequency of antigen-specific B cells in germinal centers activated by DNA-scaffolded compared to protein-scaffolded antigen may reflect an “immune focusing” effect of DNA-VLP immunization due to the inert nature of the DNA origami scaffold^28^. To directly assess competitor versus “on-target” epitope-specific B cell responses directed to the CD4 binding site (CD4bs), we analyzed germinal center responses to protein- or DNA-scaffolded immunogens in transgenic mice expressing the germline human IGHV1-2*02 gene segment knocked into the mouse Ig locus (VH1-2 mice), which is paired with endogenous mouse light chains. This mouse model is designed to assess the engagement of engineered germline targeting immunogens with a diverse repertoire of VRC01-class B cell precursors expressing human heavy chains, modeling the diversity of potential bnAb precursor B cells present in the human B cell repertoire, with a low frequency of bona fide bnAb precursors^78^. We immunized VH1-2 mice with d30_60mer-PADRE or p60mer and analyzed their GC responses in draining lymph nodes, using flow probes to characterize antigen-specific vs. competitor B cells in GCs (Fig. 5a).

**Fig. 5.**
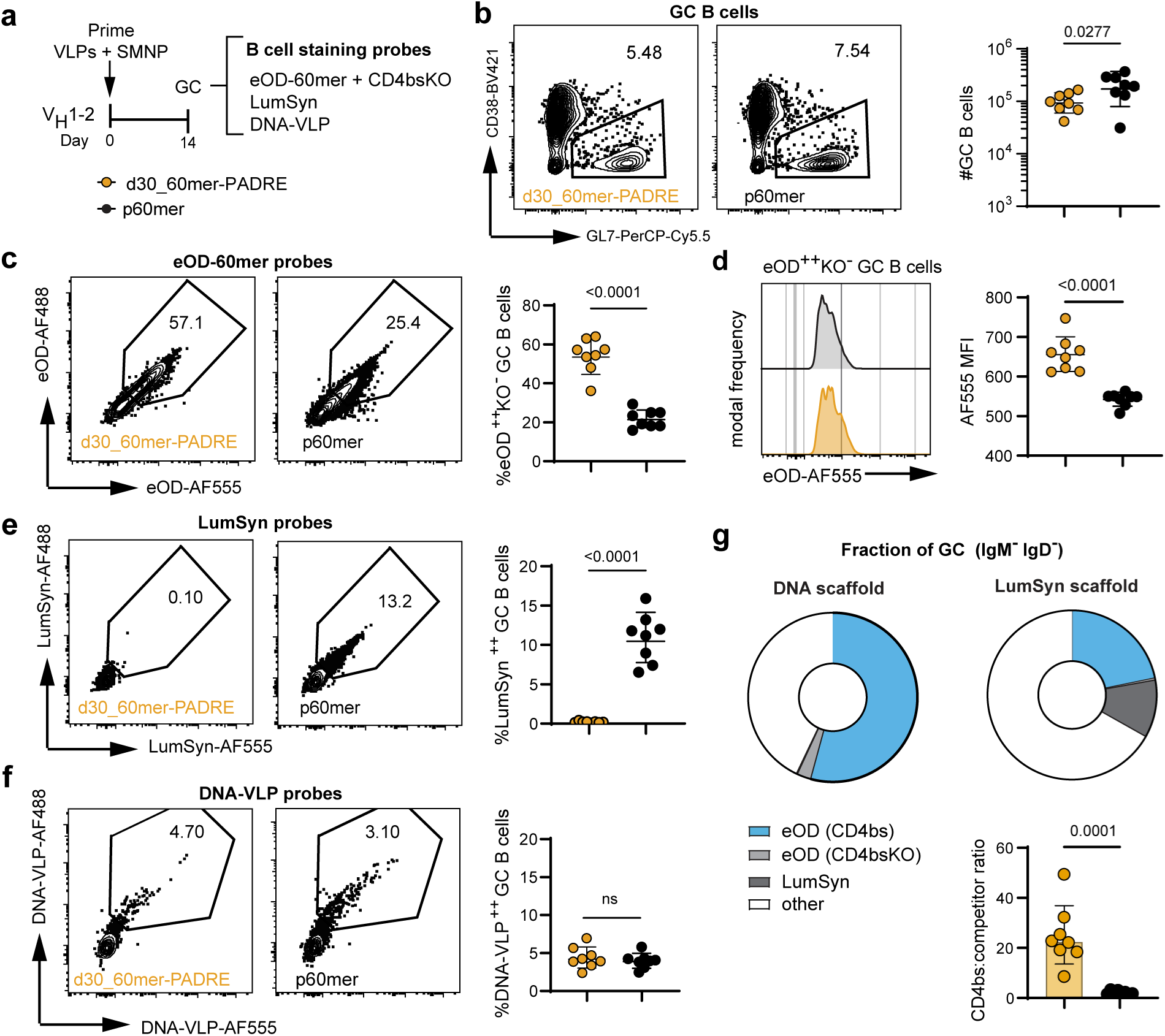
DNA-VLPs produce focused germinal centers and prime VRC01-class precursors in humanized mice. **a,** Humanized V_H_1-2 mice (*n*=8/group) were primed s.c. with 5 μg d30_60mer-PADRE or p60mer with SMNP, and inguinal lymph nodes were harvested on day 14. The lymphocytes were split into four samples and each stained with GC markers and fluorescently labeled antigen probes (eOD-60mer, bare lumazine synthase, or bare DNA-VLP (d30) probes). **b,** Total counts of B220^+^/CD38^lo^GL7^hi^ GC B cells. **c**, Representative eOD-60mer probe staining of GC B cells (left) and frequency of eOD-60mer^++^ cells out of IgM^-^ IgD^-^ GC B cells (right). **d,** Frequency of eOD-60mer^++^ eOD-CD4bs-KO^-^ (CD4bs+) out of IgM^-^ IgD^-^ GC B cells. **e,** Representative LumSyn staining of GC B cells (left) and frequency of LumSyn^++^ cells out of IgM^-^ IgD^-^ GC B cells (right). **f,** Representative DNA-VLP (d30) staining of GC B cells (left) and frequency of DNA-VLP^++^ cells out of IgM^-^IgD^-^ GC B cells (right). Statistical significance was determined using one-way ANOVA with post-hoc Tukey test for b-f. **g**, Fraction of germinal center B cells that bound to each set of antigen or scaffold probes represented as parts-of-whole plot. The green slice represents on-target CD4bs-specific GC B cells. All other colors represent competitor B cells (eOD-CD4bs-KO or scaffold-specific B cells) or other cells not reactive with any probes.

To assess on-target responses, we analyzed the staining of IgM^-^IgD^-^ GC B cells by fluorescently labeled eOD-60mer nanoparticle probes and an eOD-60mer-CD4bsKO nanoparticle probe, the latter of which contain mutations in the CD4 binding site that ablate binding by true VRC01-class precursor B cells; binding to this KO probe identifies B cells specific to eOD epitopes other than the CD4bs. Importantly, it has been previously observed that antibodies previously identified by intact eOD-60mer nanoparticle probes do not bind lumazine-specific B cells^30^. This suggests that scaffold-specific responses primed by p60mer represent B cells which recognize breakdown products of the protein nanoparticle that we have previously shown are rapidly generated by extrafollicular proteases in the lymph node ^59^. For quantifying scaffold-specific competitor B cells, we used fluorescently labeled bare lumazine synthase or bare DNA-VLPs to stain GC B cells. Comparing GC B cell binding to these different sets of nanoparticle probes quantifies relative frequencies of “on-target” CD4bs-specific (eOD^++^KO^-^) and “off-target” (CD4bs-KO^+^ or scaffold-specific) clones in GCs.

As observed in WT mice, the p60mer primed overall slightly larger total GC responses (2-fold greater than the DNA-VLP, Fig. 5b). However, we observed that the composition of GCs elicited by protein- and DNA-scaffolded antigen was very distinct: GCs formed in mice immunized with d30_60mer-PADRE were composed of nearly ∼60% on-target CD4bs-specific GC B cells (identified by eOD^++^KO^-^ gate), compared to ∼20% expanded by p60mer (Fig. 5c). Furthermore, the on-target B cells expanded by the DNA-VLP formulation had stronger staining by the antigen probes represented by higher mean fluorescence intensity, suggesting that these cells might have undergone higher extents of affinity maturation (Fig. 5d). Following p60mer immunization, LumSyn-specific B cells contributed to ∼13% of the GC, comparable in magnitude to the on-target CD4bs-directed response (Fig. 5d). When staining with bare DNA-VLP probes, some background staining was detected in GC B cells, but this staining was the same in mice primed with either DNA-VLPs and p60mer, and similar low levels of background binding were detected on B cells from naïve mice (Fig. 5f, Supplementary Fig. 7), which may reflect low-level scavenger receptor expression in B cells^79^. Compared to the p60mer, DNA-VLP immunization promoted germinal centers that contained 25-fold higher ratios of on-target CD4bs-specific B cells to off-target competitors (Fig. 5g), whereas in p60mer the frequency of epitope-specific B cells was only enriched 2-fold.

## eOD display by DNA-VLPs leads to more efficient priming of bnAb precursors and robust somatic hypermutation

Finally, we inquired whether the epitope-focused GCs produced by DNA-VLPs led to better expansion and somatic hypermutation (SHM) of VRC01-class precursors, which are defined by IGHV1-2*02 usage combined with 5-amino acid CDRL3 light chains. The VH1-2 mouse model has a diversity of V(D)J-rearranged human VH1-2 heavy chains and a fully murine LC repertoire, where only about 0.15% of B cells express 5-AA light chains in the naïve repertoire^78^, thus providing a stringent model for expanding VRC01-class precursors. We immunized VH1-2 mice with 5 μg (eOD-GT8 equivalent) of either d30_60mer-PADRE or p60mer, sorted class-switched antigen-specific GC B cells (B220^+^/CD38^lo^GL7^hi^/IgM^-^IgG^+^/eOD^++^KO^-^) on day 14, and performed single-cell sequencing to analyze the BCR repertoire. A total of 444 and 465 BCR sequences were recovered and analyzed for d30_60mer-PADRE and p60mer, respectively.

We found diverse BCR clonotypes (defined as shared CDRH3 usage) expanded by each immunogen (Fig. 6a, c). However, in DNA-VLP vaccination, several clonotypes contained significantly more expanded representatives (Fig. 6a), suggesting proliferation of clonally related B cell lineages in the GCs, aligning with our hypothesis of epitope-focused GCs. We further note that within these clonotypes, DNA-VLP immunization enriched for the five AA CDRL3 VRC01 class signature (Fig. 6b), whereas the p60mer did not (Fig. 6d). The p60mer was previously reported to enrich for this signature VH1-2 mice after a single prime immunization but this required antigen doses of 30 ug or more and 2-8 weeks^78^. Importantly, within the clonotypes expanded by DNA-VLP vaccination, the five amino acid CDRL3 signature was enriched by expansion of public B cell clones (BCRs with shared CDRH3 + CDRL3 within the independent vaccine recipients; Clone IDs - 70, 78, 116, 62, 19 and 137) (Fig. 6a, right, Extended Data Fig. 7). Expansion of multiple genetically identical B cell clones in different mice points to a reproducible mechanism for triggering of VRC01 bnAb class precursors with DNA-VLPs. The 5 AA CDRL3 was supplied by usage of mouse IGKV4-61*01 (Extended Data Fig. 7), as described in a previous study^40^. We also observed SHM in the vaccine-expanded B cells primed with both formulations, but they were not significantly different at this early timepoint. We also identified enrichment of key VRC01-class HC mutations^80^ in both formulations, particularly S55R/N mutations (Extended Data Fig. 8). Thus, epitope-focused GCs primed by DNA-VLPs can augment the maturation of bnAb precursors.

**Fig 6.**
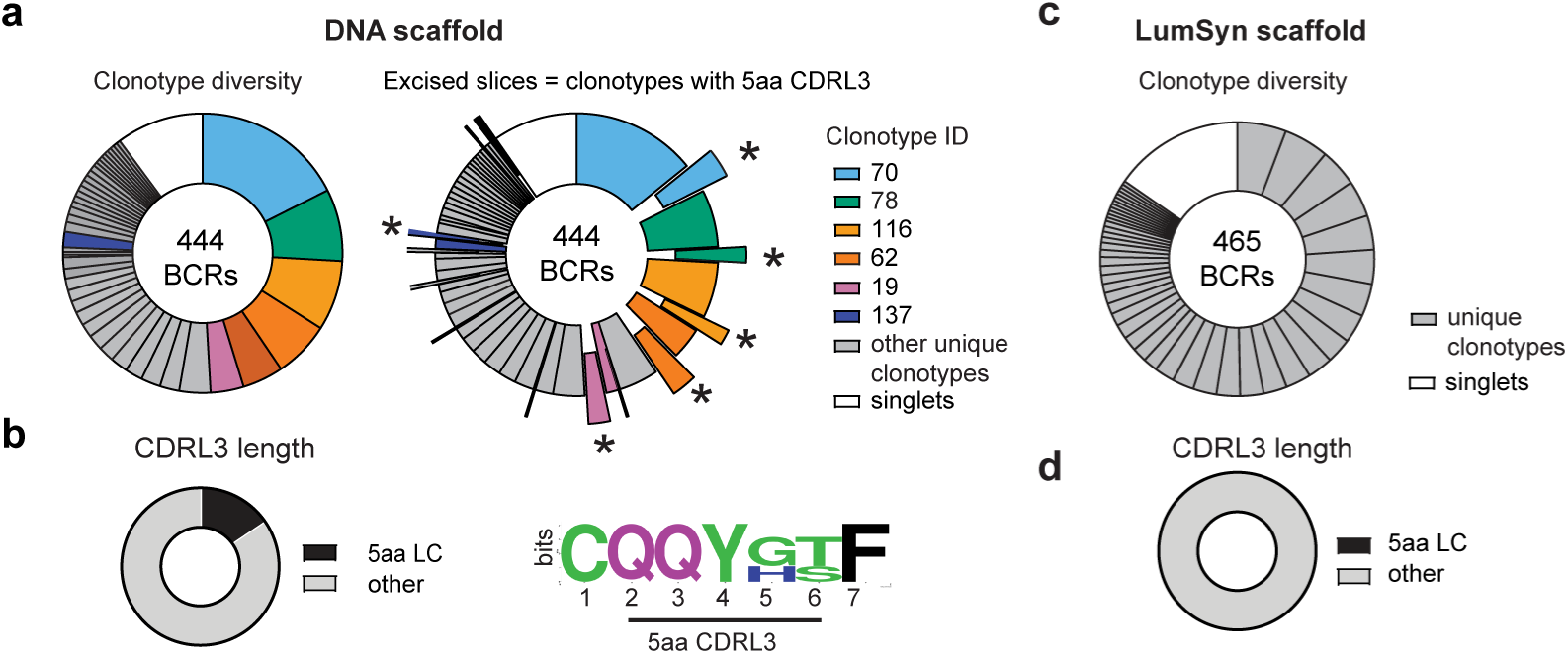
BCR sequencing reveals DNA-60mers effectively prime VRC01-class precursors in early GCs. **a,** BCR sequencing analysis of CD4bs-specific GC B cells (B220^+^ /GL7^hi^ CD38^lo^/ IgM^-^ IgG^+^/ eOD^++^ KO^-^) isolated from V_H_1-2 mice (*n*=2) immunized with d30_60mer-PADRE at day 14. (Right) B cell clonotypes were assigned based on shared CDRH3 usage. Shown in color are the top expanded clonotypes; the excised slices contain clonotypes with the short 5 amino acid CDRL3 marking the VRC01-class precursors. Asterisks represent shared public clones (shared VDJ/VJ origin) that contain the VRC01 class signature and were identically expanded in the different vaccine recipients. **b,** Distribution of light chain lengths in GC B cells from d30_60mer-PADRE immunization (right). Sequence conservation in light chains represented as a logo plot showing enrichment of VRC01-class motifs (left). **c,** Clonotype diversity in CD4bs-specific GC B cells from V_H_1-2 mice (*n*=2) immunized with p60mer. **d,** Distribution of light chain lengths in GC B cells from p60mer immunization. No 5 amino acid CDRL3 light chains were expanded by the p60mer at this early timepoint.

## Discussion

Anti-scaffold B cell responses have been previously reported for almost all protein-based nanoparticle scaffolds^17, 26, 28-30, 81, 82^, and their effects on limiting on-target immune responses and, consequently, the development of a successful vaccine against pathogens such as HIV, are still incompletely understood. Thus far, anti-scaffold antibodies have been thought to be minimally concerning for immunodominant antigens^26^, or potentially even beneficial by improving antigen capture in lymph nodes or masking scaffold epitopes in future boosts^17^. However, for subdominant antigens like HIV Env, anti-scaffold responses have been shown to dominate the serum antibody response^26^, and it has not yet been investigated how these competitor B cells influence primary GC dynamics. While strategies such as glycosylation^30-32^, PEGylation^26, 33^, and PASylation^26^ have been introduced to reduce scaffold-specific responses, such approaches are known to be imperfect. Further, even with perfect masking of exposed scaffold epitopes on assembled nanoparticles, degradation of these nanoparticles by tissue proteases that are abundant in lymph node sinuses^59^ may expose new epitopes that contribute to competitor scaffold-specific responses. Thus, our understanding of the role of protein scaffold competition on humoral immune responses remains incomplete due to limited tools and materials to test these effects of altering scaffolds without changing properties like geometry or antigen display.

Here, we instead used T-independent, inert DNA origami vaccine scaffolds to assess the impact of distracting epitopes on the potency of germinal center responses towards a displayed antigen, eOD-GT8. Notably, we found that high valency of 60 antigens and high antigen and glycan density of DNA-VLPs was critical for engaging complement pathways and promoting follicle retention and expanding antigen-specific B cells in GCs, whereas larger particles or particles with lower antigen copy numbers failed. Intriguingly, the optimal d30_60mer nanoparticle design closely mimicked the geometry of the clinical p60mer nanoparticle^29, 38^, and both particles were effectively recognized by MBL from mouse serum. However, the amount of C3 on DNA-VLPs was significantly lower compared to the protein nanoparticle, which may arise due to the porosity, surface charge, and interaction of DNA-VLPs with complement binding proteins, which may potentially limit downstream complement activation^83, 84^. This remains a design element that can be further improved for the DNA origami platform. Second, we found that augmenting the small eOD-GT8 immunogen with synthetic T cell epitopes substantially improved the immunogenicity of DNA-VLPs. Lumazine synthase and other protein scaffolds have been shown to have intrinsic T cell epitopes^74^; here we found that by adding a synthetic T cell epitope (PADRE) to the DNA-VLP, we could recapitulate scaffold-derived T cell help without introducing off-target epitopes for B cells.

Since our optimized DNA-VLP 60mer was similar to the clinical p60mer nanoparticles in regard to antigen valency, spacing, nanoparticle geometry, and presence of T cell help, we were also able to assess the effect of the scaffold material on the subsequent clonal competition in GC responses. GCs expanded by DNA-VLPs were significantly more epitope-specific than those induced by the LumSyn-based protein nanoparticle. We attribute this focusing effect due to the undiluted T cell help provided to the eOD-specific B cells, whereas in p60mer vaccination, antigen- and scaffold-specific B cells compete for LumSyn-derived Tfh help. We characterized the ratio of on-target CD4bs-specific B cells to off-target competitors (scaffold-specific B cells or non-CD4bs specific B cells) in the VH1-2 rearranging mouse model^78^ by staining lymphocytes with antigen and scaffold-only flow cytometry probes. GCs primed by DNA-VLPs were significantly more focused on the displayed antigen compared to GCs generated by p60mer, where lumazine-specific B cells competed in the GC at high frequency. Our BCR sequencing analysis confirms that epitope-focusing of the GCs enhances the priming of bnAb precursors, as evidenced by the increased number of VRC01-class precursors expanded by immune focused GCs primed with DNA-VLPs compared to the more competitive GCs primed by p60mer. Furthermore, DNA-VLPs reproducibly primed these VRC01 precursors across multiple animals. In future work, DNA-VLPs may be evaluated side by side with protein nanoparticles in heterologous boosting regimes to understand the effect of scaffold responses upon re-exposure, where feedback and imprinting mediated by pre-existing scaffold antibodies and T cell memory may be more apparent^85^.

Collectively, our results emphasize the importance of nanoscale antigen organization on lymph node delivery and illustrate that T-dependent versus -indepedent scaffold material may significantly influence clonal competition dynamics in GCs and their ability to prime desirable subdominant B cell clones, with currently unknown yet important potential consequences on limiting successful vaccine design for challenging pathogens such as HIV.

We demonstrate that DNA-VLPs are a promising, modular T-independent scaffold that is highly programmable and capable of producing potent and directed B cell responses using a clinical germline targeting immunogen, but several important issues related to potential clinical translation of are of interest for future work, such as evaluating DNA-VLP degradation due by extracellular nucleases. Additionally, while in our work we did not detect any class-switched anti-DNA or -VLP antibody responses, autoimmune risks posed by antibody responses to DNA/protein hybrids must be carefully evaluated.

## Methods

### Ethics Statement

All animal studies were performed under an institutional animal care and use committee-approved animal protocol (MIT CAC protocol # 2303000488) following local, state, and National Institutes of Health guidelines for the care and use of animals. This research does not include human participants.

### Mice

Six-to-ten-week-old female C57BL/6 mice were purchased from The Jackson Laboratory (strain no. 000664). VH1-2 mice (gift courtesy of Dr. Frederick Alt) were bred in-house and genotyped using Transnetyx. DNase I KO mice were provided by the Reizis lab at NYU Langone (original source CMMR MGI:103157) and rederived for subsequent experiments. For DNase I KO experiments, heterozygous mice were bred, pups were genotyped using PCR (Supplementary Table 2), and wildtype littermates were used as controls.

### ssDNA scaffold synthesis

Custom-length DNA scaffolds for DNA-VLPs were synthesized as previously described^86^. Briefly, SS320 *E. coli* cells were co-transformed with the corresponding plasmid and the M13cp helper plasmid (originally provided by Andrew Bradbury, Los Alamos National Laboratories). Pre-cultures of the transformed cells were grown overnight at 37 °C in 2x YT medium containing 100 µg/ml ampicillin and 15 µg/ml chloramphenicol). Cells were pelleted by centrifuging three times at 4000 g for 3 min and subsequently discarded. Phage was precipitated from the supernatant in presence of 6% (w/v) PEG8kDa and 3% (w/v) of NaCl by stirring at 4 °C for 1 h and harvested by centrifugation at 20,000 g at 4 °C for 1 h. After resuspension in TE buffer, ssDNA was extracted via the EndoFree GigaPrep purification protocol with the following modifications: Proteinase K was added to buffer P1 followed by incubation at 37 °C for 1 h, addition of buffer P2 and incubation at 70 °C for 10 min. After ssDNA purification, Triton X-144 was used to remove residual endotoxins to levels less than 0.2 EU/µmol DNA scaffold, corresponding to less than 0.000015 EU/injection into mice. Endotoxin levels were measured using ToxinSensor Gel Clot Endotoxin Assay Kits. Purity of the scaffolds was analyzed by agarose gel electrophoresis (AGE)(1.6% agarose, TAE buffer with 12 mM MgCl2, EtBr, 60 V for 150 min at room temperature).

### Oligonucleotide synthesis

DBCO-modified oligonucleotides were fabricated in-house. A (#10-1000), C (#10-1010), G (#10-1020), T (#10-1030), and DBCO-TEG (#10-1941) phosphoramidites and ancillary synthesis reagents were acquired from Glen Research and used following the manufacturer’s recommendations. Phosphoramidites were dissolved and diluted in anhydrous acetonitrile to 0.1 M prior to synthesis. Oligonucleotides were synthesized on a Biolytic Dr. Oligo 192c oligonucleotide synthesizer, at a 200 nmol scale with CPG 1000 Å standard base supports (#20-2001, #20-2011, #20-2021, #20-2031 from Glen Research) in normal mode under nitrogen. Synthesis success and yield was observed by monitoring the penultimate trityl-cleavage. Following the synthesis, oligonucleotides were cleaved and deprotected under pressure at 55°C for 2 hr in 30% ammonium hydroxide (Thermo Fisher Scientific, cat# 423305000,). The cleaved strands were desalted with acetonitrile and eluted in nuclease-free water. The DBCO-modified oligonucleotides were purified by Glen-Pak™ DNA 30 mg in a 96-Well Plate (#60-5400-01) under reduced pressure following the manufacturer’s protocols. After purification, the oligonucleotides were dried and resuspended in 1x TE and stored at 4°C. Oligonucleotide concentrations and yields were determined via UV-Vis measurements (Tecan Spark). Oligonucleotide purity was confirmed by HPLC (Waters Acquity System).

### Immunogen synthesis

eOD-GT8 monomer (with C-terminal 6x-His and cysteine), eOD-PADRE monomer (with N-terminal 6x-His and cysteine and C-terminal PADRE), and eOD-60mer were synthesized as previously reported^40^. Briefly, plasmids were transiently transfected into Expi293F cells. After 5 days of culturing in conditions described above, cell culture supernatants were collected and protein was purified in an AKTA pure chromatography system using HiTrap HP Ni sepharose affinity column, followed by size exclusion chromatography using Superdex 75 Increase 10/300 GL column (GE Healthcare). Endotoxin levels in purified protein were measured using Endosafe Nexgen-PTS system (Charles River) and were ensured to be less than 5EU/mg protein. The eOD-60mer was affinity purified by incubating with *Galanthus nivalis* lectin-conjugated agarose beads (Vector Laboratories, AL-1243-5,) overnight under gentle agitation at 4°C and eluted with lectin elution buffer containing 1 M methyl a-D-mannopyranoside (Millipore Sigma, M6882,). The resulting solution was dialyzed in PBS and further purified by size-exclusion chromatography using Superdex 200 Increase 10/300GL. Bare lumazine synthase was produced by BlueSky Bioservices as previously described^29^. Briefly, lumazine synthase from Aquifex aeolicus was expressed in E.Coli, cells were lysed, supernatant was heat-treated for 30 min at 75°C and supernatant was again clarified by centrifugation. Fully assembled particles were purified by two successive size-exclusion chromatographies, using Superdex 75 and Sephacryl 500 columns, respectively.

To add a terminal azide for SPAAC reactions, eOD-Cys monomers were reduced with 5 molar excess of TCEP, then desalted into PBS + 10 mM EDTA using Zeba 7K MWCO desalting columns (Thermo Fisher, cat# 89883). The number of free cysteines was confirmed using an Ellman’s assay. The azide linker was formed by reacting 60 mM SMCC (Thermo Fisher Scientific, cat #A35394) with 1.1x excess of Azido-PEG3-Amine (Broadpharm, cat#BP-20580) for 1 hour at room temperature. Reduced antigen was reacted with 5 molar excess of azide linker for 4 hours at room temperature, and unreacted linker was then removed by spin filter centrifugation in Amicon 10K MWCO centrifugal filters (Millipore Sigma, cat# UFC801008). Antigen was concentrated to approximately 300 μM for SPAAC reactions.

### DNA-VLP synthesis and characterization

DNA-VLPs were assembled as previously described^53^. All designs were generated using DAEDALUS^53^ (d40 VLP) or ATHENA^49^ (d30 VLP) with staples manually adjusted in Tiamat software^87^ to have outward nick positions (2 per edge). To fold origami, 30 nM of scaffold was mixed with 5–10x excess of each oligonucleotide staple in TAE buffer with 12 mM MgCl2 and thermally annealed as follows: 95°C for 5 min, 80–75°C at 1°C per 5 min, 75–30°C at 1°C per 15 min, and 30–25°C at 1°C per 10 min. DNA-VLPs were purified into PBS using Amicon Ultra 100 kDa centrifugal filters (Millipore Sigma, cat #UFC810024) spun at 2000g and stored at 4 °C. Purity and dispersity of DNA-VLPs were validated by AGE (1.6% agarose, TAE buffer with 12 mM MgCl2, SYBR Safe, 65V for 150 min) and dynamic light scattering. For antigen conjugation, DNA-VLPs were concentrated to >1 μM and reacted with at least 3 molar equivalents of eOD-Azide for 24 hours at 37°C. Excess antigen was removed by drop dialysis into 1X PBS for 16 hours (Millipore Sigma, mixed cellulose ester, 0.025 µm) and the purified nanoparticles were diluted to 100 nM and stored at 4°C. Antigen functionalization was validated using AGE (1.6% agarose, TAE buffer with 12 mM MgCl2, SYBR Safe, 65V for 150 min) and quantified using bicinchoninic acid colorimetric assay.

### Negative stain transmission electron microscopy

Uranyl formate staining was performed as previously described^28^. Briefly, DNA-VLPs were diluted to 3–5 nM in PBS and 5 μL was applied onto glow-discharged electron microscopy grids. After 30 seconds, the grids were blotted with filter paper and washed with 5 μL of fresh 2% uranyl formate solution containing 5 mM NaOH for 30 seconds. The uranyl formate solution was removed by blotting on filter paper, and the grids were stored with a desiccant before imaging. TEM was performed on an FEI Tecnai G2 Spirit Twin. Shown are images at 30K, 42K, or 67K magnifications.

### CryoEM imaging

Three microliters of the folded and purified DNA-VLP solution (approximately 300 nM) was applied onto the glow-discharged 200-mesh Quantifoil 2/1 grid, blotted for four seconds and rapidly frozen in liquid ethane using a Vitrobot Mark IV (Thermo Fisher Scientific). Grids were screened and imaged on a Talos Arctica cryo-electron microscope (Thermo Fisher Scientific) operated at 200 kV at a magnification of 79,000× (corresponding to a calibrated sampling of 1.76 Å per pixel). Micrographs were recorded by EPU software (Thermo Fisher Scientific) with a Gatan K2 Summit direct electron detector in counting mode, where each image is composed of 24 individual frames with an exposure time of 6 s and a total dose ∼63 electrons per Å2. We used a defocus range of –1.0 – –2.5 μm to collect images, which were subsequently motion-corrected using MotionCor2. Particle picking was performed manually using Relion^88^ with a particle box size of 320.

### In vitro B cell activation assays

Calcium flux assays were conducted as previously described^44^. Ramos B cells expressing germline VRC01 BCRs were stained with 10 μM Fluo-4 AM (Thermo Fisher Scientific, cat# F14217) for 30 min at 37 °C. After washing twice in serum-free RPMI medium, calcium flux assays were performed in triplicate on a Tecan plate reader at 37 °C on a 96-well microplate with 160 μl of Fluo-4 labelled Ramos cells at 2 million cells per ml. A baseline fluorescence was then recorded for 1 min, and 40 μl of antigens were added to the cells for a final concentration of 5 nM of antigen. Raw calcium traces were normalized to a common baseline by subtracting the PBS time trace at every time point, then dividing the time trace at every point by the average of the time points before antigen addition.

### Dye labeling of nanoparticles for imaging and flow probes

Protein antigens (eOD-GT8 monomers, eOD-60mer, and lumazine synthase) at 1 mg/mL were desalted into PBS and diluted 1:1 with 0.2M sodium bicarbonate buffer. Dye stocks (AF488-NHS-ester, AF647-NHS ester, AF555-NHS ester) (Thermo Fisher Scientific, cat# A20000, A20006, A20009) were prepared in DMSO, and added at 5-molar excess and reacted with antigens at 1 hour at room temperature. Due to lower efficiency of labeling lumazine synthase, reactions were performed overnight with shaking (500 rpm). Excess dye was removed by passing proteins through Zeba Spin Desalting Columns (Thermo Fisher Scientific). DNA-VLPs (d30) used as flow probes were formulated with 6 DBCO-modified oligos and reacted with 10 molar excess of AF488-Azide or AF555-Azide triethylammonium salt (Thermo Fisher Scientific, cat# A10266, A20012). Excess dye was removed by drop dialysis into 1X PBS for 16 hours (Millipore Sigma, mixed cellulose ester, 0.025 µm). For flow staining, eOD-tetramers were produced by first biotinylating Avi-tagged eOD-GT8 monomers (without PADRE) with BirA (Avidity, Inc), and mixing an excess of biotinylated eOD-GT8 monomer with fluorescently labeled Streptavidin-PE, Streptavidin-AF647, or Streptavidin-BV421 (BioLegend cat# 405203, 405237, 405226) step-wise on ice.

### Immunizations

SMNP adjuvant was prepared as previously described^54^. Antigen formulations were prepared by diluting 5 μg of the indicated antigens mixed with 5 μg of SMNP adjuvant per mouse in 1X PBS and gently mixed with a pipette. Mice (6-10 weeks old) were shaved at the tail base and immunized subcutaneously at the left and right sides of the tail base (50 μL per side). For adjuvant screening, the following formulations were used and prepared in 100 μL total volumes per mouse: SMNP (5 μg/mouse), alum in the form of Alhydrogel (InvivoGen, 100 μg/mouse), AddaVax (InvivoGen, 1:1 v/v), AS01b (GSK, 1:1 v/v), and ODN 1826 (IDT, 50 μg/mouse).

### Flow cytometry analysis of GC B and Tfh cells

Mice were euthanized at the indicated time point post-injection by CO2 asphyxiation, and inguinal lymph nodes were collected and mechanically digested into a single-cell suspension using Biomasher tubes and a motorized tissue grinder. Lymphocytes were strained twice through 60 μm Multi-Screen Mesh filter plates (Millipore Sigma, cat# MANMN6010) and washed once in 1X PBS. The cells were then stained with 1:750 dilution of Zombie UV Live Dead Stain (BioLegend, cat# 423107) diluted in PBS for 10 minutes at room temperature. Excess dye was removed by washing once with FACS buffer (1X PBS, 2% FBS, 0.01% sodium azide). The cells were then resuspended in 25 μL FACS buffer containing Fc block (BioLegend, cat# 156603) and incubated on ice for 15 minutes. Antibody mixes (2X stock) were prepared in FACS buffer and contained the following antibodies: anti-CD4 BUV737 (BD Biosciences, RM4.5), anti-B220 APC-Cy7 or PE-Cy7 (BioLegend, RA3-6B2), anti-CD38-AF488 or BV421 (BioLegend, 90), anti-GL7 PerCpCy5.5 (BioLegend, GL7), anti-CXCR5 BV605 (BioLegend, L138D7), anti-PD1-AF647 or BV421 (BioLegend, 29F.1A12), and antigen tetramer probes. For the experiment shown in Fig. 5, anti-IgM-PE-Cy7 (BioLegend, MA-69) and anti-IgD-BV785 (BioLegend, 11-26c.2a) were also included in the antibody cocktail. Tetramers were added to antibody mix so that the final amount per sample was 50 ng. For nanoparticle probes (Fig 5 & 6), nanoparticle probes were added at 5 ng/ sample. The cells in Fc block were mixed with antibody and probe mixture and incubated for 30 minutes on ice, then washed twice with FACS buffer. The cells were fixed with 2% PFA, washed once and stored in FACS buffer until flow cytometry analysis. The cells were transferred to U-bottom 96-well plates and mixed with CountBright Absolute Counting beads (Thermo Fisher Scientific, cat# C36950) before analysis. Flow cytometry was carried out on a BD LSR Fortessa or Symphony A3 cytometer in plate-mode. Full-minus-one (FMO) staining controls were included in every experiment for drawing gates.

### Lymph node microscopy

For lymph node histology, mice were immunized with 5 μg of fluorescently labeled eOD-GT8 monomer, DNA-VLPs, or protein nanoparticles and sacrificed at the indicated time points. Mice were euthanized by CO2 asphyxiation, and inguinal lymph nodes were flash frozen in OCT embedding medium and sectioned on a Leica Cryostat (10 μm thick sections) and stored at – 80°C. For immunostaining, the sections were quickly thawed, fixed with 10% neutral buffered formalin, and blocked/permeabilized with PBS containing 2% BSA and 0.01% Triton-X. The sections were then stained with 1:100 dilutions of anti-CD35-BV421 (BD Biosciences, clone 8c12), anti-CD169-AF488 (BioLegend clone 3D6.112), or anti-F4/80 (BioLegend clone BM8) in block/perm buffer for 1 hour in a humidity chamber. The slides were washed three times in PBS and mounted with ProLong Diamond Antifade mountant (Thermo Fisher Scientific) and secured with sealant. Slides were stored in the dark and imaged on a Leica Sp8 Laser Scanning Confocal Microscope with 25x water-based objective. Colocalization between channels was quantified using a MATLAB script which applied a Gaussian filter to each channel, created a binary mask to identify FDC or SSMs, and calculated the fraction of area occupied by antigen signal. In ImageJ, LUT tables were selected to optimize contrast and enhance readability for color-blindness.

For cleared lymph node imaging, mice were immunized with 5 or 10 μg of fluorescently labeled nanoparticles (specified in captions). For in situ follicle labeling, 4 μg of anti-CD35-BV421 (BD, clone 8c12) was injected subcutaneously 12–16 hours prior to tissue harvesting. Inguinal lymph nodes were isolated and fixed in 4% PFA for 24 hours and cleared using DISCO as previously described^61^. Briefly, LNs were washed twice in PBS and excess fat and connective tissue were removed. Nodes were then gradually moved into solutions containing successively higher concentrations of methanol until they were incubated for half an hour in pure methanol. Nodes were then bleached in hydrogen peroxide for one minute before being returned to methanol for half an hour. They were then gradually transferred into solutions containing increasing concentrations of tertiary-butanol before eventually being incubated in pure tertiary-butanol 0.4% α-tocopherol for one hour. Nodes were then removed from solution and allowed to dry completely before being placed in dichloromethane. After the lymph nodes dropped to the bottom of tubes following swirling, they were stored in dibenzyl ether with 0.4% α-tocopherol. Follicles were imaged using an Olympus FV1200 Laser Scanning Confocal Microscope at 10x magnification over a 300 μm distance. Lasers were set to minimize pixel saturation in the brightest samples. Images were analyzed using ImageJ software. LUT tables were selected to optimize contrast and enhance interpretability for color-blindness.

### Passivation of DNA-VLPs with polylysine-PEG

DNA-VLPs (d40_30mer) were coated with polylysine-PEG1K or 5K (Alamanda Polymers) at 1:1 N:P ratio, as previously described^65^, and complexes were analyzed by AGE (1.6% agarose, TAE buffer with 12 mM MgCl2, SYBR Safe, 60V for 150 min at room temperature). For serum stability analysis, d40_30mer was incubated with or without polymer coating at 37°C in the presence of 10% mouse serum (freshly isolated from C57BL/6 mice) and analyzed by AGE as described previously. For macrophage association, RAW264.7 murine macrophages were seeded into 24-well plates and incubated with 10 nM AF647-labeled d40_30mer with or without PEG coating. After 2 hrs of incubation, the cells were thoroughly washed, and nanoparticle signal was analyzed by flow cytometry. Lymph node imaging and measurement of GC responses induced by vaccination with uncoated or coated DNA-VLPs were performed as described above.

### ELISA analysis of serum antibody responses

Blood samples were collected from immunized mice via retro-orbital or submandibular bleeds and serum was isolated using centrifugation through Serum Gel tubes (Sarstedt) and stored at –20°C. MaxiSorp plates (Thermo Fisher Scientific) were coated with 2 µg/ml protein immunogens (eOD-GT8 monomer, eOD-CD4bs-KO monomer, or Lumazine Synthase) overnight at 4°C. For anti-dsDNA and anti-VLP ELISAs, ELISA plates were first coated first with 100 ug/mL poly-D-lysine in 1X PBS and incubated at 37°C for 1 hr. The plates were washed once with PBS and then incubated with 10 μg/mL of bare DNA-VLPs (d30) or calf-thymus DNA (Sigma Aldrich) overnight at 4°C. The plates were blocked with Casein block buffer (G-Biosciences). Plates were washed four times in 1x PBS containing 0.2% Tween-20, and dilutions of serum in blocking buffer were added and incubated for two hours. A commercial dsDNA antibody (Abcam, clone 35I9) was used as a positive control in anti-DNA assays, and murine VRC01 was included in all anti-eOD-GT8 ELISAs. Plates were washed four times and an HRP-conjugated goat anti-mouse IgG (BioRad) was added and incubated for one hour. Plates were then washed and TMB was added. The reaction was stopped with sulfuric acid once the wells containing the lowest dilutions of TMB began to develop visually and the absorbance of each well was determined. All titers reported are inverse dilutions where A450nm – A540nm (reference wavelength) equals 0.2 or as area under the curve measurements (specified in figure captions). Anti-DNA IgM ELISAs were performed using a commercial assay (Alpha Diagnostic International, cat# 5130) following manufacturer instructions.

### C3 and MBL ELISAs

ELISA plates were first coated with 100 μg/mL poly-D-lysine in 1X PBS and incubated at 37°C for 1 hr. The plates were washed with DPBS, then coated with nanoparticles overnight at 4°C (DNA-VLP concentrations were normalized so that final eOD concentration was 1 μg/mL). The plates were blocked with 1X PBS containing calcium and magnesium with 1% BSA, then washed 4 times with 1X PBS containing 0.2% Tween-20. Fresh mouse serum was collected from naive C57BL/6 mice in Sarstedt serum gel tubes and kept on ice. Serum was diluted in blocking buffer (starting at 30% v/v) and further diluted in 1:2 dilution series. Serum dilutions were transferred to ELISA plates and incubated for 1 hr at 37°C. For recombinant mouse MBL assays, MBL2 (Biotechne R&D, cat #2208) was diluted in blocking buffer starting at 100 ug/mL concentration and diluted 1:2. The plates were washed 4 times as before, then incubated with 2 μg/mL rat anti-mouse MBL antibody (Abcam, clone 14D12) or rat anti-mouse C3 antibody (Abcam, clone 11H9) diluted in blocking buffer. The plates were washed four times, then incubated with 1:5000 HRP-conjugated mouse-adsorbed goat anti-rat IgG (Bio-Rad, STAR72). Plates were then washed and TMB was added. The reaction was stopped with sulfuric acid once the wells containing the lowest dilutions of TMB began to develop visually and the absorbance of each well was determined. Complement deposition is reported as an area under the curve.

### Single cell BCR sequencing

CD4bs-specific GC B cells were FACS-sorted from inguinal lymph nodes harvested from VH1-2 mice immunized with protein or DNA-scaffolded immunogens, gated on B220^+^/CD38^lo^GL7^hi^/IgM^-^ IgG^+^/eOD^++^KO^-^ populations. We generated BCR libraries from whole transcriptome amplification (WTA) products produced using the Smart-Seq2 protocol on the FACS isolated B cells^89^. The WTA products underwent two 0.8x (v/v) SPRI bead-based cleanups and were verified using High Sensitivity D5000 ScreenTape (Agilent Technologies Inc) and quantified and normalized using the Qubit dsDNA HS Assay kit (Thermo Fisher Scientific, cat # Q32854). To enrich the BCR sequences (FR1 to CDR3), corresponding heavy and light chains were amplified (HotStarTaq Plus Master Kit, Qiagen, cat #203645) with a pool of partially degenerate primers specific against all possible IGHV (human) or IGLV (mouse) and IGKV (mouse) segments in the FR1 region (final concentration: 10 µM each) and reverse primers against the heavy or light constant regions (final concentration: 10 µM each)^89^. These primers were also built with attachments to the Illumina P7 (V region) and P5 (constant region) sequences^89^. The amplicons were quantified and normalized following BCR amplification and SPRI cleanup, after which cellular barcodes and Illumina sequencing adapters (Nextera XT Index Adapters, Illumina Inc.) were added to each amplified heavy and light chain using step-out PCR (Kapa HiFi HotStart ReadyMix; Fisher Scientific cat # 50-196-5217). After another SPRI cleanup the HC and LC samples were pooled and the single-cell BCR libraries were sequenced via paired-end 250x250 reads and 8x8 index reads on an Illumina MiSeq System (MiSeq Reagent Kit v2 (500-cycle), cat# MS-102-2003). The BCR heavy and light chains reads were then paired using the barcodes and the overlapping sequencing reads were reconstructed with PandaSeq^90^, and aligned against the human or mouse IMGT database^91^. Sequencing error correction was performed with MigMAP, a wrapper for IgBlast (https://github.com/mikessh/migmap). Consensus VH and VL/VK chain for each single cell was achieved by collating all reads with the same CDR3 sequence and then calling the top heavy and light chain sequences by frequency. The nucleotide sequences (full-length heavy chain sequences) from the public BCRs bearing the short five amino acid CDRL3 were aligned with unmutated germline (IGHV1-2*02) sequence using the ClustalW alignment tool in the MEGAx^92^. The phylogenetic trees were constructed using the neighbor-joining method with Poisson Model in the MEGAx^93^. The reliability of the tree nodes was tested using the Felsenstein bootstrap method with 500 replicates.

### Statistics

Statistics were computed in GraphPad Prism v.9 as denoted in the figure captions. For flow cytometry analyses, comparisons between two groups were made using a two-sided Student’s t-test or Mann Whitney U test if the sample size was too small to pass a normality test. For experiments with comparison with 3 or more groups, we used one-way analysis of variance (ANOVA) with host-hoc Tukey test for multiple comparisons. For data shown on log axes, geometric means and geometric S.D. values are shown. No outliers were excluded. One mouse from Fig.4 was excluded due to accidental pregnancy. Exact *P* values are denoted in the figures. For all figures, NS is not significant (*P* > 0.05).

## Data and code availability

All data generated or analyzed during this study are included in this article (and its supplementary information files). Source data are provided with this paper. The raw imaging data are available from the corresponding author upon request. BCR sequences identified in this study will be deposited in GenBank prior to publication (accession numbers, TBD). No new code was developed for this study.

## Acknowledgements

This work was supported by National Institute of Health grant R01-AI162307-03 (M.B., D.J.I.), R01AI155447, P30AI060354, R01-AI153098 (D.L.), the Ragon Institute of Mass General Brigham, MIT, and Harvard (D.J.I, D.L.), the Howard Hughes Medical Institute (D.J.I.), National Science Foundation Fellowship 4000168384 (A.R.), and National Science Foundation Fellowship 4000189657 (G.A.K.), and Novo Nordisk Foundation NNF23OC0082848 (M.O.). This work was supported in part by the Koch Institute Support (core) Grant P30-CA14051 from the National Cancer Institute and by the National Institute of Environmental Health Sciences of the NIH under award P30-ES002109.

We thank the Robert A. Swanson (1969) Biotechnology Center at the Koch Institute for technical support, specifically the Flow Cytometry and Microscopy Facilities. We acknowledge MIT.nano for the use of core characterization facilities and support from a core center grant, P30-ES002109. We thank Xiao Wang for assistance in CryoEM grid preparation and imaging. We are grateful to Benjamin Clancy for production of ssDNA scaffolds used in the study. We thank the Ragon Institute Flow Cytometry Core for assistance in FACS sorting. We are grateful to Fred Alt (source of VH1-2 mice) and Boris Reizis (source of DNase I KO mice, originally from CMMR).

## Author Contributions

A.R., G.A.K, M.B., and D.J.I conceived the study. A.R designed, folded, and characterized the nanoparticles in this study that have not previously described (d40_60mer, d30_30mer, d30_60mer). G.A.K. synthesized all DBCO-modified oligos used in this study. H.S. produced eOD immunogens in the study. T.S. provided lumazine synthase nanoparticles. M.O. performed transmission electron microscopy. A.R. and A.P.C. maintained all animal cohorts, and A.R. performed immunizations. A.R., A.P.C., M.O., V.R.L, and K.S. conducted animal procedures and tissue processing. A.R. conducted flow cytometry and microscopy experiments. J.C. and B.R. developed methods for DNase genotyping and provided support on DNase I KO experiments. L.R. and A.R. performed FACS sorting on antigen-specific B cells, and L.R. conducted whole transcriptome amplification, Illumina sequencing, and all subsequent analysis of BCR sequencing data. A.R., G.A.K., L.R., A.P.C., D.L., M.B., and D.J.I interpreted the results. A.R., M.B., and D.J.I. wrote the manuscript. All authors read and reviewed the manuscript.

## Supplementary Information is available for this paper

**Correspondence and requests for materials should be addressed to Darrell Irvine and Mark Bathe.**

## Extended Data Figures

**Extended Data Fig. 1.**
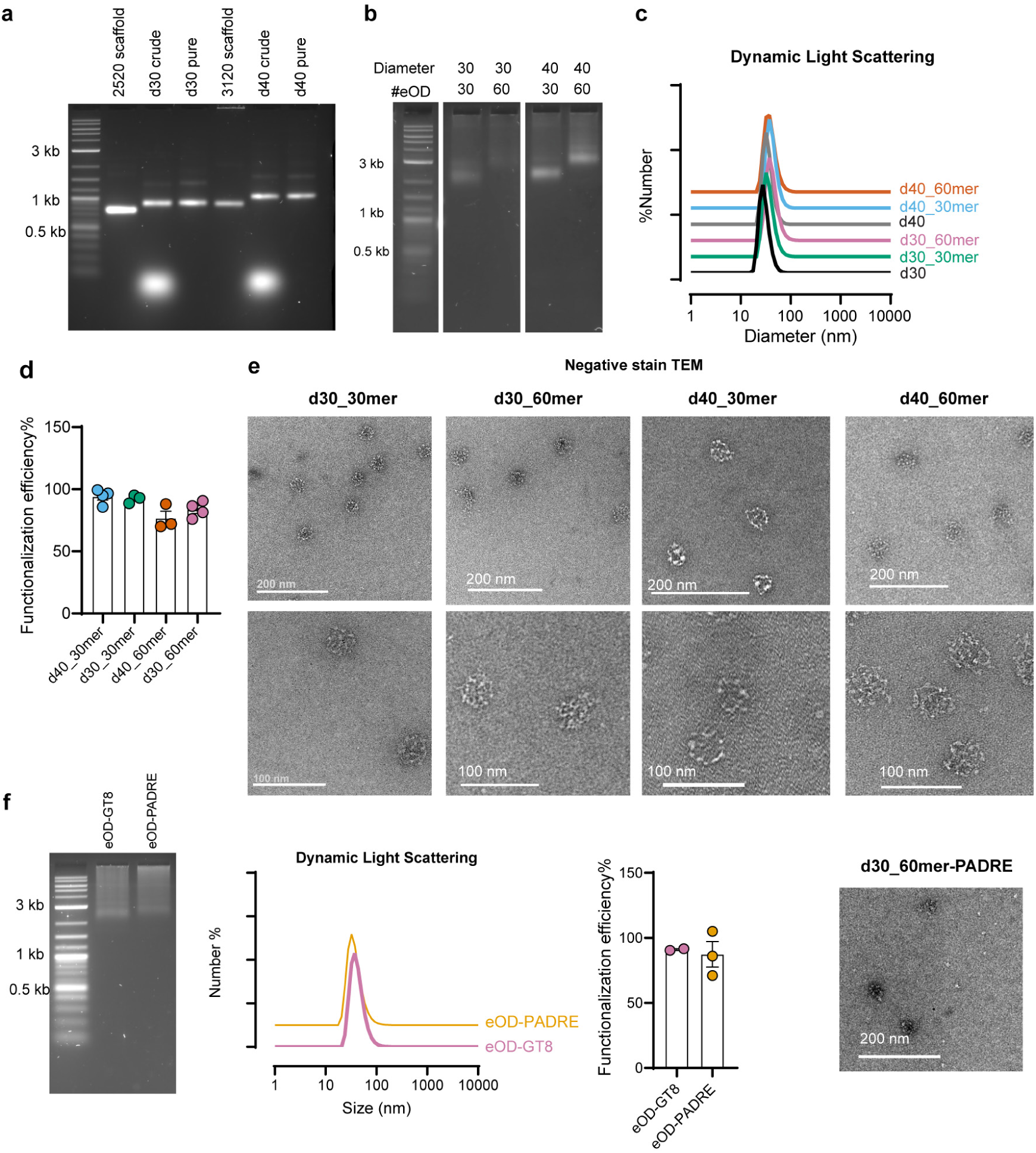
Characterization of DNA-VLPs designed in this study. **a,** Agarose gel electrophoresis (AGE) of ssDNA scaffolds, crude folded VLPs, and purified VLPs (after 3 rounds of Amicon filter centrifugation to remove excess staple oligos). Samples were run on a 1.6% agarose gel with SYBRSafe stain. **b,** AGE of DNA-VLPs following SPAAC reactions with eOD-Azide and purification with drop dialysis. **c,** Dynamic light scattering of DNA-VLPs pre-and post-antigen reaction. Shown are representative histograms, where x axis represents hydrodynamic diameter (nm) and y axis is the frequency by number. Each histogram trace is offset vertically for ease of visualization. **d,** Antigen functionalization efficiency from separate antigen preparations calculated from BCA assays. **e**, Negative stain transmission electron microscopy of each DNA-VLP design made in the study. Scale bars = 100 nm. **f**, Characterization of d30_60mer reacted with eOD-GT8 or eOD-PADE. Shown are representative AGE, dynamic light scattering, antigen functionalization efficiency, and negative stain TEM.

**Extended Data Fig. 2.**
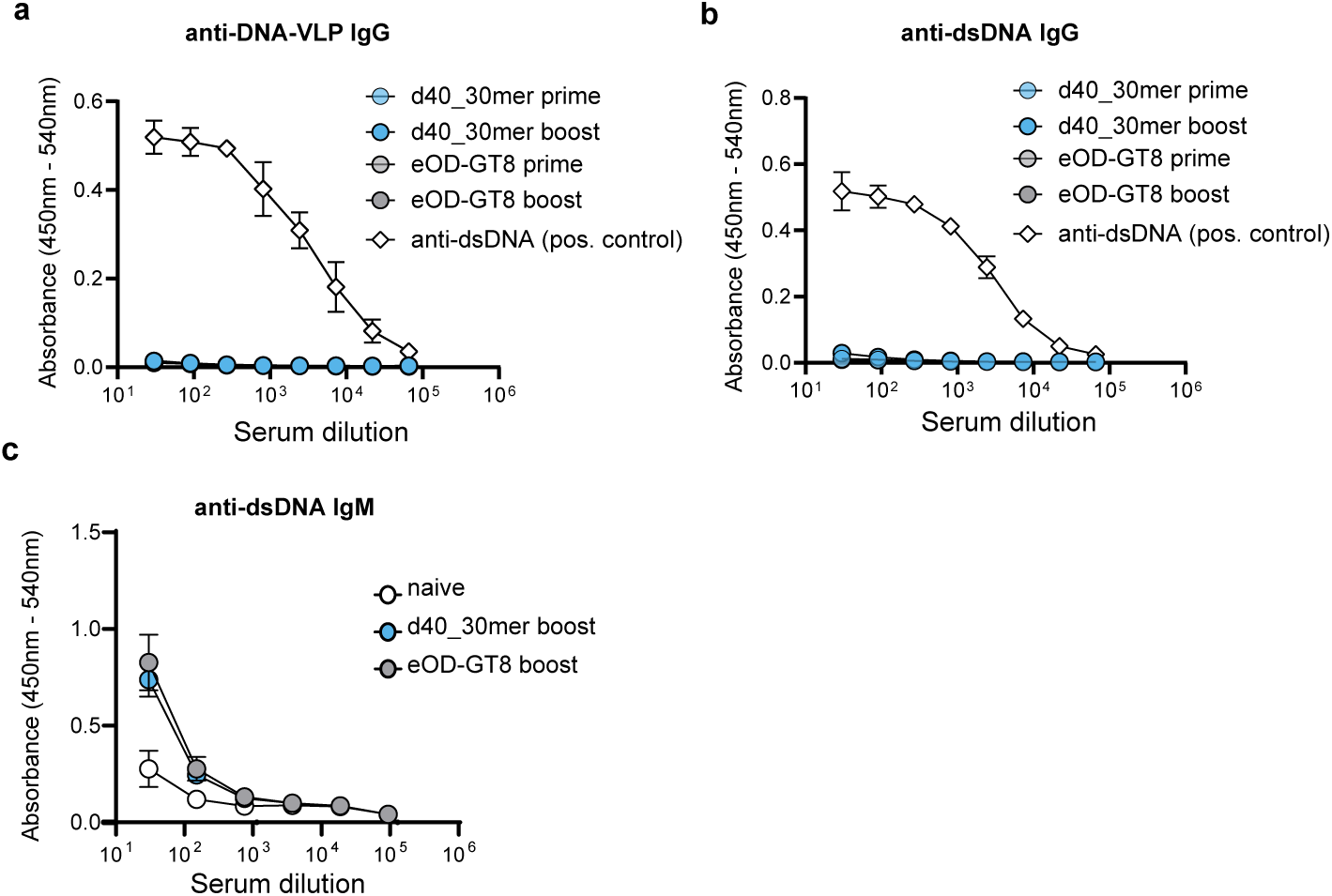
Analysis of anti-VLP and anti-dsDNA serum antibody responses, related to Figure 1. **a**, Raw serum dilutions curves of anti-origami ELISA shown in Fig 1H. Bare DNA-VLPs (d40) were used to coat ELISA plates, followed by incubations with serum from mice prime and/or boosted with eOD monomer or d40_30mer. Mouse anti-dsDNA IgG (Abcam 35I9) was used as a positive control. **b,** Anti-dsDNA ELISAs were measured by coating calf-thymus DNA on high binding ELISA plates and incubating with mouse serum samples from immunization studies. Shown are raw serum dilution curves (left) and area under the curve measurements (right). **c,** Anti-dsDNA IgM responses induced after vaccination with eOD-GT8 or d40_30mer. A commercial mouse anti-dsDNA ELISA kit was purchased from Alpha Diagnostics (cat# 5130) and used to assay IgM against dsDNA.

**Extended Data Fig. 3.**
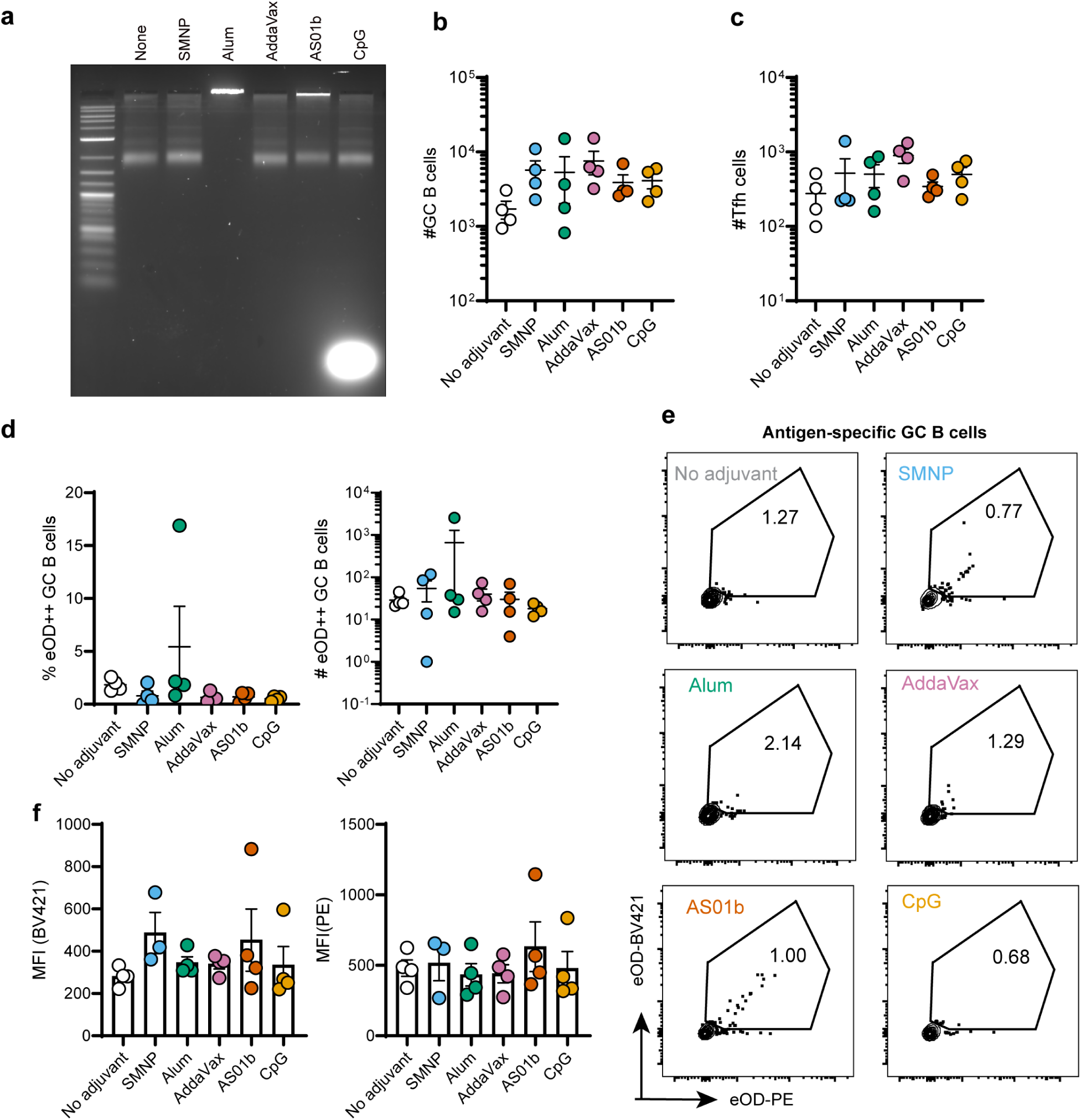
Comparison of GC responses induced by d40_30mer with common adjuvants. **a,** d40_30mer was formulated with different classes of adjuvants and gently mixed. Samples were loaded onto an agarose gel ten minutes after addition of adjuvant. Significant aggregation was observed for alum co-formulation and some minor aggregates are detected in the mixture with AS01b. **b,** Mice were primed with d40_30mer with different adjuvants and sacrificed at day 14. Inguinal lymph nodes were isolated and processed into a single cell suspension, then analyzed by flow cytometry. **c,** GC B cells (B220^+^ GL7 ^hi^, CD38 ^lo^), **d-g,** antigen-specific GC B cells, and follicular helper T cells (CD4^+^ CXCR5^hi^, PD1^hi^). No statistically significant difference was observed in cellular response between groups, as measured by one-way ELISA with post-hoc Tukey multiple comparison test. One mouse immunized with eOD-60mer (black) was used as validation for tetramer staining.

**Extended Data Fig. 4.**
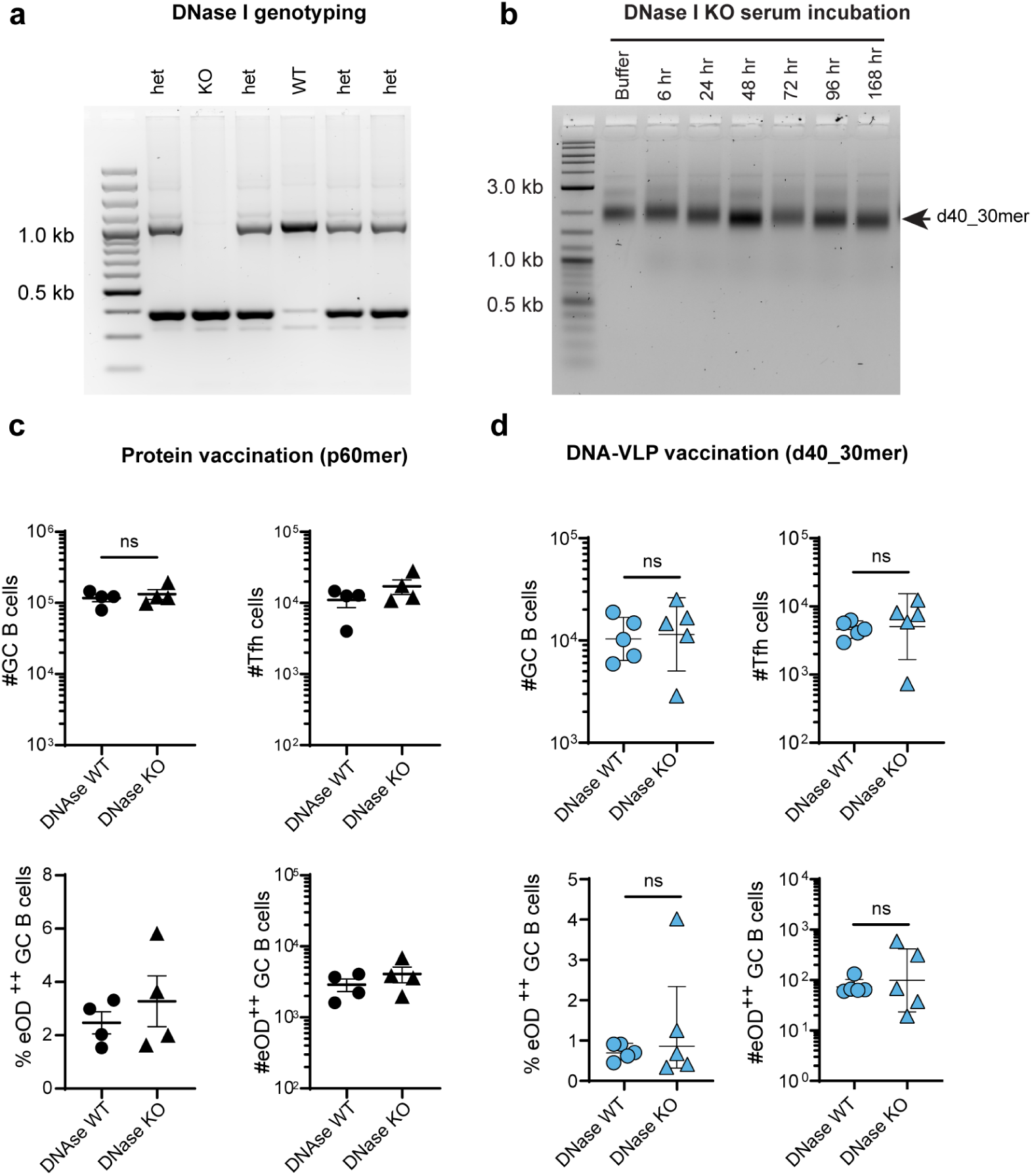
DNase I KO mice offer enhanced serum stability for DNA-VLPs but do not lead to enhanced GCs by d40_30mer vaccination. **a,** DNase I KO mice were genotyped using direct PCR from ear biopsies. Shown is a representative genotyping assay (homozygous KO with bands ∼400 bp, WT with bands at ∼1000bp, and heterozygous mice displaying both bands). **b,** Serum stability of d40_30mer incubated DNase I KO serum for up to 7 days (10% serum in PBS with calcium and magnesium). Shown are agarose gels with DNA-VLPs loaded at different time points. DNA bands do not degrade in the presence of DNase I KO serum. The slight shift downwards in the bands over time reflects the degradation of eOD-GT8 by serum proteases. **c**, WT or DNase I KO mice (n=4 mice/group) were immunized subcutaneously with 5 μg of p60mer with SMNP adjuvant. GC responses in inguinal lymph nodes were analyzed by flow cytometry on day 14. DNase I KO mice launch equally strong primary GCs in response to protein vaccination compared to WT mice. d, WT or DNase I KO mice (n=4 mice/group) were immunized subcutaneously with 5 μg of d40_30mer with SMNP adjuvant. GC responses in inguinal lymph nodes were analyzed by flow cytometry on day 14. D40_30mer primed similarly low responses in WT vs. DNase KO mice.

**Extended Data Fig. 5.**
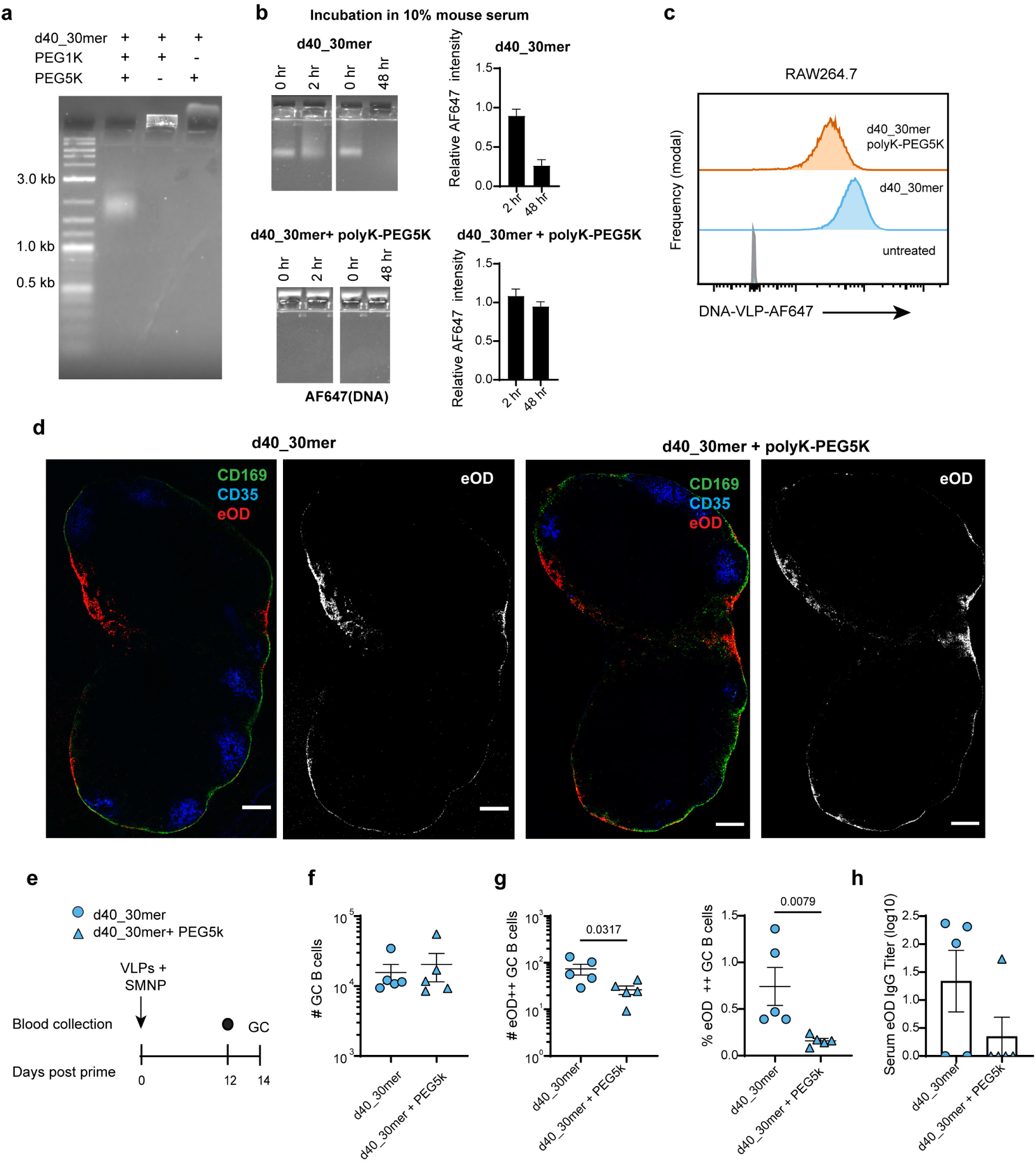
Polylysine-PEG coating on d40_30mer dampens GC responses. **a**, d40_30mer was coated with polylysine-PEG1K or 5K (Alamanda polymers) at 1:1 N:P ratio, as previously described by Ponnuswamy, et. al., and complexes were analyzed by AGE. Coated nanoparticles did not migrate through the gel due to neutralized charge. Significant aggregation was observed for the polylysine-PEG1K coating, so for subsequent studies only the PEG5K polymer was used. **b**, d40_30mer was formulated with 18x-AF647 dyes and 30 eOD-GT8 antigen and coated with polylysine-PEG5K. Bare and PEGylated VLPs were incubated with 10% fresh mouse serum (from naïve C57BL/6 mice) at 37°C for either 2 or 48 hours prior to analysis on AGE. Representative gels are shown (left) and relative AF647 band intensities (right) for bare and PEGylated particles. **c,** AF647-labeled d40_30mer with or without polylysine-PEG5K were incubated with RAW264.7 macrophages for 1 hour and analyzed by flow cytometry. **d,** d40_30mer was formulated with AF647-labeled eOD-GT8 with and without polylysine-PEG5K and injected subcutaneously into C57BL/6 mice. Inguinal lymph nodes were flash frozen at 2 hours, cryosectioned, and stained with anti-CD35 (blue) and anti-CD169 (green). Representative nodes are shown (left-composite of three channels, right-greyscale image of nanoparticle distribution only). **e,** Immunization scheme for GC analysis. Mice were immunized with d40_30mer or d40_30mer coated with polylysine-PEG5k (5 μg eOD dose with 5 μg SMNP), and GCs were analyzed by flow cytometry at day 14**. f,** Total counts of GC B cells (B220+ GL7 hi CD38 lo). **g**, Total counts and frequency of antigen-specific GC B cells. Significance was determined by Mann Whitney U-test. **h**, anti-eOD IgG titer on day 14.

**Extended Data Fig 6.**
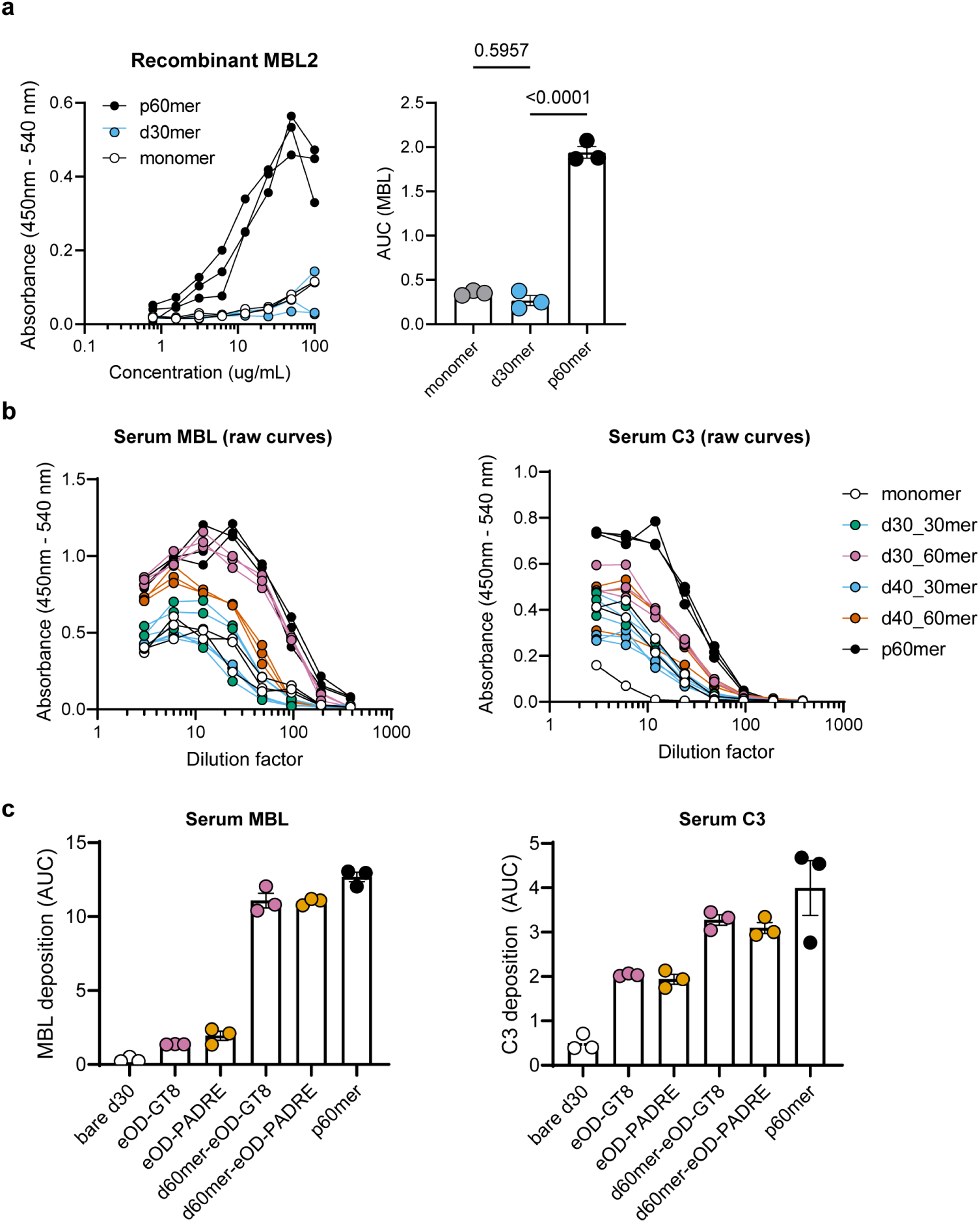
Complement deposition on DNA-VLPs. **(A)** Binding of recombinant MBL2 to antigen formulations. Antigen formulations were immobilized on ELISA plates and incubated with dilutions of recombinant mouse MBL2 (R&D/Biotechne). MBL was detected using rat anti-mouse MBL2 (Abcam 14D12) and goat anti-rat HRP conjugate. Monomer and d40_30mer had minimal deposition of MBL2, while p60mer robustly bound MBL. Shown are raw dilution curves and A.U.C measurements. (B) Raw serum dilution curves for C3 and MBL ELISAs. Antigen formulations were immobilized on ELISA plates and incubated with dilutions of fresh mouse serum (each replicate is serum pooled from 3 mice). C3 was detected with rat anti-mouse C3 (Abcam 11H9) and goat anti-rat HRP secondary. MBL was detected using rat anti-mouse MBL2 (Abcam 14D12) and goat anti-rat HRP conjugate. Shown are raw dilution curves, and corresponding AUC measurements are shown in Figure 3 in the main text. (C) MBL (left) and C3 (right) deposition ELISAs using monomer or d30_60mers formulated with eOD-GT8 or eOD-PADRE. Shown are AUC measurements. P60mer is shown as an assay control.

**Extended Data Fig. 7.**
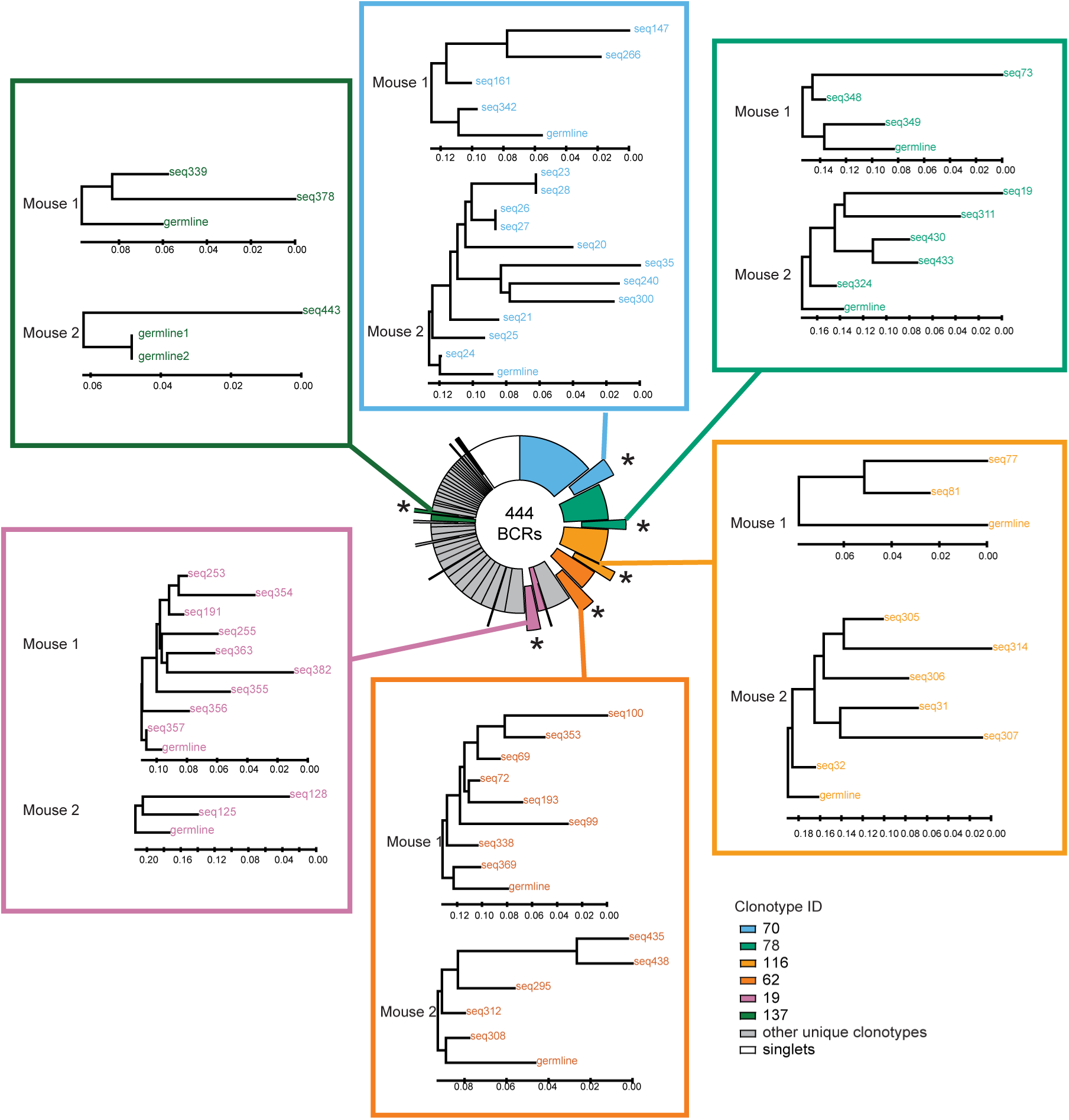
Reproducible expansion of public VRC01-class B cell clones by DNA-VLP. Phylogenetic trees of public B cell clonal lineages (shared VDJ/VJ origin within the different vaccine recipients) containing the 5 aa CDRL3 signature were constructed for each mouse. The unmutated IGHV1-2*02 nucleotide sequence was used as the reference sequence and the trees reflect mutational diversification of the HC. DNA-VLP expanded six public VRC01-class clones, presented here (Clone ID - 70, 78, 116, 62, 19 and 137). Bootstrap probability (>65%, 500 replicates) was used at the tree nodes to test reliability and the scale bar represents the nucleotide distance at an interval of 0.02 nucleotides per position in the sequence.

**Extended Data Fig. 8.**
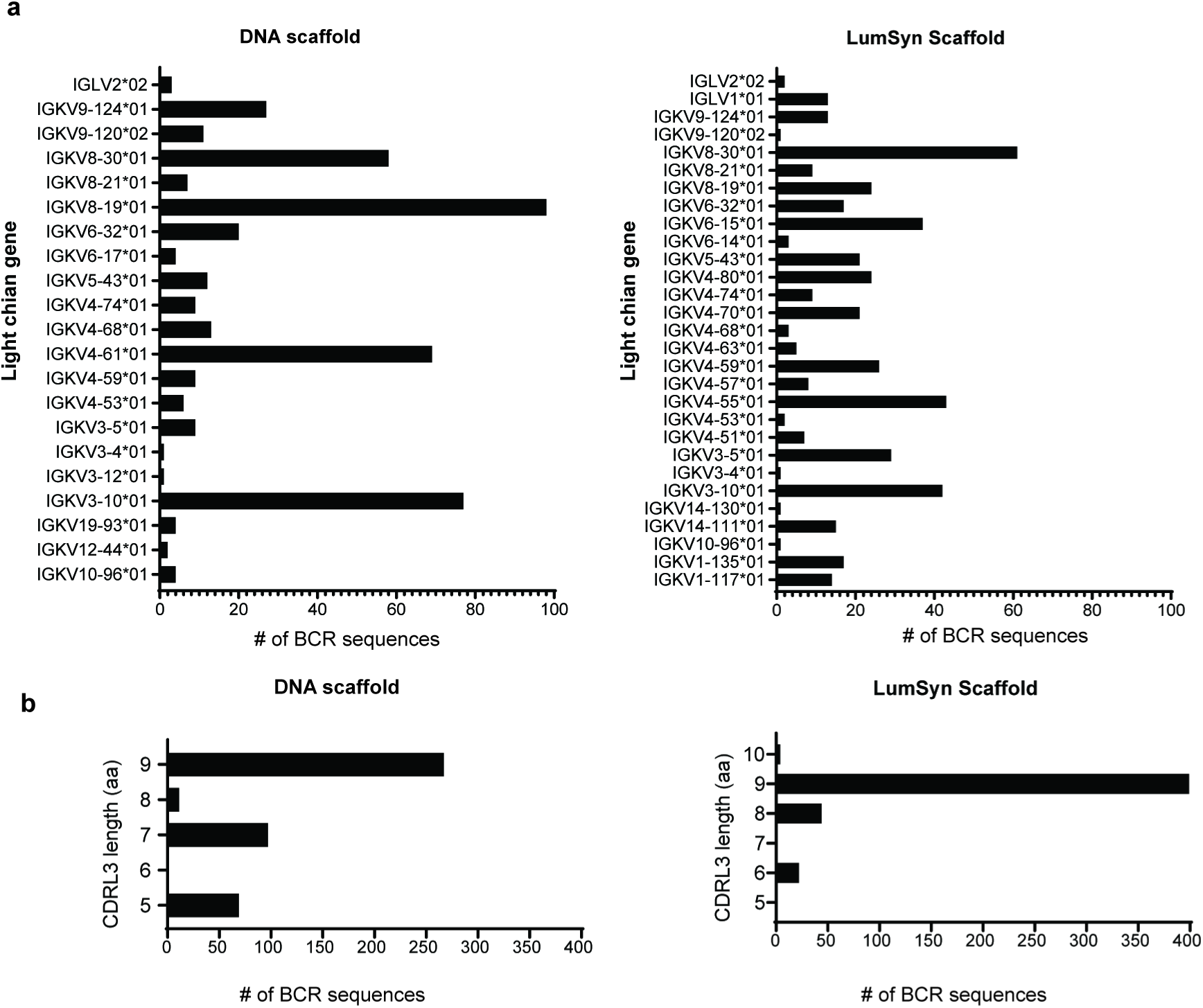
Mouse light chain gene usage and CDRL3 lengths within the eOD specific BCRs elicited by DNA-VLP or p60mer. **a,** Mouse IGKV or IGLV gene usage in the eOD specific BCRs expanded in the GC by each immunogen (n=444 BCRs for VLP-DNA; n=469 BCRs for p60mer, n=2 mice per immunogen). **b,** Corresponding CDRL3 lengths of these antigen specific BCRs. In the case of the 5 aa CDRL3 signature expanded by VLP-DNA, the LC gene used was exclusively IGKV4-61*01.

**Extended Fig. 9.**
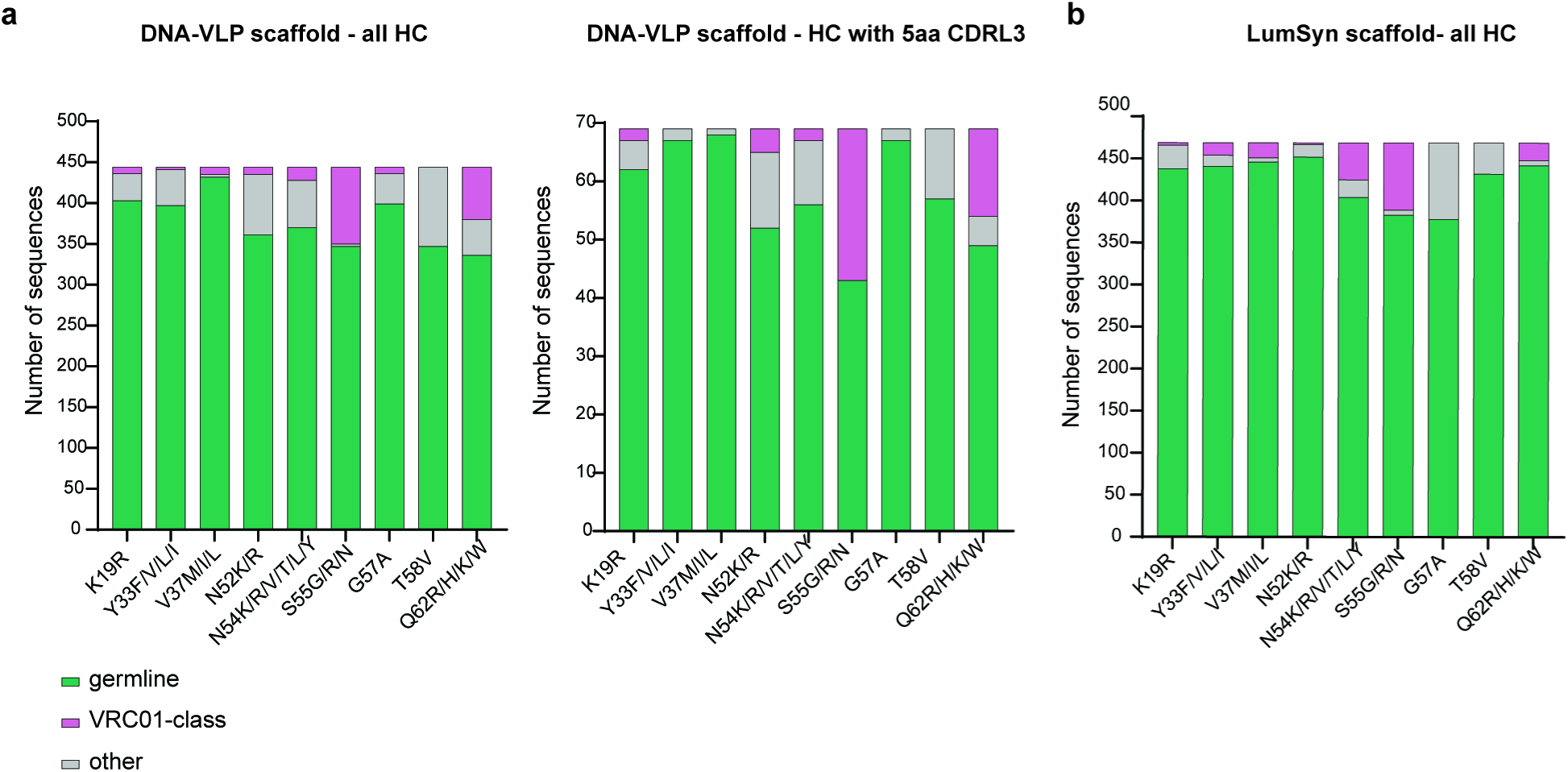
VRC01 mutations within the eOD-specific GC BCRs elicited by DNA-VLP or p60mer. **a,** stacked bar graph (left) denotes the quantity of VRC01-class mutations in the IGHV1-2*02 HC for all CD4bs-specific BCRs expanded by DNA-VLP (n=444 BCRs for VLP-DNA, n=2 mice). The adjacent graph (right) shows this for only BCRs bearing the 5 aa CDRL3 VRC01 signature. **b**, Stacked bar graph showing the quantity of VRC01-class mutations in CD4bs-specific BCRs expanded by p60mer n=469 BCRs, n=2 mice).

## References

1. Haynes, B.F. et al. Strategies for HIV-1 vaccines that induce broadly neutralizing antibodies. Nature Reviews Immunology 23, 142–158 (2023).

2. Guthmiller, J.J. et al. Broadly neutralizing antibodies target a haemagglutinin anchor epitope. Nature 602, 314–320 (2022).

3. Paules, C.I., Marston, H.D., Eisinger, R.W., Baltimore, D. & Fauci, A.S. The Pathway to a Universal Influenza Vaccine. Immunity 47, 599–603 (2017).

4. Chen, Y. et al. Broadly neutralizing antibodies to SARS-CoV-2 and other human coronaviruses. Nature Reviews Immunology 23, 189–199 (2023).

5. Durham, N.D. et al. Broadly neutralizing human antibodies against dengue virus identified by single B cell transcriptomics. eLife 8, e52384 (2019).

6. Abbott, R.K. et al. Precursor Frequency and Affinity Determine B Cell Competitive Fitness in Germinal Centers, Tested with Germline-Targeting HIV Vaccine Immunogens. Immunity 48, 133–146.e136 (2018).

7. Ng’uni, T., Chasara, C. & Ndhlovu, Z.M. Major Scientific Hurdles in HIV Vaccine Development: Historical Perspective and Future Directions. Front. Immunol. 11, 590780 (2020).

8. McGuire, A.T. et al. Engineering HIV envelope protein to activate germline B cell receptors of broadly neutralizing anti-CD4 binding site antibodies. Journal of Experimental Medicine 210, 655–663 (2013).

9. Victora, G.D. & Nussenzweig, M.C. Germinal Centers. Annu. Rev. Immunol. 40, 413–442 (2022).

10. Steichen, J.M. et al. A generalized HIV vaccine design strategy for priming of broadly neutralizing antibody responses. Science 366 (2019).

11. Schiller, J.T., Castellsagué, X. & Garland, S.M. A Review of Clinical Trials of Human Papillomavirus Prophylactic Vaccines. Vaccine 30, F123–F138 (2012).

12. Mohsen, M.O. & Bachmann, M.F. Virus-like particle vaccinology, from bench to bedside. Cellular & Molecular Immunology 19, 993–1011 (2022).

13. Cohen, A.A. et al. Mosaic nanoparticles elicit cross-reactive immune responses to zoonotic coronaviruses in mice. Science 371, 735–741 (2021).

14. Brune, K.D. et al. Plug-and-Display: decoration of Virus-Like Particles via isopeptide bonds for modular immunization. Sci Rep (2016).

15. Ols, S. et al. Multivalent antigen display on nanoparticle immunogens increases B cell clonotype diversity and neutralization breadth to pneumoviruses. Immunity 56, 2425–2441.e2414 (2023).

16. Song, J.Y. et al. Safety and immunogenicity of a SARS-CoV-2 recombinant protein nanoparticle vaccine (GBP510) adjuvanted with AS03: A randomised, placebo-controlled, observer-blinded phase 1/2 trial. eClinicalMedicine 51 (2022).

17. Kanekiyo, M. et al. Self-assembling influenza nanoparticle vaccines elicit broadly neutralizing H1N1 antibodies. Nature 499, 102–106 (2013).

18. Singh, A. Eliciting B cell immunity against infectious diseases using nanovaccines. Nature Nanotechnology 16, 16–24 (2021).

19. Marcandalli, J. et al. Induction of Potent Neutralizing Antibody Responses by a Designed Protein Nanoparticle Vaccine for Respiratory Syncytial Virus. Cell 176 (2019).

20. Bachmann, M.F. & Jennings, G.T. Vaccine delivery: a matter of size, geometry, kinetics and molecular patterns. Nat. Rev. Immunol. 10, 787–796 (2010).

21. Irvine, D.J. & Read, B.J. Shaping humoral immunity to vaccines through antigen-displaying nanoparticles. Current Opinion in Immunology 65, 1–6 (2020).

22. Batista, F.D. & Neuberger, M.S. B cells extract and present immobilized antigen: implications for affinity discrimination. The EMBO Joural 19, 513–520 (2000).

23. Batista, F.D. & Harwood, N.E. The who, how and where of antigen presentation to B cells. Nature Reviews Immunology 9, 15–27 (2009).

24. Brooks, J.F. et al. Molecular basis for potent B cell responses to antigen displayed on particles of viral size. Nature Immunology 24, 1762–1777 (2023).

25. Kato, Y. et al. Multifaceted Effects of Antigen Valency on B Cell Response Composition and Differentiation In Vivo. Immunity 53, 548–563.e548 (2020).

26. Kraft, J.C. et al. Antigen- and scaffold-specific antibody responses to protein nanoparticle immunogens. Cell Reports Medicine 3, 100780 (2022).

27. Sliepen, K. et al. Interplay of diverse adjuvants and nanoparticle presentation of native-like HIV-1 envelope trimers. npj Vaccines 6, 103 (2021).

28. Wamhoff, E.-C. et al. Enhancing antibody responses by multivalent antigen display on thymus-independent DNA origami scaffolds. Nature Communications 15, 795 (2024).

29. Leggat, D.J. et al. Vaccination induces HIV broadly neutralizing antibody precursors in humans. Science 378, eadd6502 (2022).

30. Schiffner, T. et al. Diverse competitor B cell responses to a germline-targeting priming immunogen in human Ig loci transgenic mice. bioRxiv, 2024.2001.2022.575410 (2024).

31. Duan, H. et al. Glycan Masking Focuses Immune Responses to the HIV-1 CD4-Binding Site and Enhances Elicitation of VRC01-Class Precursor Antibodies. Immunity 49, 301–311 (2018).

32. Martina, C.E., Crowe, J.E., Jr. & Meiler, J. Glycan masking in vaccine design: Targets, immunogens and applications. Front. Immunol. 14 (2023).

33. Eto, Y. et al. Optimized PEGylated Adenovirus Vector Reduces the Anti-vector Humoral Immune Response against Adenovirus and Induces a Therapeutic Effect against Metastatic Lung Cancer. Biological and Pharmaceutical Bulletin 33, 1540–1544 (2010).

34. Abbott, R.A.-O. & Crotty, S.A.-O. Factors in B cell competition and immunodominance. Immunological Reviews 296, 120–131 (2020).

35. Wheatley, A.K. et al. Immune imprinting and SARS-CoV-2 vaccine design. Trends Immunol 42 (2021).

36. Schiepers, A. et al. Molecular fate-mapping of serum antibody responses to repeat immunization. Nature 615, 482–489 (2023).

37. Zarnitsyna, V.I., Lavine, J., Ellebedy, A., Ahmed, R. & Antia, R. Multi-epitope Models Explain How Pre-existing Antibodies Affect the Generation of Broadly Protective Responses to Influenza. PLOS Pathogens 12, e1005692 (2016).

38. Jardine, J.G. et al. HIV-1 broadly neutralizing antibody precursor B cells revealed by germline-targeting immunogen. Science 351, 1458–1463 (2016).

39. Jardine, J. et al. Rational HIV immunogen design to target specific germline B cell receptors. Science 340, 711–716 (2013).

40. Jardine, J.G. et al. Priming a broadly neutralizing antibody response to HIV-1 using a germline-targeting immunogen. Science 349, 156–161 (2015).

41. deCamp, A.C. et al. Human immunoglobulin gene allelic variation impacts germline-targeting vaccine priming. medRxiv, 2023.2003.2010.23287126 (2023).

42. Cohen, K.W. et al. A first-in-human germline-targeting HIV nanoparticle vaccine induced broad and publicly targeted helper T cell responses. Science Translational Medicine 15, eadf3309 (2023).

43. Andrabi, R., Bhiman, J.N. & Burton, D.R. Strategies for a multi-stage neutralizing antibody-based HIV vaccine. Curr. Opin. Immunol. 53 (2018).

44. Veneziano, R. et al. Role of nanoscale antigen organization on B-cell activation probed using DNA origami. Nat. Nanotechnol. 15, 716–723 (2020).

45. Wamhoff, E.-C. et al. Evaluation of Nonmodified Wireframe DNA Origami for Acute Toxicity and Biodistribution in Mice. ACS Applied Bio Materials 6, 1960–1969 (2023).

46. Lucas, C.R. et al. DNA Origami Nanostructures Elicit Dose-Dependent Immunogenicity and Are Nontoxic up to High Doses In Vivo. Small (2022).

47. Du, R.R. et al. Innate Immune Stimulation Using 3D Wireframe DNA Origami. ACS Nano (2022).

48. Comberlato, A., Koga, M.M., Nüssing, S., Parish, I.A. & Bastings, M.M.C. Spatially Controlled Activation of Toll-like Receptor 9 with DNA-Based Nanomaterials. Nano Letters 22, 2506–2513 (2022).

49. Jun, H. et al. Rapid prototyping of arbitrary 2D and 3D wireframe DNA origami. Nucleic Acids Res. 49, 10265–10274 (2021).

50. Rothemund, P.W.K. Folding DNA to create nanoscale shapes and patterns. Nature 440, 297–302 (2006).

51. Knappe, G.A., Wamhoff, E.-C. & Bathe, M. Functionalizing DNA origami to investigate and interact with biological systems. Nature Reviews Materials 8, 123–138 (2023).

52. Knappe, G.A., Wamhoff, E.-C., Read, B.J., Irvine, D.J. & Bathe, M. In Situ Covalent Functionalization of DNA Origami Virus-like Particles. ACS Nano 15, 14316–14322 (2021).

53. Veneziano, R. et al. Designer nanoscale DNA assemblies programmed from the top down. Science 352, 1534 (2016).

54. Silva, M. et al. A particulate saponin/TLR agonist vaccine adjuvant alters lymph flow and modulates adaptive immunity. Science Immunology 6, eabf1152 (2021).

55. Bradley, A. et al. The mammalian gene function resource: the International Knockout Mouse Consortium. Mamm Genome (2012).

56. Lacey, K.A. et al. Secreted mammalian DNases protect against systemic bacterial infection by digesting biofilms. Journal of Experimental Medicine 220, e20221086 (2023).

57. Cyster, J.G. B cell follicles and antigen encounters of the third kind. Nature Immunology 11, 989–996 (2010).

58. Swartz, M.A., Hubbell, J.A. & Reddy, S.T. Lymphatic drainage function and its immunological implications: From dendritic cell homing to vaccine design. Seminars in Immunology 20, 147–156 (2008).

59. Aung, A. et al. Low protease activity in B cell follicles promotes retention of intact antigens after immunization. Science 379, eabn8934 (2023).

60. Gonzalez, S.F. et al. Complement-Dependent Transport of Antigen into B Cell Follicles. The Journal of Immunology 185, 2659–2664 (2010).

61. Tokatlian, T. et al. Innate immune recognition of glycans targets HIV nanoparticle immunogens to germinal centers. Science 363, 649–654 (2019).

62. Phan, T.G., Grigorova, I., Okada, T. & Cyster, J.G. Subcapsular encounter and complement-dependent transport of immune complexes by lymph node B cells. Nature Immunology 8, 992–1000 (2007).

63. Reddy, S.T. et al. Exploiting lymphatic transport and complement activation in nanoparticle vaccines. Nature Biotechnology 25, 1159–1164 (2007).

64. Taban, Q., Mumtaz, P.T., Masoodi, K.Z., Haq, E. & Ahmad, S.M. Scavenger receptors in host defense: from functional aspects to mode of action. Cell Communication and Signaling 20, 2 (2022).

65. Ponnuswamy, N. et al. Oligolysine-based coating protects DNA nanostructures from low-salt denaturation and nuclease degradation. Nat. Commun. 8, 15654 (2017).

66. Zeng, Y.C. et al. Fine tuning of CpG spatial distribution with DNA origami for improved cancer vaccination. Nature Nanotechnology (2024).

67. Wang, W.X. et al. Universal, label-free, single-molecule visualization of DNA origami nanodevices across biological samples using origamiFISH. Nature Nanotechnology 19, 58–69 (2024).

68. Rodríguez-Franco, H.J., Weiden, J. & Bastings, M.M.C. Stabilizing Polymer Coatings Alter the Protein Corona of DNA Origami and Can Be Engineered to Bias the Cellular Uptake. ACS Polymers Au 3, 344–353 (2023).

69. Koga, M.M., Comberlato, A., Rodríguez-Franco, H.J. & Bastings, M.M.C. Strategic Insights into Engineering Parameters Affecting Cell Type-Specific Uptake of DNA-Based Nanomaterials. Biomacromolecules 23, 2586–2594 (2022).

70. Wamhoff, E.-C. et al. Controlling Nuclease Degradation of Wireframe DNA Origami with Minor Groove Binders. ACS Nano 16, 8954–8966 (2022).

71. Read, B.J. et al. Mannose-binding lectin and complement mediate follicular localization and enhanced immunogenicity of diverse protein nanoparticle immunogens. Cell Reports 38, 110217 (2022).

72. Sheriff, S., Chang Cy Fau - Ezekowitz, R.A. & Ezekowitz, R.A. Human mannose-binding protein carbohydrate recognition domain trimerizes through a triple alpha-helical coiled-coil. Nature Stuctural Biology (1994).

73. Ramiscal, R.R. & Vinuesa, C.G. T-cell subsets in the germinal center. Immunological Reviews 252 (2013).

74. Xu, Z. et al. Incorporation of a Novel CD4+ Helper Epitope Identified from Aquifex aeolicus Enhances Humoral Responses Induced by DNA and Protein Vaccinations. iScience 23, 101399 (2020).

75. La Rosa, C. et al. Clinical Evaluation of Safety and Immunogenicity of PADRE-Cytomegalovirus (CMV) and Tetanus-CMV Fusion Peptide Vaccines With or Without PF03512676 Adjuvant. The Journal of Infectious Diseases 205, 1294–1304 (2012).

76. Alexander, J. et al. Development of high potency universal DR-restricted helper epitopes by modification of high affinity DR-blocking peptides. Immunity 1, 751–761 (1994).

77. Alexander, J. et al. Linear PADRE T Helper Epitope and Carbohydrate B Cell Epitope Conjugates Induce Specific High Titer IgG Antibody Responses1. The Journal of Immunology 164, 1625–1633 (2000).

78. Tian, M. et al. Induction of HIV Neutralizing Antibody Lineages in Mice with Diverse Precursor Repertoires. Cell 166, 1471–1484.e1418 (2016).

79. Domange Jordö, E., Wermeling, F., Chen, Y. & Karlsson, M.C.I. Scavenger receptors as regulators of natural antibody responses and B cell activation in autoimmunity. Molecular Immunology 48, 1307–1318 (2011).

80. Caniels, T.G., et al. Germline-targeting HIV-1 Env vaccination induces VRC01-class antibodies with rare insertions. Cell Reports Medicine 4 (2023).

81. Heinimäki, S. et al. Antigenicity and immunogenicity of HA2 and M2e influenza virus antigens conjugated to norovirus-like, VP1 capsid-based particles by the SpyTag/SpyCatcher technology. Virology 566, 89-97 (2022).

82. Brinkkemper, M. et al. Mosaic and mixed HIV-1 glycoprotein nanoparticles elicit antibody responses to broadly neutralizing epitopes. PLOS Pathogens 20, e1012558 (2024).

83. Moghimi, S.M. et al. Material properties in complement activation. Advanced Drug Delivery Reviews 63, 1000–1007 (2011).

84. Trouw, L.A., Nilsson Sc Fau - Gonçalves, I., Gonçalves I Fau - Landberg, G., Landberg G Fau - Blom, A.M. & Blom, A.M. C4b-binding protein binds to necrotic cells and DNA, limiting DNA release and inhibiting complement activation. J Exp Med 12, 1937-1948 (2005).

85. Schiepers, A., Van’t Wout, M.F.L., Hobbs, A., Mesin, L. & Victora, G.D. Opposing effects of pre-existing antibody and memory T cell help on the dynamics of recall germinal centers. Immunity 57, 1618–1628 (2024).

## Methods References

86. Shepherd, T.R., Du, R.R., Huang, H., Wamhoff, E.-C. & Bathe, M. Bioproduction of pure, kilobase-scale single-stranded DNA. Scientific Reports 9, 6121 (2019).

87. Williams, S. et al. in DNA Computing. (eds. A. Goel, F.C. Simmel & P. Sosík) 90-101 (Springer Berlin Heidelberg, Berlin, Heidelberg; 2009).

88. Scheres, S.H.W. RELION: Implementation of a Bayesian approach to cryo-EM structure determination. Journal of Structural Biology 180, 519–530 (2012).

89. Sangesland, M. et al. Germline-Encoded Affinity for Cognate Antigen Enables Vaccine Amplification of a Human Broadly Neutralizing Response against Influenza Virus. Immunity 51, 735–749.e738 (2019).

90. Masella, A.P., Bartram, A.K., Truszkowski, J.M., Brown, D.G. & Neufeld, J.D. PANDAseq: paired-end assembler for illumina sequences. BMC bioinformatics 13, 31 (2012).

91. Shi, B. et al. Comparative analysis of human and mouse immunoglobulin variable heavy regions from IMGT/LIGM-DB with IMGT/HighV-QUEST. Theor Biol Med Model 11, 30 (2014).

92. Stecher, G., Tamura, K. & Kumar, S. Molecular Evolutionary Genetics Analysis (MEGA) for macOS. Mol Biol Evol 37 (2020).

93. Kumar, S., Stecher, G., Li, M., Knyaz, C. & Tamura, K. MEGA X: Molecular Evolutionary Genetics Analysis across Computing Platforms. Mol Biol Evol 6 (2018).

