## Supplementary Information for "DNA origami vaccines program antigen-focused germinal centers"

**This PDF file includes:**

Supplementary Figure 1. Overview of DNA-VLP synthesis

Supplementary Figure 2. Additional AGE verification of d40_30mer formulation.

Supplementary Figure 3. Flow cytometry gating for GC B and Tfh cell panel.

Supplementary Figure 4. Antigen probe staining in naïve mice.

Supplementary Figure 5. Gating for sorting CD4bs+ GC B cells for BCR sequencing.

Supplementary Table 1. Scaffold sequences

Supplementary Table 2. Staple oligonucleotide sequences

Supplementary Table 3. Primers for DNase I genotyping

Supplementary Table 4. Primers for BCR sequencing.

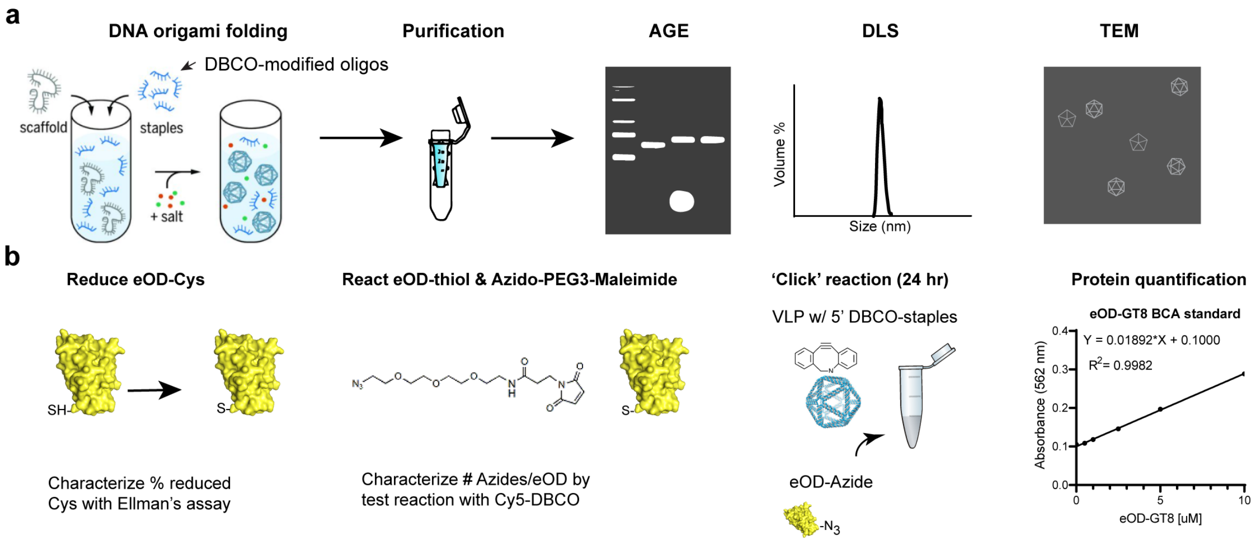


**Supplementary Fig. 1. Overview of DNA-VLP synthesis. a,** DNA origami is folded by mixing ssDNA circular DNA scaffold with pool of staple oligonucleotides containing a mixture of unmodified "vanilla" staples and 5' DBCO-modified staples. The particles are thermally annealed in an overnight folding ramp. Excess staples are purified away using Amicon centrifugal filters 100 kDa MWCO. The folding of the particles is verified using gel mobility assays, dynamic light scattering, and negative stain TEM. **b,** eOD-GT8 is expressed with a terminal cysteine. For antigen conjugation, eOD-GT8-Cys is treated with TCEP for thiol reduction then reacted with Azido-PEG3-Maleimide (reacted from SMCC and Azido-PEG3-Amine). Excess linker is removed by Zeba desalting columns or Amicon filters (10 kDa MWCO). The eOD-Azide is then reacted with DNA-VLPs folded with 5' DBCO-oligos and reacted for a minimum of 24 hours at 37°C. Excess protein is removed by drop dialysis into 1X PBS. The amount of reacted antigen is quantified using a micro-BCA assay, using known dilutions of eOD-GT8 as a calibration curve.

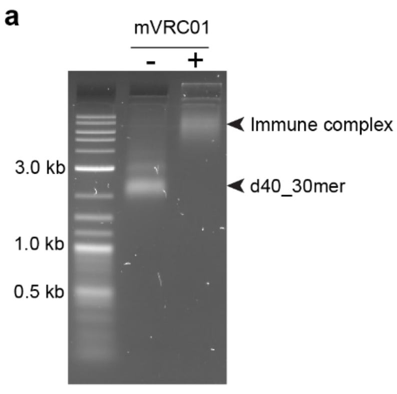


**Supplementary Fig. 2. Additional AGE verification of d40_30mer formulation. a**, 300 ng of d40_30mer was incubated with excess murine VRC01 antibody for 30 minutes in PBS, then analyzed on a 1.6% agarose gel. The reduced mobility through the gel indicates that the attached eOD-GT8 was bioavailable and properly oriented for antibody recognition. **b,** d40_30mer was mixed with SMNP adjuvant (at appropriate ratio to mimic injection doses) and incubated at room temperature in sterile PBS for 15 minutes prior to running on an agarose gel to mimic conditions prior to mouse injection. No aggregation was observed after mixing the VLPs with this adjuvant.

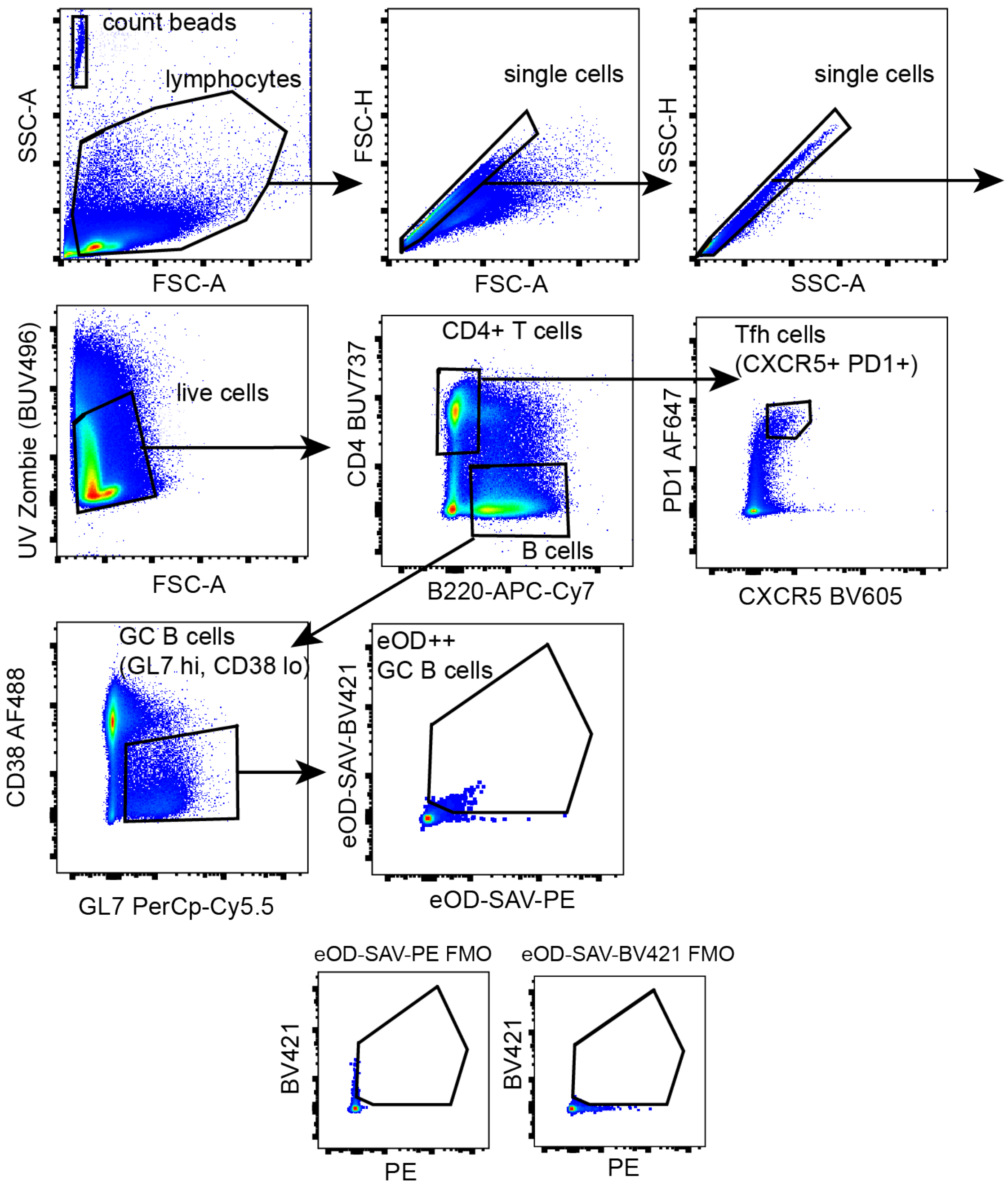


**Supplementary Fig. 3. Flow cytometry gating for GC B and Tfh cell panel.** Example flow gating used to identify GC B cells, follicular helper T cells, and antigen-specific GC B cells. Gates were drawn based on full-minus-one staining controls (FMOs). Example FMOs for antigen tetramer gating is shown at the bottom. Absolute cell counts were determined by calculating the ratio of count bead events to cell events.

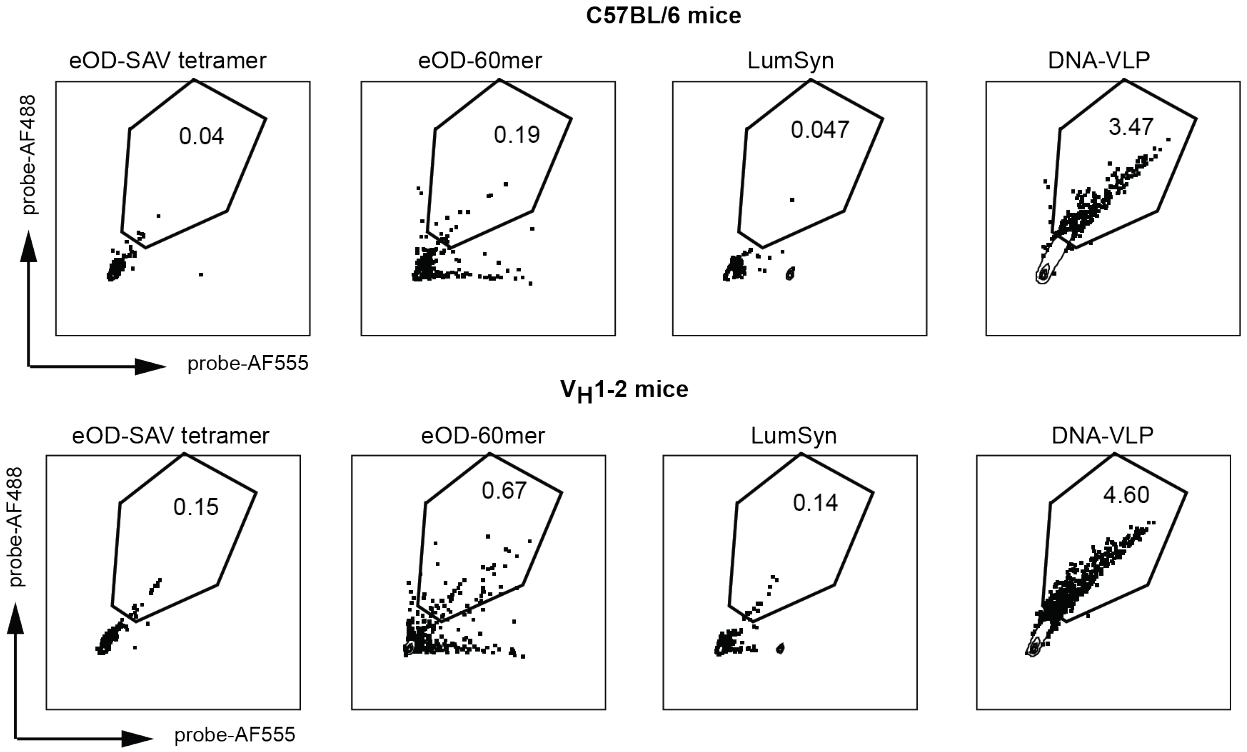


**Supplementary Fig. 4. Antigen probe staining in naïve mice.** Spleens from naïve C57BL/6, VH1-2, or VRC01$^{HHLL}$ mice were harvested, digested into single cell suspensions after red blood cell lysis, and stained with live/dead stain, B220 antibody, and indicated fluorescent flow probes (eOD-SAV tetramer, eOD-60mer, LumSyn, or bare DNA-VLPs). Shown is probe staining in B220$^{+}$ B cells. Protein antigen probes showed low background in C57BL/6 and VH1-2 cells. DNA-VLPs resulted in ~3-6% background binding in B cells from naïve mice of all 3 strains.

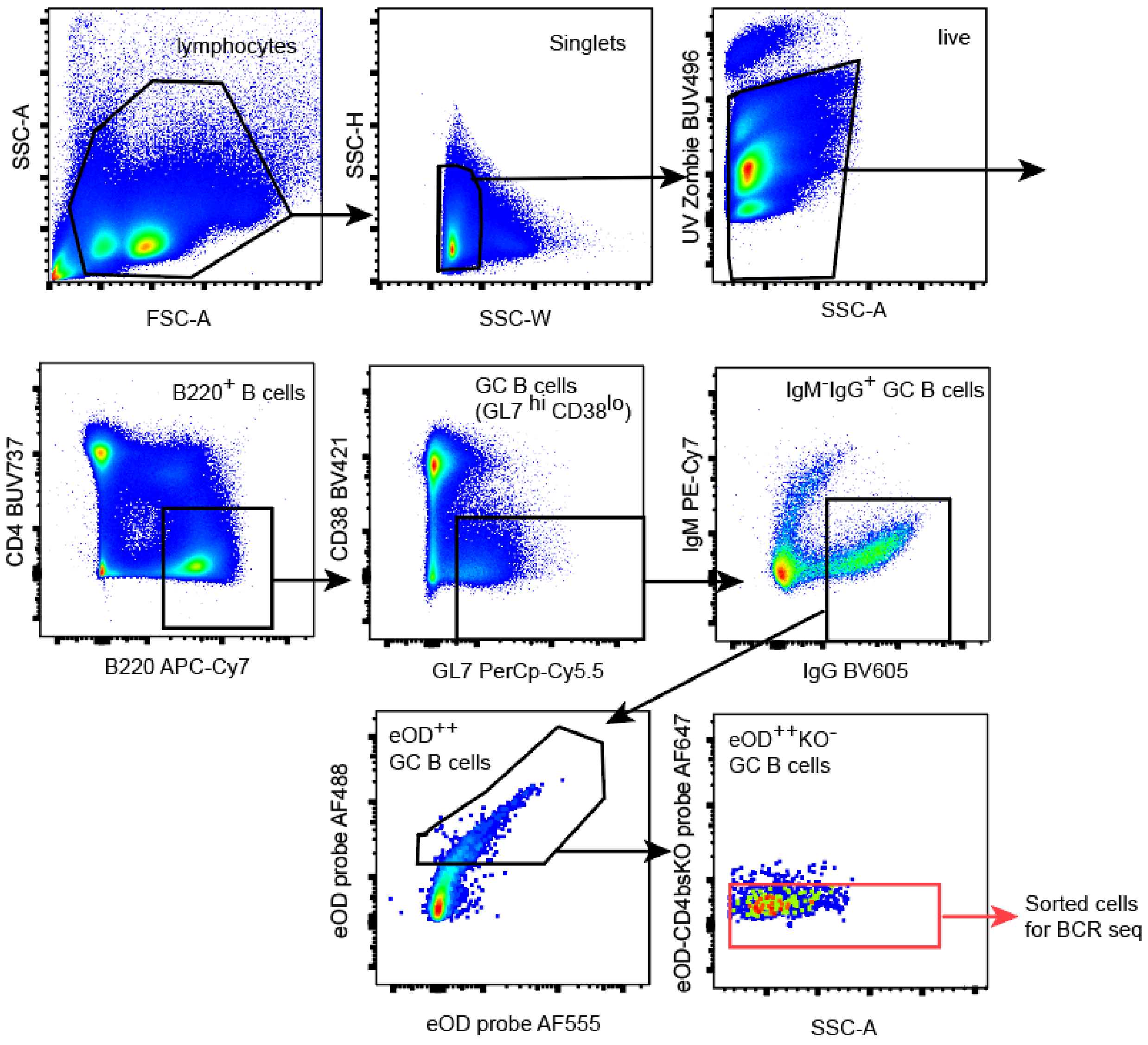


**Supplementary Fig. 5. Gating for sorting of CD4bs+ GC B cells for BCR sequencing.** Gating strategy used to identify live/B220$^{+}$/ GL7$^{hi}$ CD38$^{lo}$/IgM$^{-}$IgG$^{+}$/eOD$^{++}$ KO$^{-}$) CD4bs+ GC B cells. Gates were validated based on full-minus-one staining controls (FMOs).

**Supplementary Table 1. Scaffold sequences**

**3120 scaffold (for d40 VLPs)**

AGAATTTCGCCATAGGTATGCTTAAGGAAGTCGAGATTGCGAACCATTACCGAGACTATGGCTTCATGT
GGTGATTTCACCCGACCCACCCTTGGCGCCAGCTTTACGCAGCTTCCTGACGATACGTGGTGTAACG
TTGTGTTTGGCAATGGAAACCGAGATCAACTATTTCTAATGCTGATATAGCAGAGTCTCGCGTCTATCA
TACGCAAGTCGCACGTCATTTTCGAGAGCAGCGTAAGACTCTGAAGGTCATGAGCCCAGATGTTATTA
CCCTCTACCTATAAACATCAAAATTGTAGTCGTTTTACAGTCCATCGTCGCTCCAGAGCGAAGATTAAG
GTTAGATCTAGATTATCTTTGCACGTGTGGACCGACGCAGCTGGGGCTCTAGCTCCACTACGGTTACG
AAACTGCTGAACGATCTGGTCCACTTCAAGATTCACACATCGTTTCATTCTTTGGACAACCAACACTCT
CAGTCAGAGTTTCGAGTATAATAATTCTTCCGCGCTAGGGTAAAAAGCAGATATGGGGAGACATTCCG
GGCTTTTGAGCCGATACACTAAGCACTTGACATACTCACATCAGTAGAGGTTAACATTCATGACTATCA
CGCGCTGCAGGAGCGCAACGCAATTAATGTGCGCCCTGTAGCGGCGCATTAAGCGCGGCGGGTGTG
GTGGTTACGCGCAGCGTGACCGCTACACTTGCCAGCGCCCTAGCGCCCGCTCCTTTCGCTTTCTTCC
CTTCCTTTCTCGCCACGTTCGCCGGCTTTCCCCGTCAAGCTCTAAATCGGGGGCTCCCTTTAGGGTT
CCGATTTAGTGCTTTACGGCACCTCGACCCCAAAAAACTTGATTAGGGTGATGGTTCACGTAGTGGGC
CATCGCCCTGATAGACGGTTTTTCGCCCTTTGACGTTGGAGTCCACGTTCTTTAATAGTGGACTCTTG
TTCCAAACTGGAACAACACTCAACCCTATCTCGGTCTATTCTTTTGATTTATAAGGGATTTTGCCGATTT
CGGCCTATTGGTTAAAAAATGAGCTGATTTAACAAAAATTTAACGCGAATTACAACCGGGGTACATATGA
TTGGGGTCTGACGCTCAGTGGAACGAAAACTCACGTTAAGGGATTTTGGTCATGAGATTATCAAAAAG
GATCTTCACCTAGATCCTTTTAAATTAAAAATGAAGTTTTAAATCAATCTAAAGTATATATGAGTAAACTTG
GTCTGACAGTTACCAATGCTTAATCAGTGAGGCACCTATCTCAGCGATCTGTCTATTTCGTTCATCCAT
AGTTGCCTGACTCCCCGTCGTGTAGATAACTACGATACGGGAGGGCTTACCATCTGGCCCCAGTGCT
GCAATGATACCGCGAGACCCACGCTCACCGGCTCCAGATTTATCAGCAATAAACCAGCCAGCCGGAA
GGGCCGAGCGCATAAGTGGTCCTGCAACTTTATCCGCCTCCATCCAGTCTATTAATTGTTGCCGGGAA
GCTAGAGTAAGTAGTTCGCCAGTTAATAGTTTGCGCAACGTTGTTGCCATTGCTACAGGCATCGTGGT
GTCACGCTCGTCGTTTGGTATGGCTTCATTCAGCTCCGGTTCCCAACGATCAAGGCGAGTTACATGAT
CCCCCATGTTGTGCAAAAAAGCGGTTAGCTCCTTCGGTCCTCCGATCGTTGTCAGAAGTAAGTTGGC
CGCAGTGTTATCACTCATGGTTATGGCAGCACTGCATAATTCTCTTACTGTCATGCCATCCGTAAGATG
CTTTTCTGTGACTGGTGAGTACTCAACCAAGTCATTCTGAGAATAGTGTATGCGGCGACCGAGTTGCT
CTTGCCCGGCGTCAATACGGGATAATACCGCGCCACATAGCAGAACTTTAAAAGTGCTCATCATTGGA
AAACGTTCTTCGGGGCGAAAACTCTCAAGGATCTTACCGCTGTTGAGATCCAGTTCGATGTAACCCAC
TCGTGCACCCAACTGATCTTCAGCATCTTTTACTTTCACCAGCGTTTCTGGGTGAGCAAAAACAGGAA
GGCAAAATGCCGCAAAAAAGGGAATAAGGGCGACACGGAAATGTTGAATACTCATACTCTTCCTTTTT
CAATATTATTGAAGCATTTATCAGGGTTATTGTCTCATGAGCGGATACATATTTGAATGTATTTAGAAAAAT
AAACAAATAGGGGTTCCGCGCACATTTCCCCGAAAAGTGCCACCTGACGTCTAAGAAACCATTATTAT
CATGACATTAACCTATAAAAATAGGCGTATCACGAGGCCCTTTCGTCGAATTCGTCGTCGTCCCCTCAA
ACTCTTGGGTGGAGAGGCTATTCGTTTAAGGTCACATCGCATGTAATTTACTTATTCTCTGTTGTTGAG
CCACCCGGGCGCCAGATTTTGTTTAAAGCTTTGTCTCTTAGTTTGTATAGACAGATTCAGAGTGCAAG
GTTTCGTTCGCTCGTACCTGGTTTTCCCTGGTTCTTCACAGATAGGATTTGACTTTCTACAACACTTAT
GCGGCTTCCTACCCGTTTGAAGGCCGATACAGGTGCTGCGCAAAATGCGGGCGAACATAGAGTATCA
AAACAACGCCTTCTAATCTAGGAATATAGGGAAGATACGTATTTGCTACCATGCTTTCTTGGGTCATTAA
CGACCAACCTCTTTTCTTTTAAAGTAGGATTGCACAATGAATGAATACACGTGGTCCGATAACTGACCA
AGTAACATGGTTATCACTAGATGTCCGCCAGACGTGTGCAAACCAACCCGGGAGTTACGTCACTAATC
CTTCGCTACGTCGTGAAGATATTTACTTGTGAATATCGAGGGTAATAAGATAATAGACTGTGACTAGTAT
TGCCAGACTGTCGCTACCTGCAACACATAACTATCCTGAGGTTACTGCATAGTACTGATTACACCCGAG
TCAAAATTTCTAACTTCTAACATGTACCTAGTAACCAGCTCAATAATTATGTCAGAATATAGCTCTGGGAA
CCCTCGGACAATTATGATACACGGTATTAATATCTTGCTTGCGTTAGCCACTTCTCATCTTTGGATACCG
ATTCTATTTTGCATAGCAGTTCCTTTTACACATATA

**2520 scaffold (for d30 VLPs)**

GAGCGCAACGCAATTAATGTGCGCCCTGTAGCGGCGCATTAAGCGCGGCGGGTGTGGTGGTTACG
CGCAGCGTGACCGCTACACTTGCCAGCGCCCTAGCGCCCGCTCCTTTCGCTTTCTTCCCTTCCTTTC

TCGCCACGTTCGCCGGCTTTCCCCGTCAAGCTCTAAATCGGGGGCTCCCTTTAGGGTTCCGATTTAG
TGCTTTACGGCACCTCGACCCCAAAAAACTTGATTAGGGTGATGGTTCACGTAGTGGGCCATCGCCC
TGATAGACGGTTTTTCGCCCTTTGACGTTGGAGTCCACGTTCTTTAATAGTGGACTCTTGTTCCAAAC
TGGAACAACACTCAACCCTATCTCGGTCTATTCTTTTGATTTATAAGGGATTTTGCCGATTTCGGCCT
ATTGGTTAAAAAATGAGCTGATTTAACAAAAATTTAACGCGAATTACAACCGGGGTACATATGATTGG
GGTCTGACGCTCAGTGGAACGAAAACTCACGTTAAGGGATTTTGGTCATGAGATTATCAAAAAGGAT
CTTCACCTAGATCCTTTTAAATTAAAAATGAAGTTTTAAATCAATCTAAAGTATATATGAGTAAACTTGG
TCTGACAGTTACCAATGCTTAATCAGTGAGGCACCTATCTCAGCGATCTGTCTATTTCGTTCATCCAT
AGTTGCCTGACTCCCCGTCGTGTAGATAACTACGATACGGGAGGGCTTACCATCTGGCCCCAGTGC
TGCAATGATACCGCGAGACCCACGCTCACCGGCTCCAGATTTATCAGCAATAAACCAGCCAGCCGG
AAGGGCCGAGCGCAtAAGTGGTCCTGCAACTTTATCCGCCTCCATCCAGTCTATTAATTGTTGCCGG
GAAGCTAGAGTAAGTAGTTCGCCAGTTAATAGTTTGCGCAACGTTGTTGCCATTGCTACAGGCATCG
TGGTGTCACGCTCGTCGTTTGGTATGGCTTCATTCAGCTCCGGTTCCCAACGATCAAGGCGAGTTAC
ATGATCCCCCATGTTGTGCAAAAAAGCGGTTAGCTCCTTCGGTCCTCCGATCGTTGTCAGAAGTAAG
TTGGCCGCAGTGTTATCACTCATGGTTATGGCAGCACTGCATAATTCTCTTACTGTCATGCCATCCGT
AAGATGCTTTTCTGTGACTGGTGAGTACTCAACCAAGTCATTCTGAGAATAGTGTATGCGGCGACCG
AGTTGCTCTTGCCCGGCGTCAATACGGGATAATACCGCGCCACATAGCAGAACTTTAAAAGTGCTCA
TCATTGGAAAACGTTCTTCGGGGCGAAAACTCTCAAGGATCTTACCGCTGTTGAGATCCAGTTCGAT
GTAACCCACTCGTGCACCCAACTGATCTTCAGCATCTTTTACTTTCACCAGCGTTTCTGGGTGAGCAA
AAACAGGAAGGCAAAATGCCGCAAAAAAGGGAATAAGGGCGACACGGAAATGTTGAATACTCATACT
CTTCCTTTTTCAATATTATTGAAGCATTTATCAGGGTTATTGTCTCATGAGCGGATACATATTTGAATG
TATTTAGAAAAATAAACAAATAGGGGTTCCGCGCACATTTCCCCGAAAAGTGCCACCTGACGTCTAA
GAAACCATTATTATCATGACATTAACCTATAAAAATAGGCGTATCACGAGGCCCTTTCGTCGAATTCG
TCGTCGTCCCCTCAAACTCTTGGGTGGAGAGGCTATTCGTTTAAGGTCACATCGCATGTAATTTACTT
ATTCTCTGTTGTTGAGCCACCCGGGCGCCAGATTTTGTTTAAAGCTTTGTCTCTTAGTTTGTATAGAC
AGATTCAGAGTGCAAGGTTTCGTTCGCTCGTACCTGGTTTTCCCTGGTTCTTCACAGATAGGATTTGA
CTTTCTACAACACTTATGCGGCTTCCTACCCGTTTGAAGGCCGATACAGGTGCTGCGCAAAATGCGG
GCGAACATAGAGTATCAAAACAACGCCTTCTAATCTAGGAATATAGGGAAGATACGTATTTGCTACCA
TGCTTTCTTGGGTCATTAACGACCAACCTCTTTTCTTTTAAAGTAGGATTGCACAATGAATGAATACAC
GTGGTCCGATAACTGACCAAGTAACATGGTTATCACTaGATGTCCGCCAGACGTGTGCAAACCAACC
CGGGAGTTACGTCACTAATCCTTCGCTACGTCGTGAAGATATTTACTTGTGAATATCGAGGGTAATAA
GATAATAGACTGTGACTAGTATTGCCAGACTGTCGCTACCTGCAACACATAACTATCCTGAGGTTACT
GCATAGTACTGATTACACCCGAGTCAAAATTTCTAACTTCTAACATGTACCTAGTAACCAGCTCAATAA
TTATGTCAGAATATAGCTCTGGGAACCCTCGGACAATTATGATACACGGTATTAATATCTTGCTTGCG
TTAGCCACTTCTCATCTTTGGATACCGATTCTATTTTGCATAGCAGTTCCTTTTACACATATAAGAATTT
CGCCATAGGTATGCTGCAG

**Supplementary Table 2. Staple oligonucleotide sequences**

**d40 staple sequences**

| Number | Sequence |
|---|---|
| 2 | ACATCTGGATGGCGAAATTCTTATATGTGGTAGAGGGTAATA |
| 3 | GCTCATGACCAAGCATACCT |
| 4 | GTTCGCAATCTTTTTTCGACTTCCTTTTCAGAGTCTTTTTTTACGCTGCTCT |
| 5 | CGGTAATGTCACCACATGAAGCCGTCCGAGG |
| 6 | GTTCCCAGACCGTGTATCATAATTATAGTCT |
| 7 | GCGAGACTCTGTTTTTCTATATCAGCCCAAACACAACTTTTTGTTACACCACCCAAGGGTGGGTTTTTTCGGGTGAAA |
| 8 | AGCTGGCGGTATCGTCAGGAAGCTAACCTCA |
| 9 | GGATAGTTAATCAGTACTATGCAGTGCGTAA |
| 10 | TTCCATTGATTAGAAATAGTTGAGTAGCGAA |
| 11 | GGATTAGTTAAATATCTTCACGACTCTCGGT |
| 12 | TGATAGACCGAAAATGACGTGCGACACGTGC |
| 13 | AAAGATAAAGCTGCGTCGGTCCACTTGCGTA |
| 14 | CTACAATTTTGTTTTTATGTTTATAGTAAAAGGAACTTTTTTGCTATGCAAATTTTTATAGGTTTTTTTAATGTCATG |
| 15 | GTCAAGTGCGACGATGGACTGTAAAACGACTGATGTGAGTAT |
| 16 | CTTAGTGTATCGCTCTGGAG |
| 17 | TCTAGATCTAATTTTTCCTTAATCTTCGGCTCAAAAGTTTTTCCCGGAATGT |
| 18 | AGAATTATTATTTTTTACTCGAAACTGAAACGATGTGTTTTTTGAATCTTGACCGTAGTGGAGTTTTTCTAGAGCCCC |
| 19 | TTTCGTAAAGTGGACCAGATCGTACATCTAG |
| 20 | TGATAACCTGCACACGTCTGGCGGTCAGCAG |
| 21 | CAAAGAATCTGACTGAGAGTGTTTGATAACA |
| 22 | CTGCGGCCCTGCCATAACCATGAGGGTTGTC |
| 23 | AGCGCGGACTCCCCATATCTGCTTACCCGCC |
| 24 | GCGCTTAAGCGCGTAACCACCACTTTACCCT |
| 25 | CAACGTCAAAGTTTTTGGCGAAAAACAGTCATGAATGTTTTTTTAACCTCTA |
| 26 | GATGGCCCTGCGCTCCTGCAGCGCGTGATCGTCTATCAGGGC |
| 27 | ACTACGTGAATTAATTGCGT |
| 28 | TGCGCCGCTACTTTTTAGGGCGCACACCATCACCCTATTTTTATCAAGTTTT |
| 29 | GAACCCTAAAGTTTTTGGAGCCCCCGCGTGGCGAGAATTTTTAGGAAGGGAAGGCAAGTGTAGTTTTTCGGTCACGCT |
| 30 | AGGGCGCTGAAAGCGAAAGGAGCTAACCGCT |
| 31 | TTTTTGCAGGAGGACCGAAGGAGCGGGCGCT |
| 32 | CCGGCGAAATTTAGAGCTTGACGATCATATG |
| 33 | TACCCCGGTGAGCGTCAGACCCCAGGGAAAG |
| 34 | CTAAATCGTTGGGGTCGAGGTGCCAGACCGA |
| 35 | GATAGGGTATAAATCAAAAGAATGTAAAGCA |
| 36 | CTTAGACGACTATTAAAGAACGTGGACTCATAATAATGGTTT |
| 37 | TCAGGTGGCAACAAGAGTCC |
| 38 | TGAGTGTTGTTTTTTTCCAGTTTGGACTTTTCGGGGATTTTTAATGTGCGCG |
| 39 | ATTCAAATATGTTTTTTATCCGCTCACGTAGTTATCTTTTTTACACGACGGGCCGAAATCGGCTTTTTAAAATCCCTT |

40 ATGGATGAGCTCATTTTTTAACCAATAGGGAGTCAGGCAACT
41 ACGAAATAGATGTTAAATCA
42 TAACGTGAGTTTTTTTTTCGTTCCACTTGTAATTCGCTTTTTGTTAAATTTTCAGATCGCTGATTTTTGATAGGTGCC
43 AAATCCCTCTTTTTGATAATCTCGAACCGGA
44 GCTGAATGTCGCCTTGATCGTTGGATGACCA
45 CAAGTTTACTCTTTTTATATATACTTAAAAGGATCTATTTTTGGTGAAGATC
46 TTTAATTTTAGATTGATTTAAAACTAGCTTC
47 CCGGCAACTGGCGAACTACTTACTCTTCATT
48 TGTCAGACTCACTGATTAAGCATGGAGCCGG
49 TGAGCGTGTATTGCTGATAAATCTTGGTAAC
50 CTGATAAAAGATGGTAAGCCCTCCCGTATTGAGACAATAACC
51 TGCTTCAATACACTGGGGCC
52 GGTCTCGCGGTTTTTTATCATTGCAGATATTGAAAAATTTTTGGAAGAGTATCCTTTTTTGCGTTTTTGCATTTTGCC
53 TGAAAGTAAAATTTTTGATGCTGAAGGGCCCTTCCGGTTTTTCTGGCTGGTT
54 ACGAGTGGTGCAGGACCACTTATGCGCTCATCAGTTGGGTGC
55 GTTACATCGACGGATAAAGT
56 AATTAATAGACTTTTTTGGATGGAGGACTGGATCTCATTTTTACAGCGGTAA
57 TTCCAATGATGTTTTTAGCACTTTTAATTGACGCCGGTTTTTGCAAGAGCAAAACGTTGCGCATTTTTAACTATTAAC
58 CAATGGCAACCTCGGTCGCC
59 GCCTGTAGGCATACACTATTCTCAGAATGGTGACACCACGAT
60 CAACATGGGGGTTTTTATCATGTAACAAGCCATACCATTTTTAACGACGAGCACTTGGTTGAGTTTTTTACTCACCAG
61 CAGTAAGAGAATTTTTTTATGCAGTGAACTTACTTCTTTTTTGACAACGATC
62 TGGCATGATCACAGAAAAGCATCTCATTGTG
63 CAATCCTAGACCACGTGTATTCATTTACGGA
64 TATCCCGTAAGTTCTGCTATGTGGGAAAGCA
65 TGGTAGCATCGTTAATGACCCAACGCGGTAT
66 AGAACGTTGATCCTTGAGAGTTTTCTTCAAA
67 CGGGTAGGAGCACCTGTATCGGCCGCCCCGA
68 AACGCTGGTTCCTGTTTTTGCTCGTGAAGAA
69 CCAGGGAAAAGTCAAATCCTATCTACCCAGA
70 CCTTATTCGAGTATTCAACATTTCTGGCTCA
71 ACAACAGAAATCTGGCGCCCGGGCGTGTCGC
72 CTAAATACGAACCCCTATTTGTTTAGCCTCT
73 CCACCCAATGTGACCTTAAACGAATATTTTT
74 CCAAAGATAAGGGCCTCGTGATACGCCTAATAGAATCGGTAT
75 GAGAAGTGGCAATTCGACGA
76 GAATAAGTAAATTTTTTTACATGCGAGAGTTTGAGGGTTTTTGACGACGACG
77 TAACGCAAGCATTTTTAGATATTAATAGCTATATTCTTTTTTGACATAATTAAAGAGACAAAGTTTTTCTTTAAACAA
78 TAGGTACACTGAATCTGTCTATACAAACTTTGAGCTGGTTAC
79 TGTTAGAAGTACCTTGCACT
80 AAGCCGCATAATTTTTGTGTTGTAGAAACCAGGTACGTTTTTAGCGAACGAA
81 TAGAAATTTTGTTTTTACTCGGGTGTATGTGTTGCAGTTTTTGTAGCGACAGATGTTCGCCCGTTTTTCATTTTGCGC
82 TTGATACTCTTCTGGCAATA

83 GCGTTGTTCTAGTCACAGTCTATTATCTTCCTAGATTAGAAG

84 ATGTTACTTGGTTTTTTCAGTTATCGCTTTAAAAGAATTTTTAAGAGGTTGGAATACGTATCTTTTTTTCCCTATATT

85 ATTACCCTCGATTTTTTATTCACAAGGACGTAACTCCTTTTTCGGGTTGGTT

**Modified oligos (d40_30mer)**

2 |DBCO|TTACATCTGGATGGCGAAATTCTTATATGTGGTAGAGGGTAATA

5 |DBCO|TTCGGTAATGTCACCACATGAAGCCGTCCGAGG

9 |DBCO|TTGGATAGTTAATCAGTACTATGCAGTGCGTAA

11 |DBCO|TTGGATTAGTTAAATATCTTCACGACTCTCGGT

13 |DBCO|TTAAAGATAAAGCTGCGTCGGTCCACTTGCGTA

15 |DBCO|TTGTCAAGTGCGACGATGGACTGTAAAACGACTGATGTGAGTAT

20 |DBCO|TTTGATAACCTGCACACGTCTGGCGGTCAGCAG

22 |DBCO|TTCTGCGGCCCTGCCATAACCATGAGGGTTGTC

24 |DBCO|TTGCGCTTAAGCGCGTAACCACCACTTTACCCT

26 |DBCO|TTGATGGCCCTGCGCTCCTGCAGCGCGTGATCGTCTATCAGGGC

31 |DBCO|TTTTTTTGCAGGAGGACCGAAGGAGCGGGCGCT

32 |DBCO|TTCCGGCGAAATTTAGAGCTTGACGATCATATG

35 |DBCO|TTGATAGGGTATAAATCAAAAGAATGTAAAGCA

36 |DBCO|TTCTTAGACGACTATTAAAGAACGTGGACTCATAATAATGGTTT

40 |DBCO|TTATGGATGAGCTCATTTTTTAACCAATAGGGAGTCAGGCAACT

44 |DBCO|TTGCTGAATGTCGCCTTGATCGTTGGATGACCA

47 |DBCO|TTCCGGCAACTGGCGAACTACTTACTCTTCATT

48 |DBCO|TTTGTCAGACTCACTGATTAAGCATGGAGCCGG

50 |DBCO|TTCTGATAAAAGATGGTAAGCCCTCCCGTATTGAGACAATAACC

54 |DBCO|TTACGAGTGGTGCAGGACCACTTATGCGCTCATCAGTTGGGTGC

59 |DBCO|TTGCCTGTAGGCATACACTATTCTCAGAATGGTGACACCACGAT

62 |DBCO|TTTGGCATGATCACAGAAAAGCATCTCATTGTG

64 |DBCO|TTTATCCCGTAAGTTCTGCTATGTGGGAAAGCA

67 |DBCO|TTCGGGTAGGAGCACCTGTATCGGCCGCCCCGA

68 |DBCO|TTAACGCTGGTTCCTGTTTTTGCTCGTGAAGAA

71 |DBCO|TTACAACAGAAATCTGGCGCCCGGGCGTGTCGC

73 |DBCO|TTCCACCCAATGTGACCTTAAACGAATATTTTT

74 |DBCO|TTCCAAAGATAAGGGCCTCGTGATACGCCTAATAGAATCGGTAT

78 |DBCO|TTTAGGTACACTGAATCTGTCTATACAAACTTTGAGCTGGTTAC

83 |DBCO|TTGCGTTGTTCTAGTCACAGTCTATTATCTTCCTAGATTAGAAG

**Modified oligos (d40_60mer)**

2 |DBCO|TTACATCTGGATGGCGAAATTCTTATATGTGGTAGAGGGTAATA

5 |DBCO|TTCGGTAATGTCACCACATGAAGCCGTCCGAGG

9 |DBCO|TTGGATAGTTAATCAGTACTATGCAGTGCGTAA

11 |DBCO|TTGGATTAGTTAAATATCTTCACGACTCTCGGT

13 |DBCO|TTAAAGATAAAGCTGCGTCGGTCCACTTGCGTA

15 |DBCO|TTGTCAAGTGCGACGATGGACTGTAAAACGACTGATGTGAGTAT

20 |DBCO|TTTGATAACCTGCACACGTCTGGCGGTCAGCAG

22 |DBCO|TTCTGCGGCCCTGCCATAACCATGAGGGTTGTC

24 |DBCO|TTGCGCTTAAGCGCGTAACCACCACTTTACCCT

26 |DBCO|TTGATGGCCCTGCGCTCCTGCAGCGCGTGATCGTCTATCAGGGC

31 |DBCO|TTTTTTTGCAGGAGGACCGAAGGAGCGGGCGCT

32 |DBCO|TTCCGGCGAAATTTAGAGCTTGACGATCATATG

35 |DBCO|TTGATAGGGTATAAATCAAAAGAATGTAAAGCA

36 |DBCO|TTCTTAGACGACTATTAAAGAACGTGGACTCATAATAATGGTTT

40 |DBCO|TTATGGATGAGCTCATTTTTTAACCAATAGGGAGTCAGGCAACT

44 |DBCO|TTGCTGAATGTCGCCTTGATCGTTGGATGACCA

47 |DBCO|TTCCGGCAACTGGCGAACTACTTACTCTTCATT

48 |DBCO|TTTGTCAGACTCACTGATTAAGCATGGAGCCGG

50 |DBCO|TTCTGATAAAAGATGGTAAGCCCTCCCGTATTGAGACAATAACC

54 |DBCO|TTACGAGTGGTGCAGGACCACTTATGCGCTCATCAGTTGGGTGC

59 |DBCO|TTGCCTGTAGGCATACACTATTCTCAGAATGGTGACACCACGAT

62 |DBCO|TTTGGCATGATCACAGAAAAGCATCTCATTGTG

64 |DBCO|TTTATCCCGTAAGTTCTGCTATGTGGGAAAGCA

67 |DBCO|TTCGGGTAGGAGCACCTGTATCGGCCGCCCCGA

68 |DBCO|TTAACGCTGGTTCCTGTTTTTGCTCGTGAAGAA

71 |DBCO|TTACAACAGAAATCTGGCGCCCGGGCGTGTCGC

73 |DBCO|TTCCACCCAATGTGACCTTAAACGAATATTTTT

74 |DBCO|TTCCAAAGATAAGGGCCTCGTGATACGCCTAATAGAATCGGTAT

78 |DBCO|TTTAGGTACACTGAATCTGTCTATACAAACTTTGAGCTGGTTAC

83 |DBCO|TTGCGTTGTTCTAGTCACAGTCTATTATCTTCCTAGATTAGAAG

6 |DBCO|TTGTTCCCAGACCGTGTATCATAATTATAGTCT

8 |DBCO|TTAGCTGGCGGTATCGTCAGGAAGCTAACCTCA

10 |DBCO|TTTTCCATTGATTAGAAATAGTTGAGTAGCGAA

12 |DBCO|TTTGATAGACCGAAAATGACGTGCGACACGTGC

19 |DBCO|TTTCGTAAAGTGGACCAGATCGTACATCTAG

21 |DBCO|TTCAAAGAATCTGACTGAGAGTGTTTGATAACA

23 |DBCO|TTAGCGCGGACTCCCCATATCTGCTTACCCGCC

30 |DBCO|TTAGGGCGCTGAAAGCGAAAGGAGCTAACCGCT

33 |DBCO|TTTACCCCGGTGAGCGTCAGACCCCAGGGAAAG

34 |DBCO|TTCTAAATCGTTGGGGTCGAGGTGCCAGACCGA

43 |DBCO|TTAAATCCCTCTTTTTGATAATCTCGAACCGGA

46 |DBCO|TTTTTAATTTTAGATTGATTTAAAACTAGCTTC

49 |DBCO|TTTGAGCGTGTATTGCTGATAAATCTTGGTAAC

63 |DBCO|TTCAATCCTAGACCACGTGTATTCATTTACGGA
65 |DBCO|TTTGGTAGCATCGTTAATGACCCAACGCGGTAT
66 |DBCO|TTAGAACGTTGATCCTTGAGAGTTTTCTTCAAA
69 |DBCO|TTCCAGGGAAAAGTCAAATCCTATCTACCCAGA
70 |DBCO|TTCCTTATTCGAGTATTCAACATTTCTGGCTCA
72 |DBCO|TTCTAAATACGAACCCCTATTTGTTTAGCCTCT
3 |DBCO|TTGCTCATGACCAAGCATACCT
16 |DBCO|TTCTTAGTGTATCGCTCTGGAG
27 |DBCO|TTACTACGTGAATTAATTGCGT
37 |DBCO|TTTCAGGTGGCAACAAGAGTCC
41 |DBCO|TTACGAAATAGATGTTAAATCA
51 |DBCO|TTTGCTTCAATACACTGGGGCC
55 |DBCO|TTGTTACATCGACGGATAAAGT
58 |DBCO|TTCAATGGCAACCTCGGTCGCC
75 |DBCO|TTGAGAAGTGGCAATTCGACGA
79 |DBCO|TTTGTTAGAAGTACCTTGCACT
82 |DBCO|TTTTGATACTCTTCTGGCAATA

**d30 sequences**

2 CTATGGCTTTTTGAAATTCTTAATCGGTATCCATTTTTAAGATGAGAAGTGGCGTCAAAGG
3 AAAATAGATATGTGTAAAAGGAACGTAGCGGT
4 CGGGGAATTTTTAGCCGGCGAAGAGCGGGCGCTTTTTTAGGGCGCTGGCAAGTTGCTATGC
5 AGCGAAAGCGTGGCGAGAAAGGAACGTGAACC
6 TATCAGTTTTTGGCGATGGCCCACTAGGGAAGAA
7 TATAAATCTTTTTAAAAGAATAGGAACAAGAGTCTTTTTCACTATTAAACCGTC
8 GCGAAAAAGAACGTGGACTCCAACTAACGCAA
9 CCAGTTTGACCGAGATAGGGTTGAGGTCAGTT
10 CGAAGGTTTTTATTAGTGACGGACATCTAGTGTTTTTATAACCATGTTACTTGTGTTGTT
11 GTCTGGCGTAACTCCCGGGTTGGTATAATTGT
12 GCAAGATATTTTTTTAATACCGTGTATCTTGCACAC
13 GGAAGCCGCCTGTATCGGCCTTCATGGTCGTTAATGACCCTTTTTAAGAAAGCATCGTAG
14 GTATCTTCCCACAAGTAAATATCTTCACGAGGTAGCAAATAC
15 TTACTTTTTCCTCGATATTCTATATTCCTATTTTTGATTAGAAGGTAAA
16 TTATGTTTTTTGTTGCAGGTATATCTTA
17 CAGTCTATGCGACAGTCTGGCAATCTGACAT
18 CCGAGGGTTTTTTTCCCAGAGCTATATTACTAGTCA
19 TTACATGTTTTTCGATGTGACCGAGTATTCAACTTTTTATTTCCGTGTGATAG
20 TTTTTTGCAGTACTATGCAGTAACCTCAGCGCCCTTATTCCC
21 TACAGGTTTTTGCGCACATTAAATTTTGACTCTTTTTGGGTGTAATCGGCA
22 AAGTTAGAATTGCGTTGCGCTCCTGCAGCAGGTACATGTTAG
23 AATTATTGATTTTTGCTGGTTACTATAC

24 ACGCTGGTTTTTTGAAAGTAAAATTACATCGAACTTTTTTGGATCTCAAGCCGC
25 CTTGAGAGCACACCCGCCGCGCTTAATGCCAGCGGTAAGATC
26 CACGCTGCTTTTTGCGTAACCACTTTTCGCCCCGTTTTTAAGAACGTTTTCCAAAAAAGCAT
27 CGAGTGGGGATGCTGAAGATCAGTGTGCTGCC
28 GGTTGAGTGGCATGACAGTTTTTTAAGAGAATTATGCATGGGTGCA
29 CTTACGGATACTCACCAGTCACAGTGATGAGCACTTTTAATTTTTAGTTCTGCTATTGA
30 GGGGGATTTTTTCATGTAACTTCTCAGAATGATTTTTCTT
31 TACACTATCGCCTTGATCGTTGGGAACCGCTCGGTCGCCGCA
32 TAAATCTTTTTGGAACCCTAAATTGACGCCGGTTTTTGCAAGAGCAAGAGC
33 TATCCCGTAGGGAGCCCCCGATTTAGAGCTGTGGCGCGGTAT
34 TAAGCCTTTTTCTCCCGTATCCCGGTTGTAATTTTTTTCGCGTTAAAAGCAC
35 AGCTCATTTTTGGGGTCGAGGTGCCGTAATTTTTGTTAAATC
36 ATCACCCTTTTTTAATCAAGTTTTTTTAACCAATTTTTTAGGCCGAAATCGGCAAAAATCCC
37 CGACGGGGCAGACCCCAATCATATGTACCGTAGTTATCTACA
38 AGCGTAGTCAGGCAACTTTTTTATGGATGAACGAAATGATAAAT
39 TAGGTGATTTTTAGATCCTTTTAGTTTTCGTTCTTTTTCACTG
40 TTAACGTGTGATAATCTCATGACCAAATCCCT
41 TCGCTGAGTTTTTATAGGTGCCTGTTTACTCATATTTTTTATACTTTAGGATC
42 ATCGGACCTTTTTACGTGTATTCATTCAATTTTTAATTTAAAAGATTGATTTAAAACTTCTTGTGCAA
43 CAGACCAACACTGATTAAGCATTGTATCTGTG
44 CGCAGCACATAAGTGTTGTTTTTTAGAAAGTCAAATCCGTAACTGT
45 TCCTACTTTTTTTTAAAAGAAAAGAGGTAACGGGTA
46 GAAACCTTTTTTTGCACTCTGAAAACAAAATCTGTTTTTGCGCCCGGGTTCGCCCGCATTTTTTTTTG
47 CTCTATGTGGCTCAACAACAGAGAATAAGCGTTGTTTTGATA
48 AAGCTTTATCTGTCTATACAAACTATTCGACGA
49 AGTTTGTTTTTAGGGGACGACGACGAAAGAGACA
50 TTTATTTTTTTTTTCTAAATAACCCTGATAAATTTTTTGCTTCAATACCAAG
51 AAGAGTATTTAAACGAATAGCCTCTCCACATATTGAAAAAGG
52 AGACAATACATTCAAATATGTATCACCCAGAA
53 TTTTGCCTTTTTTTCCTGTTTTTGCTCCGCTCATG
54 AAGGGCCTTTTTTCGTGATACGTGGTTTCTTAGTTTTTACGTCAGGTGATTTG
55 GAACCCCTGCACTTTTCGGGGAAAGGCCAACT
56 ATAACCATTTTTTGAGTGATAACACTGCTGTGCGCG
57 TACTTCTGTTTTTACAACGATCGAACAT
58 TTTTGCACGAGGACCGAAGGAGCTGCGCAAAC
59 TGTAGCTTTTTAATGGCAACAACGTTAACCGCTT
60 CGCTCGTTTTTGCCCTTCCGGGGTGAGCGTGGTTTTTGTCTCGCGGTATGCC
61 TGGGGCCAAACGACGAGCGTGACACCACGATCATTGCAGCAC
62 TGAATGATTTTTAGCCATACCAGATGG
63 CTGGAGCCCTGGCTGGTTTATTGCTAGACAGA
64 GATAATAACCTATTTTTATAGGTTCTAGCTTC
65 TATTAACTTTTTTGGCGAACTACTTACTAATGTCAT
66 CCGGCAACTTTTTAATTAATAGATTATG

67 AAGAACCATTTTTGGGAAAACCAGGTACAAAGTTGCAGGACCACCTGGATGGAGGCGGATGAGCGAAC

**Modified oligos (d30_30mer)**

3 |DBCO|TTAAAATAGATATGTGTAAAAGGAACGTAGCGGT
5 |DBCO|TTAGCGAAAGCGTGGCGAGAAAGGAACGTGAACC
8 |DBCO|TTGCGAAAAAGAACGTGGACTCCAACTAACGCAA
9 |DBCO|TTCCAGTTTGACCGAGATAGGGTTGAGGTCAGTT
11 |DBCO|TTGTCTGGCGTAACTCCCGGGTTGGTATAATTGT
13 |DBCO|TTGGAAGCCGCCTGTATCGGCCTTCATGGTCGTTAATGACCCTTTTTAAGAAAGCATCGTAG
14 |DBCO|TTGTATCTTCCCACAAGTAAATATCTTCACGAGGTAGCAAATAC
17 |DBCO|TTCAGTCTATGCGACAGTCTGGCAATCTGACAT
20 |DBCO|TTTTTTTTGCAGTACTATGCAGTAACCTCAGCGCCCTTATTCCC
22 |DBCO|TTAAGTTAGAATTGCGTTGCGCTCCTGCAGCAGGTACATGTTAG
25 |DBCO|TTCTTGAGAGCACACCCGCCGCGCTTAATGCCAGCGGTAAGATC
27 |DBCO|TTCGAGTGGGGATGCTGAAGATCAGTGTGCTGCC
29 |DBCO|TTCTTACGGATACTCACCAGTCACAGTGATGAGCACTTTTAATTTTTAGTTCTGCTATTGA
31 |DBCO|TTTACACTATCGCCTTGATCGTTGGGAACCGCTCGGTCGCCGCA
33 |DBCO|TTTATCCCGTAGGGAGCCCCCGATTTAGAGCTGTGGCGCGGTAT
35 |DBCO|TTAGCTCATTTTTGGGGTCGAGGTGCCGTAATTTTTGTTAAATC
37 |DBCO|TTCGACGGGGCAGACCCCAATCATATGTACCGTAGTTATCTACA
40 |DBCO|TTTTAACGTGTGATAATCTCATGACCAAATCCCT
43 |DBCO|TTCAGACCAACACTGATTAAGCATTGTATCTGTG
45 |DBCO|TTTCCTACTTTTTTTTAAAAGAAAAGAGGTAACGGGTA
46 |DBCO|TTGAAACCTTTTTTTGCACTCTGAAAACAAAATCTGTTTTTGCGCCCGGGTTCGCCCGCATTTTTTT TTG
47 |DBCO|TTCTCTATGTGGCTCAACAACAGAGAATAAGCGTTGTTTTGATA
48 |DBCO|TTAAGCTTTATCTGTCTATACAAACTATTCGACGA
51 |DBCO|TTAAGAGTATTTAAACGAATAGCCTCTCCACATATTGAAAAAGG
52 |DBCO|TTAGACAATACATTCAAATATGTATCACCCAGAA
55 |DBCO|TTGAACCCCTGCACTTTTCGGGGAAAGGCCAACT
58 |DBCO|TTTTTTGCACGAGGACCGAAGGAGCTGCGCAAAC
61 |DBCO|TTTGGGGCCAAACGACGAGCGTGACACCACGATCATTGCAGCAC
63 |DBCO|TTCTGGAGCCCTGGCTGGTTTATTGCTAGACAGA
64 |DBCO|TTGATAATAACCTATTTTTATAGGTTCTAGCTTC

**Modified oligos (d30_60mer)**

2 |DBCO|TTCTATGGCTTTTTGAAATTCTTAATCGGTATCCATTTTTAAGATGAGAAGTGGCGTCAAAGG
3 |DBCO|TTAAAATAGATATGTGTAAAAGGAACGTAGCGGT
4 |DBCO|TTCGGGGAATTTTTAGCCGGCGAAGAGCGGGCGCTTTTTTAGGGCGCTGGCAAGTTGCTATGC
5 |DBCO|TTAGCGAAAGCGTGGCGAGAAAGGAACGTGAACC

7 |DBCO|TTTATAAATCTTTTTAAAAGAATAGGAACAAGAGTCTTTTTCACTATTAAACCGTC
8 |DBCO|TTGCGAAAAAGAACGTGGACTCCAACTAACGCAA
9 |DBCO|TTCCAGTTTGACCGAGATAGGGTTGAGGTCAGTT
10 |DBCO|TTCGAAGGTTTTTATTAGTGACGGACATCTAGTGTTTTTATAACCATGTTACTTGTGTTGTT
11 |DBCO|TTGTCTGGCGTAACTCCCGGGTTGGTATAATTGT
12 |DBCO|TTGCAAGATATTTTTTTAATACCGTGTATCTTGCACAC
13 |DBCO|TTGGAAGCCGCCTGTATCGGCCTTCATGGTCGTTAATGACCCTTTTTAAGAAAGCATCGTAG
14 |DBCO|TTGTATCTTCCCACAAGTAAATATCTTCACGAGGTAGCAAATAC
17 |DBCO|TTCAGTCTATGCGACAGTCTGGCAATCTGACAT
18 |DBCO|TTCCGAGGGTTTTTTTCCCAGAGCTATATTACTAGTCA
19 |DBCO|TTTTACATGTTTTTCGATGTGACCGAGTATTCAACTTTTTATTTCCGTGTGATAG
20 |DBCO|TTTTTTTTGCAGTACTATGCAGTAACCTCAGCGCCCTTATTCCC
21 |DBCO|TTTACAGGTTTTTGCGCACATTAAATTTTGACTCTTTTTGGGTGTAATCGGCA
22 |DBCO|TTAAGTTAGAATTGCGTTGCGCTCCTGCAGCAGGTACATGTTAG
23 |DBCO|TTAATTATTGATTTTTGCTGGTTACTATAC
24 |DBCO|TTACGCTGGTTTTTTGAAAGTAAAATTACATCGAACTTTTTTGGATCTCAAGCCGC
25 |DBCO|TTCTTGAGAGCACACCCGCCGCGCTTAATGCCAGCGGTAAGATC
26 |DBCO|TTCACGCTGCTTTTTGCGTAACCACTTTTCGCCCCGTTTTTAAGAACGTTTTCCAAAAAAGCAT
27 |DBCO|TTCGAGTGGGGATGCTGAAGATCAGTGTGCTGCC
28 |DBCO|TTGGTTGAGTGGCATGACAGTTTTTTAAGAGAATTATGCATGGGTGCA
29 |DBCO|TTCTTACGGATACTCACCAGTCACAGTGATGAGCACTTTTAATTTTTAGTTCTGCTATTGA
31 |DBCO|TTTACACTATCGCCTTGATCGTTGGGAACCGCTCGGTCGCCGCA
32 |DBCO|TTTAAATCTTTTTGGAACCCTAAATTGACGCCGGTTTTTGCAAGAGCAAGAGC
33 |DBCO|TTTATCCCGTAGGGAGCCCCCGATTTAGAGCTGTGGCGCGGTAT
35 |DBCO|TTAGCTCATTTTTGGGGTCGAGGTGCCGTAATTTTTGTTAAATC
36 |DBCO|TTATCACCCTTTTTTAATCAAGTTTTTTTAACCAATTTTTTAGGCCGAAATCGGCAAAAATCCC
37 |DBCO|TTCGACGGGGCAGACCCCAATCATATGTACCGTAGTTATCTACA
38 |DBCO|TTAGCGTAGTCAGGCAACTTTTTTATGGATGAACGAAATGATAAAT
39 |DBCO|TTTAGGTGATTTTTAGATCCTTTTAGTTTTCGTTCTTTTTCACTG
40 |DBCO|TTTTAACGTGTGATAATCTCATGACCAAATCCCT
41 |DBCO|TTTCGCTGAGTTTTTATAGGTGCCTGTTTACTCATATTTTTTATACTTTAGGATC
42 |DBCO|TTATCGGACCTTTTTACGTGTATTCATTCAATTTTTAATTTAAAAGATTGATTTAAAACTTCTTGTGCAA
43 |DBCO|TTCAGACCAACACTGATTAAGCATTGTATCTGTG
44 |DBCO|TTCGCAGCACATAAGTGTTGTTTTTTAGAAAGTCAAATCCGTAACTGT
45 |DBCO|TTTCCTACTTTTTTTTAAAAGAAAAGAGGTAACGGGTA
46 |DBCO|TTGAAACCTTTTTTTGCACTCTGAAAACAAAATCTGTTTTTGCGCCCGGGTTCGCCCGCATTTTTTTTTG
47 |DBCO|TTCTCTATGTGGCTCAACAACAGAGAATAAGCGTTGTTTTGATA
48 |DBCO|TTAAGCTTTATCTGTCTATACAAACTATTCGACGA
49 |DBCO|TTAGTTTGTTTTTAGGGGACGACGACGAAAGAGACA

51 |DBCO|TTAAGAGTATTTAAACGAATAGCCTCTCCACATATTGAAAAAGG
52 |DBCO|TTAGACAATACATTCAAATATGTATCACCCAGAA
53 |DBCO|TTTTTTGCCTTTTTTTCCTGTTTTTGCTCCGCTCATG
54 |DBCO|TTAAGGGCCTTTTTTCGTGATACGTGGTTTCTTAGTTTTTACGTCAGGTGATTTG
55 |DBCO|TTGAACCCCTGCACTTTTCGGGGAAAGGCCAACT
56 |DBCO|TTATAACCATTTTTTGAGTGATAACACTGCTGTGCGCG
57 |DBCO|TTTACTTCTGTTTTTACAACGATCGAACAT
58 |DBCO|TTTTTTGCACGAGGACCGAAGGAGCTGCGCAAAC
59 |DBCO|TTTGTAGCTTTTTAATGGCAACAACGTTAACCGCTT
60 |DBCO|TTCGCTCGTTTTTGCCCTTCCGGGGTGAGCGTGGTTTTTGTCTCGCGGTATGCC
61 |DBCO|TTTGGGGCCAAACGACGAGCGTGACACCACGATCATTGCAGCAC
62 |DBCO|TTTGAATGATTTTTAGCCATACCAGATGG
63 |DBCO|TTCTGGAGCCCTGGCTGGTTTATTGCTAGACAGA
64 |DBCO|TTGATAATAACCTATTTTTATAGGTTCTAGCTTC
65 |DBCO|TTTATTAACTTTTTTGGCGAACTACTTACTAATGTCAT
66 |DBCO|TTCCGGCAACTTTTTAATTAATAGATTATG
67 |DBCO|TTAAGAACCATTTTTGGGAAAACCAGGTACAAAGTTGCAGGACCACCTGGATGGAGGCGGATGAGCGAAC

**Supplementary Table 3. Primers for DNase I genotyping**

| | |
|---|---|
| DNase 1 Common FWD: | CTC ATC ATC TCT GTA GAC ATC |
| DNase 1 WT Rev: | TCA CCT GGC CCT CCA AAC |
| DNase 1 KO Rev: | GTC TGT CCT AGC TTC CTC ACT G |

**Supplementary Table 4. Primers used for BCR sequencing**

Heavy chain, Forward primers HUMAN

| | Sequence |
|---|---|
| P7_VH-1_v1 | GTCTCGTGGGCTCGGAGATGTGTATAAGAGACAGCAGGTSCAGCTGGTRCAGTC |
| P7_VH-2_v1 | GTCTCGTGGGCTCGGAGATGTGTATAAGAGACAGCAGRTCACCTTGAAGGAGTC |
| P7_VH-3_v1 | GTCTCGTGGGCTCGGAGATGTGTATAAGAGACAGSAGGTGCAGCTGGTGGAGTC |
| P7_VH-4_v1 | GTCTCGTGGGCTCGGAGATGTGTATAAGAGACAGCAGGTGCAGCTGCAGGAGTC |
| P7_VH-5_v1 | GTCTCGTGGGCTCGGAGATGTGTATAAGAGACAGGARGTGCAGCTGGTGCAGTC |
| P7_VH-6_v1 | GTCTCGTGGGCTCGGAGATGTGTATAAGAGACAGCAGGTACAGCTGCAGCAGTC |

Heavy chain, IgG Reverse primer HUMAN

| | |
|---|---|
| P5_C_gamma_v1_hs | TCGTCGGCAGCGTCAGATGTGTATAAGAGACAGGACSGATGGGCCCTTGGTGGA |

Heavy chain, IgM Reverse primer HUMAN

| | |
|---|---|
| P5_C_mu_v1_hs | TCGTCGGCAGCGTCAGATGTGTATAAGAGACAGGAAAAGGGTTGGGGCGGATGC |

Heavy chain, V gene Forward primers MOUSE

| | |
|---|---|
| 5' Igh_V_Ms | GTCTCGTGGGCTCGGAGATGTGTATAAGAGACAGGGGAATTCGAGGTGCAGCTGCAGGAGTCTGG |

Heavy chain, Reverse primers MOUSE

| | |
|---|---|
| 3' Igh_G1_C_Ms | TCGTCGGCAGCGTCAGATGTGTATAAGAGACAGGCTCAGGGAAATAGCCCTTGAC |
| 3' Igh_G2c_C_Ms | TCGTCGGCAGCGTCAGATGTGTATAAGAGACAGGCTCAGGGAAATAACCCTTGAC |
| 3' Igh_G2b_C_Ms | TCGTCGGCAGCGTCAGATGTGTATAAGAGACAGACTCAGGGAAGTAGCCCTTGAC |
| 3' Igh_G3_C_Ms | TCGTCGGCAGCGTCAGATGTGTATAAGAGACAGGCTCAGGGAAGTAGCCTTTGAC |

Light chain Lambda, Forward primers MOUSE

| | |
|---|---|
| 5' Igl_V1/2_Ms | GTCTCGTGGGCTCGGAGATGTGTATAAGAGACAGCAGGCTGTTGTGACTCAG |
| 5' Igl_V3_Ms | GTCTCGTGGGCTCGGAGATGTGTATAAGAGACAGCAACTTGTGCTCACTCAG |

Light chain Lambda, Reverse primer MOUSE

| | |
|---|---|
| 3' Igl_C_Ms | TCGTCGGCAGCGTCAGATGTGTATAAGAGACAGCTCYTCAGRGGAAGGTGGRAACA |

Light chain Kappa, Forward primer MOUSE

| | |
|---|---|
| 5' Igk_V_Ms | GTCTCGTGGGCTCGGAGATGTGTATAAGAGACAGGAYATTGTGMTSACMCARWCTMCA |

Light chain Kappa, Reverse primer MOUSE

| | |
|---|---|
| 3' Igk_C_Ms | TCGTCGGCAGCGTCAGATGTGTATAAGAGACAGGATGGTGGGAAGATGGATACAGTT |

Heavy chain, IgM Reverse primer MOUSE

| | |
|---|---|
| 3' Igh_M_C_Ms | TCGTCGGCAGCGTCAGATGTGTATAAGAGACAGAGGGGGAAGACATTTGGGAAGGAC |
